# Targeting Chromatin Effector Pygo2 to Enhance Immunotherapy in Prostate Cancer

**DOI:** 10.1101/2022.06.16.496505

**Authors:** Yini Zhu, Yun Zhao, Jiling Wen, Sheng Liu, Tianhe Huang, Ishita Hatial, Xiaoxia Peng, Hawraa Al Janabi, Gang Huang, Jackson Mittlesteadt, Michael Cheng, Atul Bhardwaj, Brandon L. Ashfeld, Kenneth R. Kao, Dean Y. Maeda, Xing Dai, Olaf Wiest, Brian S.J. Blagg, Xuemin Lu, Liang Cheng, Jun Wan, Xin Lu

## Abstract

The noninflamed microenvironment in prostate cancer represents a barrier to immunotherapy. Genetic alterations underlying cancer cell-intrinsic oncogenic signaling have emerging roles in shaping the immune landscape. Recently, we identified Pygopus 2 (*PYGO2*) as the driver oncogene for the amplicon at 1q21.3. Here, using transgenic mouse models of metastatic prostate adenocarcinoma, we found that *Pygo2* deletion decelerated tumor progression, diminished metastases, and extended survival. Pygo2 loss augmented the infiltration of cytotoxic T lymphocytes (CTLs) and sensitized tumor cells to T cell killing. Mechanistically, Pygo2 orchestrated a p53/Sp1/Kit/Ido1 signaling network to foster a microenvironment hostile to CTLs. Genetic or pharmacological inhibition of Pygo2 enhanced the anti-tumor efficacy of immunotherapies using immune checkpoint blockade, adoptive cell transfer, or myeloid-derived suppressor cell inhibitors. In human prostate cancer samples, Pygo2 expression was inversely correlated with CD8^+^ T cells. Our results highlight a promising path to improving immunotherapy with targeted therapy for lethal prostate cancer.

## INTRODUCTION

Tumor resistance to immunotherapy remains a significant challenge ^2^. Among cancer types refractory to immune check blockade (ICB), advanced prostate cancer (PCa) exhibits overwhelming *de novo* resistance to anti-CTLA4 or anti-PD1 therapies ^3-6^. A tumor microenvironment (TME) poorly infiltrated by immune cells or infiltrated by broad immunocytes but void of cytotoxic T lymphocytes (CTLs) in the tumor core is considered immunologically cold. To combat resistance to immunotherapy, therapeutic efforts using targeted agents to convert cold to hot TME are promising approaches. PCa is generally considered immunologically cold, where T lymphocytes are primarily located in the adjacent normal structures ^8^, or the tumor stroma, but rarely the invasive epithelium ^9^. A cold TME can be shaped by genetic alterations and oncogenic pathways intrinsic to cancer cells ^12,13^. Advanced PCa is characterized by rampant chromosomal instability and copy number alterations, including deletions and amplifications ^14^. Using genetically engineered mouse (GEM) models, studies have revealed the differential effects of the loss of distinct tumor suppressor genes (*Pten, Zbtb7a, p53, Pml*, and *Smad4*) on the infiltration frequency and activity of various myeloid populations ^15,16^. For example, we reported that in the *PB-Cre4*^*+*^ *Pten*^*L/L*^ *Smad4*^*L/L*^ model, *Smad4* loss caused Yap1-mediated upregulation of Cxcl5 in tumor cells, which in turn recruited Cxcr2^+^ polymorphonuclear myeloid-derived suppressor cells (PMN-MDSCs) to antagonize anti-tumor T-cell immunity ^15^. The contribution of oncogenes amplified in the PCa genome to the immunosuppressive TME is poorly understood. This is an important question because of the potential therapeutic opportunities associated with targeting amplified oncogenes.

Pygopus family PHD finger 2 (*PYGO2*) was recently identified through an *in vivo* functional screen as the driver oncogene for the amplicon at 1q21.3 in human PCa ^17^. Copy number gain or amplification of *PYGO2* was detected in over 50% of primary and metastatic castration-resistant PCa cases and was associated with a higher Gleason score, shorter disease-free survival, and shorter biochemical recurrence ^17^. At the protein level, while PYGO2 expression was not detectable in the normal prostate, high PYGO2 expression was correlated with a higher Gleason score, biochemical recurrence, and metastasis to lymph nodes and bone ^17,18^. Overexpression of PYGO2 has been documented in ovarian ^19^, breast ^20^, cervical ^21^, hepatic ^22^, lung ^23^, intestinal ^24^, and brain cancers^25^. Therefore, understanding and targeting PYGO2 may have translational significance in various cancer types. As a chromatin effector, PYGO2 anchors to chromatin through interactions between its plant homeodomain (PHD) and H3K4me2/3, histone modifications that mark active transcription ^26^. PYGO2 in turn recruits histone acetyltransferases or histone methyltransferases to promote histone modifications and augment Wnt/β-catenin– mediated transcriptional activation ^27^. As an emerging epigenetic switch, PYGO2 regulates stem cell self-renewal, somatic cell division, and hormone-induced gene expression through Wnt-dependent ^28,29^ and Wnt-independent pathways ^30,31^. To date, studies on PYGO2 have focused on its cell-autonomous functions. In the current study, we used GEM models of PCa to discover the cell non-autonomous role of PYGO2 in shaping the immunosuppressive TME of PCa, particularly the poor infiltration and activity of effector T cells. Importantly, genetic ablation or pharmacological inhibition of PYGO2 sensitizes PCa to ICB, adoptive T-cell therapy (ACT), and PMN-MDSC inhibition, illuminating a clinical path hypothesis for combining PYGO2-targeted therapy and immunotherapy in the treatment of lethal PCa.

## RESULTS

### Pygo2 promotes PCa progression and metastasis in GEM and syngeneic models

The function of Pygo2 during spontaneous PCa development was not defined. To investigate this, we crossed Pygo2 loxP allele ^32^ with the metastatic prostate adenocarcinoma GEM model, *PB-Cre4*^*+*^ *Pten*^*L/L*^ *Smad4*^*L/L*^ (pDKO) ^15,33^, and generated *PB-Cre4*^*+*^ *Pten*^*L/L*^ *Smad4*^*L/L*^ *Pygo2*^*L/L*^ (pTKO) mice (**Fig. 1a**). Prostate-specific Pygo2 loss was evident in pTKO mice (**Fig. 1b**). pTKO mice exhibited decelerated tumor growth by approximately two months, as detected by magnetic resonance imaging (MRI) (**Fig. 1c**). The median survival of pTKO mice was extended by ten weeks compared with pDKO mice (**Fig. 1d**). To compare histological features at equivalent tumor sizes, we harvested tumors from 12-week pDKO mice and 18-week pTKO mice (n=5, **Fig. 1e**). Immunohistochemistry (IHC) showed lower proliferation and stronger apoptosis in pTKO tumors than pDKO tumors (**Fig. 1f**).

**Fig. 1.**
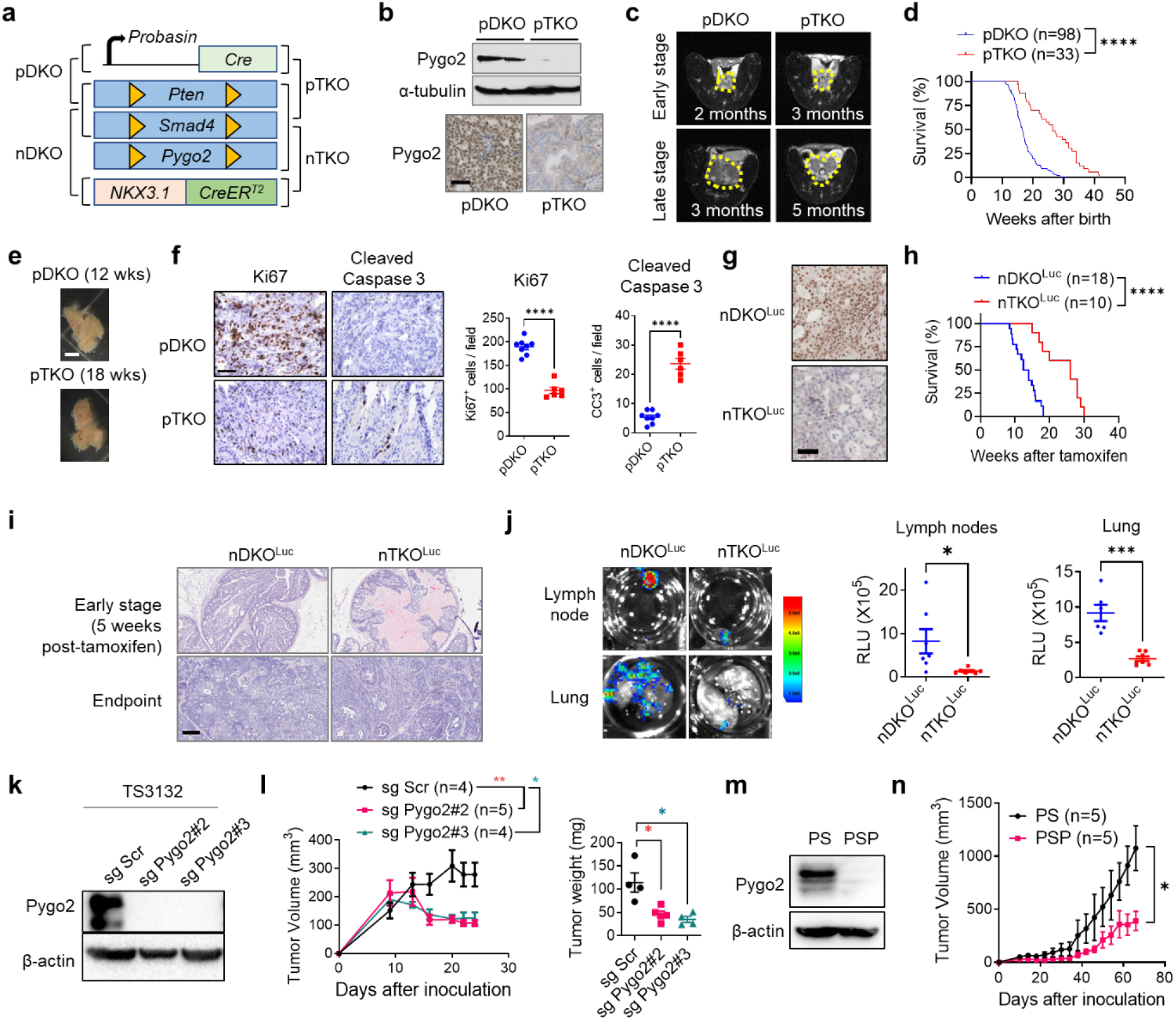
Pygo2 promotes PCa progression and metastasis in GEM and syngeneic models. **(a)** The diagram for the generation of conditional knockout models including pDKO, pTKO, nDKO and nTKO. **(b)** Pygo2 protein expression in pDKO and pTKO tumors evaluated by western blot and IHC. Scale bar, 50µm. **(c)** Representative MRI images of pDKO and pTKO tumors (yellow contour) at early and late stages. **(d)** Kaplan–Meier curves of pDKO (n=98) and pTKO (n=33) mice. **(e)** Representative photographs of tumors from 12-week pDKO and 18-week pTKO mice, respectively. **(f)** Ki67 and cleaved caspase-3 IHC staining and quantification for pDKO and pTKO tumors (n = 6-8). Scale bar, 50µm. **(g)** IHC for Pygo2 in nDKO^Luc^ and nTKO^Luc^ tumors. Scale bar 50µm. **(h)** Kaplan–Meier curves of nDKO^Luc^ (n=18) and nTKO^Luc^ (n=10) mice after tamoxifen induction. **(i)** H&E staining for nDKO^Luc^ and nTKO^Luc^ at early stage and endpoint. Scale bar, 200µm. **(j)** Bioluminescence images (left) and quantification (right) of metastases at draining lymph nodes and lungs from nDKO^Luc^ (n=6) and nTKO^Luc^ (n=7) mice at the endpoint. **(k)** Pygo2 knockout in TS3132 with two different sgRNA, validated by western blot. **(l)** Subcutaneous tumor volume (left) and endpoint weight (right) for TS3132 sublines in nude mice (n = 4-5). **(m)** Western blot validating Pygo2 expression in PS and PSP cell lines. **(n)** Syngeneic tumor growth curves for PS (n=5) and PSP (n=5) in C57BL/6 mice. In (d)(h), ****P<0.0001, log-rank test. In (f)(j)(l)(n), error bars represent SEM; *P<0.05, **P<0.01, ***P<0.001, ****P<0.0001, Student’s t-test.

The tamoxifen-inducible *Nkx3*.*1*^*CreERT2*^ allele enables temporal control of gene deletion in prostatic epithelial cells ^34^. We generated *Nkx3*.*1*^*CreERT2/+*^ *Pten*^*L/L*^ *Smad4*^*L/L*^ *Rosa26-LSL-Luc*^*L/L*^ (nDKO^Luc^) and *Nkx3*.*1*^*CreERT2/+*^ *Pten*^*L/L*^ *Smad4*^*L/L*^ *Pygo2*^*L/L*^ *Rosa26-LSL-Luc*^*L/L*^ (nTKO^Luc^) mice and confirmed tamoxifen-induced Pygo2 expression loss (**Fig. 1g**). Consistent with the *PB-Cre4*-based models, nTKO^Luc^ mice survived 12.8 weeks (median survival) longer than nDKO^Luc^ mice (**Fig. 1h**), consistent with slower tumor growth in the former (**Fig. 1i**). Metastases to draining lymph nodes and lungs were also attenuated in nTKO^Luc^ compared to nDKO^Luc^ mice (**Fig. 1j**).

To facilitate mechanistic studies, we used CRISPR/cas9 to knockout *Pygo2* in the previously reported murine PCa cell line TS3132, which was derived from the pDKO model and formed tumors when implanted in immune-deficient mice ^33^ (**Fig. 1k**). Pygo2 knockout significantly decreased colony formation (**Extended Data Fig. 1a-b**) and attenuated subcutaneous tumor growth (**Fig. 1l**). TS3132 sublines were labeled with a tk-GFP-luciferase reporter ^35^ and injected intracardially into nude mice. Pygo2 knockout largely depleted the metastatic ability in bone, lungs, liver, and brain (**Extended Data Fig. 1c-e**). TS3132 was derived from mice with a mixed genetic background; therefore, it cannot grow in immune-competent mice. To facilitate the study of Pygo2 function in tumor immune regulation, we established *Pten/Smad4* (PS) and *Pten/Smad4/Pygo2* (PSP) cell lines from pDKO and pTKO tumors (**Fig. 1m**). When injected subcutaneously into C57BL/6 males, PS tumors grew significantly faster than PSP tumors (**Fig. 1n**). These newly established GEM and syngeneic models reinforced the PCa-promoting function of Pygo2 and prompted us to investigate the previously uncharacterized mechanisms underlying this function.

### Pygo2 restricts CTL infiltration and attenuates CTL killing of PCa cells

To assess the potential role of Pygo2 in modulating the TME, we used mass cytometry (CyTOF) to quantify the primary immune cell populations in pDKO and pTKO tumors. CD8^+^ T cells were significantly increased in pTKO tumors (**Fig. 2a, Extended Data Fig. 2a**). The higher infiltration of total T cells and CD8^+^ T cell subsets in pTKO than pDKO tumors was validated by IHC (**Fig. 2b**). When nDKO^Luc^ and nTKO^Luc^ tumors were compared using flow cytometry, Pygo2-deficient tumors had increased CD8^+^ T cells, increased CD4^+^ T cells, and decreased T_reg_ fraction in CD4^+^ T cells (**Fig. 2c**). Similar differences were observed in pDKO and pTKO tumors (**Extended Data Fig. 2b**). We previously reported that the most prominent immune population in pDKO tumors was PMN-MDSCs ^15^. PMN-MDSCs are tumor-infiltrating neutrophils with immunosuppressive activity ^36^. We confirmed similar levels of PMN-MDSCs and macrophages between pDKO and pTKO tumors and between nDKO^Luc^ and nTKO^Luc^ tumors (**Extended Data Fig. 2c**), suggesting that the infiltration difference of T cell subsets by Pygo2 loss was unlikely to be explained by changes in PMN-MDSCs. Consistent with spontaneous tumors, syngeneic PSP tumors harbored more CD8^+^ and CD4^+^ T cells and a higher CD8^+^/T_reg_ ratio (**Fig. 2d**). Critically, this pattern was reversed by restoring Pygo2 expression in PSP cells (**Fig. 2d**), supporting a causal role of Pygo2 in PCa cells in dictating T cell phenotypes.

**Fig. 2.**
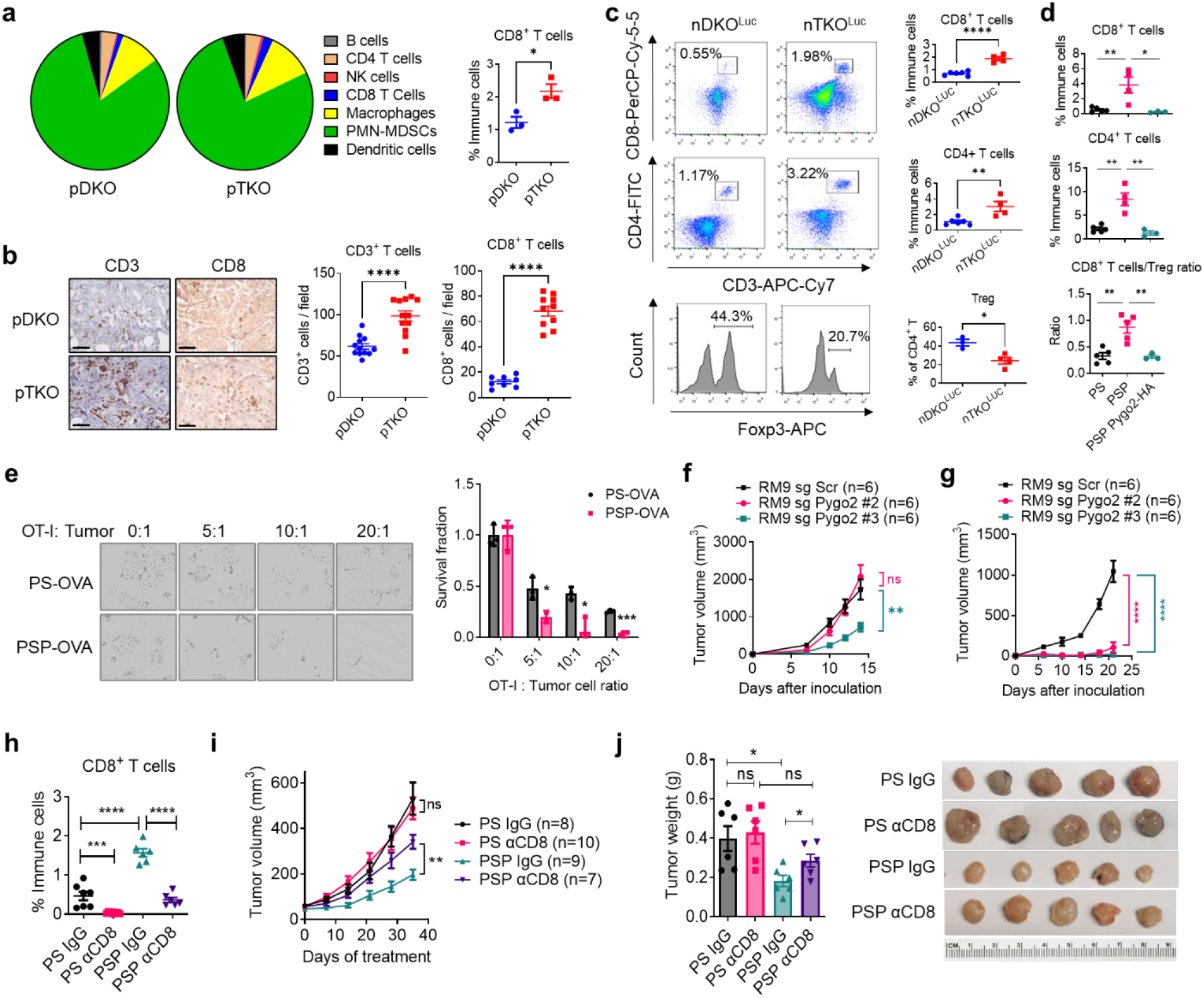
Pygo2 inhibits CTL infiltration and attenuates CTL killing of PCa cells. **(a)** Major immune population fractions in pDKO and pTKO tumors (n=3/genotype) quantified by CyTOF (left) with CD8^+^ T cells showing significant difference (right). **(b)** IHC staining and quantification for CD8 and CD3 in pDKO and pTKO tumors (n=12). **(c)** Flow cytometry quantification of CD8^+^, CD4^+^ and T_reg_ percentages for nDKO^Luc^ and nTKO^Luc^ tumors (n = 3-6). **(d)** Flow cytometry quantification of CD8^+^ T, CD4^+^ T, and CD8^+^ effector/T_reg_ ratio for syngeneic subcutaneous tumors formed by PS, PSP, or PSP-Pygo2-HA cells (n = 3-5). **(e)** T cell cytotoxicity assay to compare the killing of PS-OVA and PSP-OVA cells by antigen-stimulated OT-I T-cells at different E:T ratios. Viable cancer cells were detected by resazurin (n=3). **(f-g)** Tumor growth curves of RM9 sgScr and sgPygo2 sublines in nude (f) or C57BL/6 mice (g) (n=6). **(h-i)** Flow cytometry quantification of intratumoral CD8^+^ T cells and tumor growth curves for syngeneic tumors formed by PS or PSP and treated with isotype IgG or anti-CD8 antibody (n = 6-10). (**j**) Endpoint tumor weight (n=6) and representative tumor photographs. In all panels, error bars represent SEM; ns, not significant, *P<0.05, **P<0.01, ***P<0.001, ****P<0.0001, Student’s t-test.

Based on the negative impact of Pygo2 on effector T cells in the TME, we postulated that Pygo2 in PCa cells might drive resistance to T-cell killing. We stably expressed chicken ovalbumin (OVA) in PS and PSP cell lines and co-cultured these sublines with OVA-specific TCR-transgenic CD8^+^ T cells (OT-I). PSP-OVA cells were more sensitive to OT-I T-cell killing than PS-OVA cells (**Fig. 2e**). To rule out that the function of Pygo2 in regulating tumor-T cell interactions is specific to the Pten/Smad4 model, we silenced Pygo2 expression with CRISPR/cas9 in the murine PCa cell line RM9 (transformed by *ras* and *myc* ^37^) (**Extended Data Fig. 2d**). Applying the OVA/OT-I system to RM9 sublines corroborated that Pygo2 loss sensitized PCa cells to T-cell killing (**Extended Data Fig. 2e**). While Pygo2 knockout in RM9 affected tumor growth modestly in nude mice (**Fig. 2f**), it augmented T cell infiltration and dramatically decreased tumor formation in C67BL/6 mice (**Fig. 2g, Extended Data Fig. 2f**). We depleted CD8^+^ T cells in C57BL/6 mice bearing PS and PSP tumors using an anti-CD8 neutralizing antibody (**Fig. 2h, Extended Data Fig. 2g**). CTL ablation had little impact on PS tumor growth but significantly restored the PSP tumor growth (**Fig. 2i-j**). Taken data from different models, we conclude that Pygo2 expression in PCa cells elicits cell non-autonomous activity to restrict effector T cell infiltration and cytotoxicity.

### Pygo2 promotes PCa progression through Kit upregulation in a Wnt-independent manner

Despite functional and clinical validation of Pygo2 in driving PCa progression ^17,18^, the mechanism underlying Pygo2 function in PCa remains poorly understood. To identify Pygo2-regulated genes, we dissociated pDKO and pTKO tumors with *B. Licheniformis* protease at 4°C to minimize artificial changes in gene expression patterns ^38^, followed by epithelial cell purification and microarray profiling (**Fig. 3a**). We identified 379 differentially expressed (DE) probes (p<0.05) between pDKO and pTKO tumor cells and validated several by qRT-PCR (**Extended Data Fig. 3a-c, Supplementary Table 1**). Gene set enrichment analysis (GSEA) with MSigDB hallmark gene sets showed that the p53 pathway and epithelial-mesenchymal transition (EMT) pathway were enriched in pDKO tumor cells, whereas immune-related pathways, such as interferon α response, interferon γ response, complement, and IL6-JAK-STAT3 signaling, were enriched in pTKO tumor cells (**Fig. 3b, Supplementary Table 2-3**). We reasoned that Pygo2 might exert its immunomodulatory activity through specific mediators. To find the mediator(s), we performed an upstream analysis based on DE genes with Ingenuity pathway analysis (IPA). A list of putative upstream regulators was identified, including the receptor tyrosine kinase Kit, which was downregulated in pTKO tumor cells (**Supplementary Table 4**). Among the genes downregulated in pTKO PCa cells, multiple were mapped as Kit downstream genes by IPA (**Fig. 3c**). We validated Pygo2-loss-induced *Kit* downregulation using sorted pDKO and pTKO PCa cells (**Fig. 3d**). At the protein level, Kit and some of the Kit-downstream signaling proteins were attenuated in pTKO tumors compared to pDKO tumors (**Fig. 3e**). Among them, Ido1 was reported to drive Kit-induced T-cell suppression in gastrointestinal stromal tumors (GIST) ^39^. Kit downregulation by Pygo2 knockout was evident in PCa cell lines (**Fig. 3f**). IHC confirmed higher Kit and Ido1 expression in pDKO than pTKO tumors (**Fig. 3g**).

**Fig. 3.**
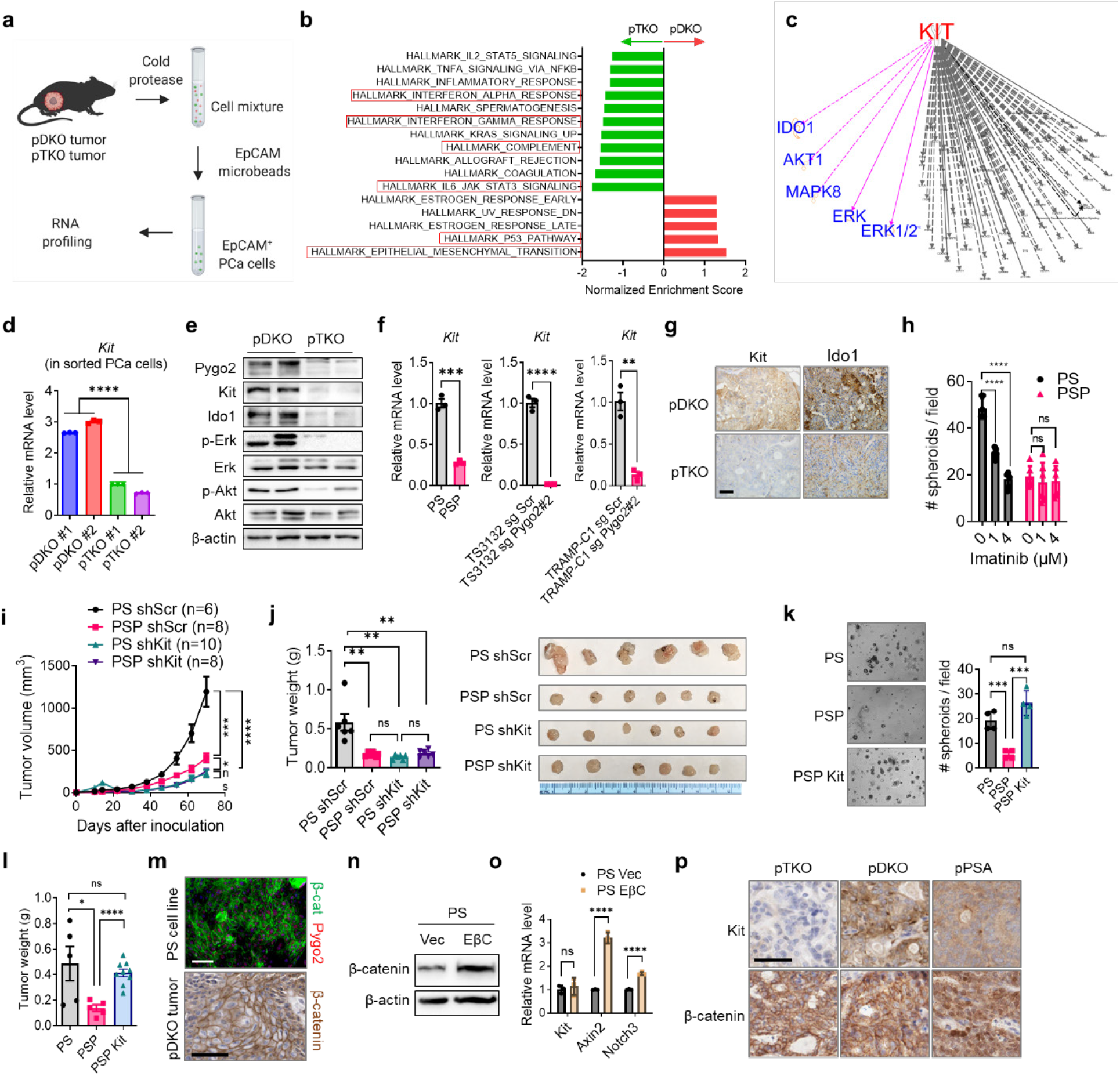
Pygo2 promotes PCa progression through Kit upregulation in a Wnt-independent manner. **(a)** The schematic of tumor cell purification with EpCAM mcirobeads from pDKO (n=3) and pTKO (n=2) tumors followed by transcriptomic profiling. **(b)** Top enriched MSigDB hallmark gene sets in pDKO (red) or pTKO (green) tumor cells. **(c)** KIT downstream targets drawn based on IPA knowledgebase. Targets highlighted in purple include a number of DE genes (Ido1, Akt1, Mapk8) based on the transcriptomic profiling. **(d)** qRT-PCR validation of *Kit* differential expression using purified tumor cells from pDKO (n=3) and pTKO tumors (n=3). **(e)** Western blot validation of Kit and selected downstream targets in pDKO and pTKO tumors. **(f)** qRT-PCR measurement of *Kit* expression in mouse PCa cell lines (n=3). **(g)** IHC for Kit and Ido1 in pDKO and pTKO tumors. Scale bar, 50μm. **(h)** Spheroid assay of PS and PSP cell lines treated with vehicle or imatinib (n=6). **(i)** Syngeneic tumor growth curves for PS and PSP sublines in C57BL/6 mice (n = 6-10). **(j)** Endpoint tumor weight and representative photographs of tumors formed by PS and PSP sublines (n=6). **(k)** Spheroid assay for PS, PSP and PSP-Kit (n=4). **(l)** Endpoint tumor weight for syngeneic tumors formed by PS (n=5), PSP (n=5) or PSP-Kit (n=9). **(m)** Representative β-catenin immunostaining results for PS cell line or pDKO tumor. Scale bar, 50μm. **(n)** Western blot detecting the ectopic overexpression of EβC in PS cells. **(o)** qRT-PCR detecting the effect of EβC on the expression of *Kit* and Wnt targets (*Axin2, Notch3*) in PS cells (n=3). **(p)** IHC detecting Kit and β-catenin in pTKO, pDKO and pPSA prostate tumors. Scale bar, 50μm. In (d)(f)(h)(i)(j)(k)(l)(o), error bars represent SEM; ns, not significant, *P<0.05, **P<0.01, ***P<0.001, ****P<0.0001, Student’s t-test.

Kit has oncogenic functions in GIST and acute myeloid leukemia. To determine whether Kit is essential for the pro-tumor function of Pygo2, we first confirmed that Kit inhibitor imatinib decreased PS spheroid formation and migration but had no effect on PSP (**Fig. 3h, Extended Data Fig. 3d**). Next, *Kit* shRNA knockdown decelerated the growth of PS tumors but generated no further tumor-retarding effect on the slow-growing PSP tumors in C57BL/6 mice (**Fig. 3i-j, Extended Data Fig.3e**), supporting that Kit is downstream of Pygo2. Rescuing Kit expression in PSP recovered spheroid formation (**Fig. 3k**) and *in vivo* tumorigenicity to the level of PS (**Fig. 3l**). These results established a causal relationship of the Pygo2-Kit axis in driving PCa.

To determine whether Pygo2 function in PCa and the Pygo2-Kit axis involve Wnt/β-catenin signaling, we first stained β-catenin in PS cells and pDKO tumors and found an almost exclusive cell membrane (but not nuclear) localization of β-catenin (**Fig. 3m**). This result is consistent with the lack of enrichment of the Wnt signaling pathway in a previous study that compared pDKO and *PB-Cre4*^*+*^ *Pten*^*L/L*^ tumors ^33^, indicating that Wnt signaling is not activated in pDKO. Moreover, a survey of the expression of Wnt target genes in sorted pDKO and pTKO PCa cells revealed no difference between the two genotypes (**Extended Data Fig. 3f**). To test the involvement of Wnt signaling more directly, we used several methods to modify the pathway and examine its effect on Kit expression. First, when canonical Wnt signaling was activated in PS cells by LiCl or Wnt3a conditioned medium treatment, Kit expression was not induced (**Extended Data Fig. 3g-h**). Next, overexpression of constitutively active β-catenin (EβC) in PS or RM9 cells (**Fig. 3n, Extended Data Fig. 3i**) enhanced the expression of classical Wnt/β-catenin targets but failed to affect Kit (**Fig. 3o, Extended Data Fig. 3j**). Lastly, we compared Kit levels between prostate tumors from pTKO, pDKO, and *PB-Cre4*^*+*^ *Pten*^*L/L*^ *Smad4*^*L/L*^ *Apc*^*L/L*^ (pPSA) mice. We recently reported the development of aggressive PCa and penile cancer in pPSA mice ^40^. We confirmed that despite the *Apc*-loss-induced nuclear localization of β-catenin in pPSA tumors, no further increase in Kit staining was observed in pPSA tumors compared with pDKO tumors (**Fig. 3p**). Our results argue against the role of Wnt/β-catenin in Pygo2-driven Kit expression and indicate that Pygo2 regulates Kit expression in a previously uncharacterized fashion.

### Pygo2 cooperates with p53 to upregulate the Sp1/Kit axis

Pygo2 depends on co-factors to regulate gene transcription. We first performed a connection analysis from Pygo2 to Kit using IPA (**Fig. 4a**). Only four factors were predicted to be potential mediators from Pygo2 to Kit, including CTNNB1 (β-catenin), UBC (ubiquitin C), PDGFRA (platelet-derived growth factor receptor A), and PDGFRB. We ruled out these factors because, first, β-catenin was not involved in Pygo2-Kit regulation (see above). Second, the expression of PDGFRA and PDGFRB showed no difference between pDKO and pTKO PCa cells (**Extended Data Fig. 4a**). Third, the connection between UBC to Pygo2 and Kit is based on protein-protein interactions that affect protein stability; thus, it is unlikely to account for Kit mRNA changes. Can Pygo2 regulate Kit by controlling the expression or activity of particular transcription factor (TF) that directly or indirectly regulate Kit? We used IPA to identify all TFs upstream of Kit (**Fig. 4a**). We filtered through them and focused on TP53 (i.e., p53) based on three findings. First, p53 pathway was among the top enriched pathways in pDKO tumor cells (**Fig. 3b**). Second, a TF enrichment analysis using the upregulated genes in pDKO PCa cells identified p53 as an enriched TF in pDKO (**Fig. 4b, Supplementary Table 5-6**). Third, Pygo2 was reported to induce the accumulation and acetylation of p53 in hair follicle early progenitor cells ^28^.

**Fig. 4.**
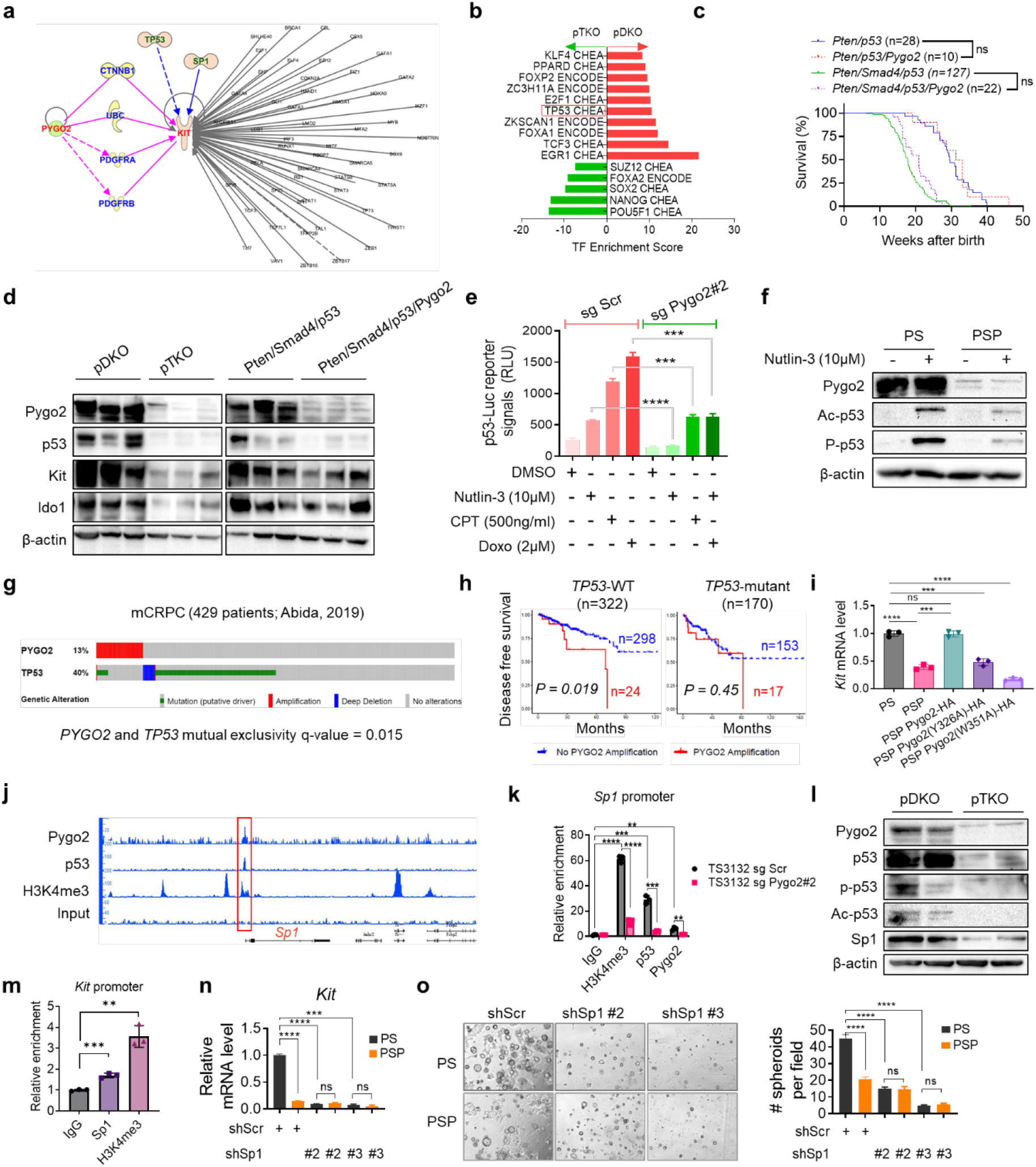
Pygo2 cooperates with p53 to upregulate the Sp1/Kit axis. **(a)** IPA result finding putative connectors (blue) between Pygo2 and Kit (red), and finding TFs (green, grey) regulating Kit. **(b)** TF enrichment analysis with Enrichr using ENCODE and ChEA databases for genes significantly upregulated in pDKO PCa cells compared with pTKO PCa cells. **(c)** Kaplan-Meier curves of four GEM cohorts, Pten/p53 (n=28), Pten/p53/Pygo2 (n=10), Pten/Smad4/p53 (n=127), and Pten/Smad4/p53/Pygo2 (n=22). ns, not significant, log-rank test. **(d)** Western blot measurement of Pygo2, p53, Kit and Ido1 in tumor lysates from pDKO, pTKO, Pten/Smad4/p53 and Pten/Smad4/p53/Pygo2 (n=3). **(e)** p53 reporter assay to detect the p53 activity in TS3132 sgSCr and Pygo2-knockout cells treated with DMSO (vehicle), nutlin-3, CPT or doxorubicin (n=3). **(f)** Western blot measurement of Pygo2, acetyl-p53 (Lys379), phospho-p53 (Ser15) in PS and PSP cell lines treated with vehicle or nutlin-3 for 24h. **(g)** Genomic alterations of *PYGO2* and *TP53* in the mCRPC database (n=429, ref Abida et al.), visualized by cBioPortal. **(h)** Disease free survival for patients stratified based on *TP53* mutation status followed by *PYGO2* amplification status. Dataset from TCGA (Firehose Legacy), n and P values denoted in the graphs, P values based on log-rank test. **(i)** qRT-PCR quantification of *Kit* expression in PS, PSP, and PSP sublines expressing HA-tagged WT or mutant murine Pygo2 (n=3). **(j)** IGB genomic views of the chromatin association of Pygo2, p53, and H3K4me3 at the genomic locus around *Sp1*, based on CUT&RUN-seq of PS cell line. Red rectangle highlights the peaks in the *Sp1* promoter region. **(k)** CUT&RUN-qPCR to quantify the association of Pygo2, p53, and H3K4me3 to the *Sp1* promoter region in TS3132 sublines. IgG was the negative control (n=3). **(l)** Western blot measurement of Pygo2, p53, acetyl-p53 (Lys379), phospho-p53 (Ser15), and Sp1 in pDKO and pTKO tumors. **(m)** CUT&RUN-qPCR to quantify the association of Sp1 and H3K4me3 to the *Kit* promoter region in TS3132. IgG was the negative control (n=3). **(n)** qRT-PCR quantification of *Kit* expression in PS and PSP sublines with Sp1 shRNA knockdown or shScr control (n=3). **(o)** Spheroid formation ability by PS and PSP sublines with Sp1 shRNA knockdown or shScr control (n = 7-10). In (e) (i) (k) (m) (n) (o), error bars represent SEM; ns, not significant, **P<0.01, ***P<0.001, ****P<0.0001, Student’s t-test.

Can p53 mediate Kit regulation by Pygo2 and even play an indispensable role in Pygo2 function in PCa? To test this functionally, we deleted *p53* with the *PB-Cre4* driver and compared the survival of two pairs of mouse cohorts: *PB-Cre4*^*+*^ *Pten*^*L/L*^ *p53*^*L/L*^ (Pten/p53) compared with *PB-Cre4*^*+*^ *Pten*^*L/L*^ *p53*^*L/L*^ *Pygo2*^*L/L*^ (Pten/p53/Pygo2), *PB-Cre4*^*+*^ *Pten*^*L/L*^ *Smad4*^*L/L*^ *p53*^*L/L*^ (Pten/Smad4/p53) compared with *PB-Cre4*^*+*^ *Pten*^*L/L*^ *Smad4*^*L/L*^ *Pygo2*^*L/L*^ *p53*^*L/L*^ (Pten/Smad4/p53/Pygo2). Strikingly, while *Pygo2* knockout extended survival substantially in the Pten/Smad4 genetic backdrop (**Fig. 1d**), *Pygo2* loss did not affect survival in the Pten/p53 or Pten/Smad4/p53 backdrops (**Fig. 4c**). At the expression level, p53 was higher in pDKO tumors than in pTKO tumors (**Fig. 4d**). Kit and its downstream protein, Ido1, were also higher in pDKO tumors than in pTKO tumors, yet these two proteins remained unaltered between Pten/Smad4/p53 and Pten/Smad4/p53/Pygo2 tumors (**Fig. 4d**) or derived cell lines (**Extended Data Fig. 4b**). To investigate how Pygo2 and p53 regulate Kit, we ruled out direct transcriptional regulation of Pygo2 on p53 (**Extended Data Fig. 4c-d**). Instead, evidence suggests that Pygo2 cooperates with p53 to regulate downstream genes: Pygo2 and p53 proteins co-localized in the nuclei, detected by proximity ligation assay (PLA) (**Extended Data Fig. 4e**); Pygo2 and p53 interaction was observed using co-immunoprecipitation (co-IP) followed by western blotting (**Extended Data Fig. 4f**).

Post-translational modifications of p53 regulate p53 stability and activity. Pygo2 recruits histone acetyltransferases (e.g., CBP/p300 and the STAGA complex) to modulate the activity of transcriptional co-factors ^27,41^. Acetylation of p53 by CBP/p300 directly affects its transcriptional activity ^42^. Using co-IP in PS cells, we confirmed the interaction between Pygo2, CBP/p300 and p53 (**Extended Data Fig. 4f**). A p53 reporter assay using TS3132 and Pygo2-knockout subline indicated that Pygo2 deletion dampened p53 activity stimulated by nutlin-3, camptothecin (CPT), or doxorubicin (**Fig. 4e**). Consistently, p53 acetylation and phosphorylation, indicative of p53 activity, were more pronounced in PS than PSP upon nutlin-3 (**Fig. 4f**) or CPT treatments (**Extended Data Fig. 4g**). Hyperactivated p53 induces cell cycle arrest and apoptosis. However, Pygo2 did not seem to participate in this aspect of p53 function, as PS and PSP cells showed similar cell cycle profiles upon stress from nutlin-3, CPT, or doxorubicin (**Extended Data Fig. 4h**). To examine the Pygo2/p53 interaction in clinical samples, we surveyed the genetic status of *PYGO2* and *TP53* in the SU2C/PCF human mCRPC dataset ^43^. *PYGO2* amplifications (13%) were mutually exclusive with *TP53* alterations (40%, mainly mutations and deep deletions) (**Fig. 4g**). Furthermore, in the PCa TCGA cohort ^44^, if patients were stratified into *TP53*-wild type (WT) and *TP53*-mutant groups, *PYGO2* amplification correlated with worse disease-free survival only in the *TP53*-WT group (**Fig. 4h**). Therefore, mouse and human genetic evidence support a critical role of p53 in Pygo2 function in PCa.

Next, we investigated how the Pygo2/p53 interaction regulates Kit expression. Pygo2 binding to histone H3K4me2/3 was crucial for Kit regulation, because Kit expression in PSP cells was rescued by ectopic expression of WT Pygo2, but not Pygo2 mutants (Y326A, W351A) deficient in H3K4me2/3 binding ^30,45^ (**Fig. 4I, Extended Data Fig. 4i**). The global association of Pygo2, p53, and H3K4me3 with chromatin was assessed using the CUT & RUN assay in PS cells. Neither Pygo2 nor p53 directly bound to *Kit* promoter region (**Extended Data Fig. 4j-k**), suggesting the existence of an intermediate regulator. A co-localized binding peak of Pygo2, p53, and H3K4me3 was found near the promoter region of *Sp1* (**Fig. 4j**), which was validated by qPCR using TS3132 sublines (**Fig. 4k**). Sp1 is often associated with a poor prognosis and regulates transcription in a context-dependent manner ^46^. Moreover, Sp1 binds to *Kit* promoter region to mediate transcription in hematopoietic cells ^47,48^. Therefore, we tested whether Sp1 was an intermediate between Pygo2/p53 and Kit. Sp1 was expressed at a lower level in PSP than PS, but showed no difference between cell lines derived from Pten/Smad4/p53 and Pten/Smad4/p53/Pygo2 tumors (**Extended Data Fig. 4l**), consistent with the hypothesis that Pygo2 sustains Sp1 expression in a p53-dependent manner. At the protein level, Sp1 was expressed at a higher level in pDKO tumors than in pTKO tumors, in concordance with the higher p53 modification and levels in pDKO tumors (**Fig. 4l**). The binding of Sp1 to *the Kit* promoter region was confirmed by the CUT&RUN-qPCR assay (**Fig. 4m**). Sp1 was silenced in PS cells with shRNA (**Extended Data Fig. 4M**), leading to downregulation of *Kit* expression (**Fig. 4n**) and spheroid formation **(Fig. 4o)**. Plus, Sp1 knockdown in PSP cells did not further reduce *Kit* expression or spheroid formation (**Fig. 4n-o**). We establish a previously uncharacterized pathway in PCa cells, where Pygo2 engages p53 to bind to the Sp1 promoter and sustain Sp1 expression, and Sp1 subsequently promotes Kit transcription and expression.

### Pygo2 downregulates T cell infiltration through the Kit-Ido1 pathway

To determine whether Kit is responsible for Pygo2 function in CTL impairment, we dissociated the syngeneic tumors formed by PS and PSP sublines with Kit knockdown or restoration (**Fig. 3**) and compared T cell infiltration frequencies. Kit knockdown in PS increased CD8^+^ T cell infiltration in PS tumors to the same level as in PSP tumors, and stable expression of the same Kit shRNA in PSP did not alter CD8^+^ T cell infiltration (**Fig. 5a-b**). Ectopic Kit expression in PSP cells brought the level of CD8^+^ T cells back to that of PS tumors. The CD8^+^-T/T_reg_ ratio increased after Kit knockdown in PS and decreased when Kit expression was restored in PSP (**Fig. 5a-b**). Kit expression on the tumor cell surface was confirmed using flow cytometry of freshly isolated tumors (**Fig. 5c**). By comparing PS-OVA and PSP-OVA with their respective Kit-knockdown sublines for OT-I T cell killing, we found that Kit silencing sensitized PS-OVA, but not PSP-OVA, to killing (**Fig. 5d-e**). Consistent with the critical role of p53 in Pygo2-Kit regulation, when Pten/Smad4/p53 and Pten/Smad4/p53/Pygo2 tumors were compared, the T cell subsets showed no differences (**Extended Data Fig. 5a**). These results indicate that Kit is the critical downstream mediator for Pygo2 to evade the anti-tumor effect from CTLs.

**Fig. 5.**
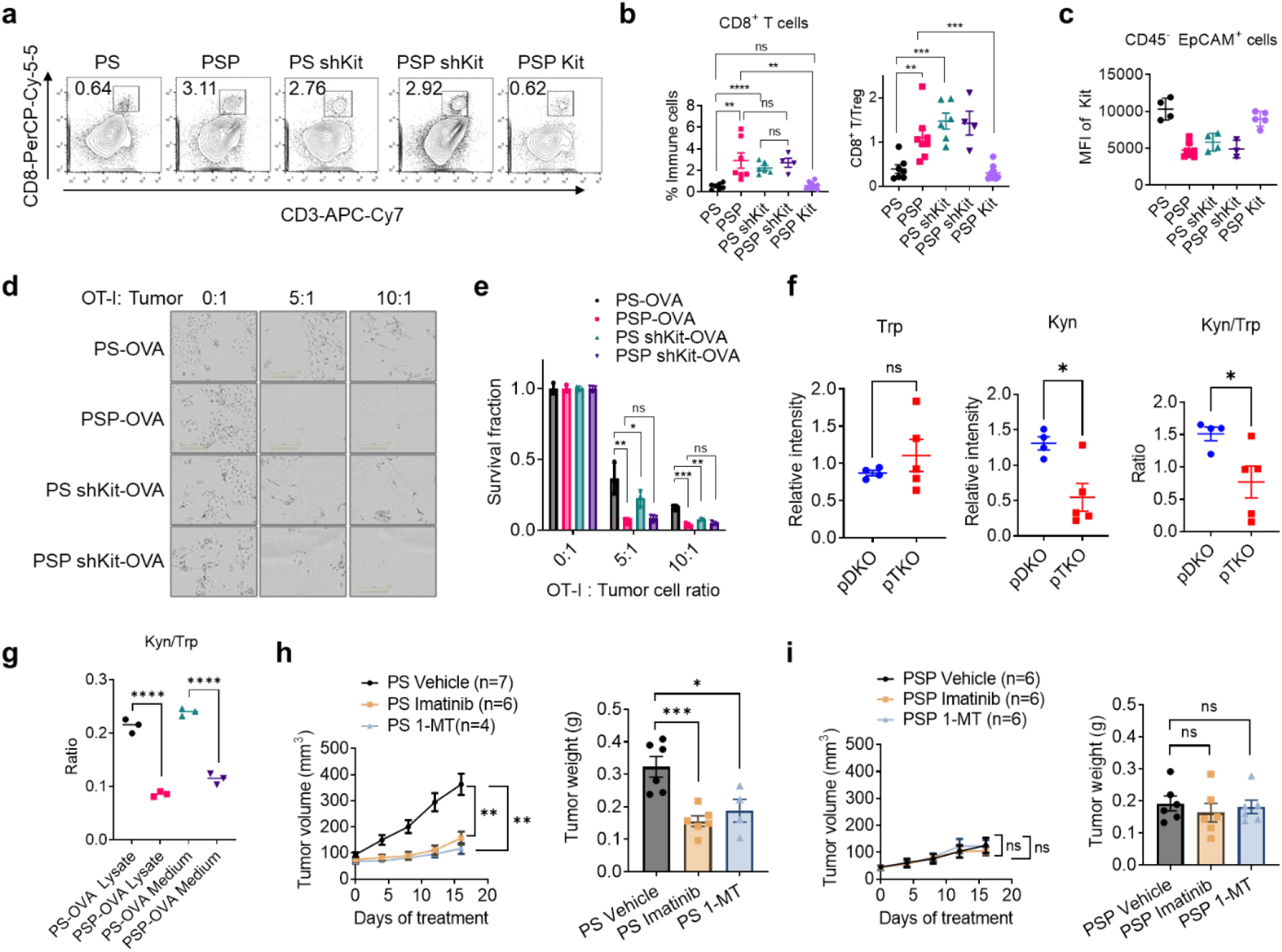
Pygo2 downregulates T cell infiltration through the Kit-Ido1 pathway. **(a)** Representative flow cytometry plots for measuring CD3^+^ CD8^+^ T lymphocyte infiltration in syngeneic tumors formed by PS, PSP and sublines. Values indicate the percentage of all CD45^+^ immune cells. **(b)** Quantification of tumor-infiltrating CD8^+^ T cells and the CD8^+^-T/T_reg_ ratio for syngeneic tumors (n = 4-9). **(c)** Kit protein level measured by flow cytometry on cancer cells freshly isolated from tumors formed by PS, PSP and sublines (n = 3-8). MFI, median fluorescence intensity. **(d-e)** T cell cytotoxicity assay to compare the killing of PS-OVA and PSP-OVA cells with and without *Kit* knockdown by antigen-stimulated OT-I T-cells at different E:T ratios (n=3). Viable cancer cells were detected by microscopy (E) and resazurin (F). **(f)** The normalized Trp and Kyn levels and their ratios in pDKO (n=4) and pTKO (n=5) tumor lysates, measured by mass spectrometry. **(g)** Kyn/Trp ratios measured by mass spectrometry for cell lysates and conditioned medium of PS-OVA (n=3) and PSP-OVA (n=3) co-cultured with OT-I T cells at ratio 1:1. **(h-i)** Growth curves and endpoint weight of syngeneic tumors formed by PS (n = 4-7) or PSP (n=6) in C57BL/6 mice and treated with vehicle, imatinib (50mg/kg, twice/daily), or 1-MT (400mg/kg, twice/daily). In (b)(c)(e)(f)(g)(h)(i), error bars represent SEM; ns, not significant, *P<0.05, **P<0.01, ***P<0.001, ****P<0.0001, Student’s t-test.

The Kit-Ido1 axis plays an important role in cancer cell-induced CTL dysregulation in GIST through producing immunosuppressive metabolites of tryptophan (Trp) ^39^. In PCa models, Kit and Ido1 shared the same expression pattern (**Fig. 3e, 3g, 4d**). We used mass spectrometry to quantify the relative abundances of Trp and the metabolite kynurenine (Kyn) in pDKO and pTKO tumor lysates. Although Trp levels showed no difference, Kyn and Kyn/Trp ratios were significantly lower in pTKO tumors (**Fig. 5f**). The Kyn/Trp ratio was also much higher in the lysate and medium of PS-OVA than PSP-OVA cells (**Fig. 5g**). Ido1 inhibitor 1-methyltryptophan (1-MT) enhanced OT-I T cell killing of PS cells (**Extended Data Fig. 5b**). To target the Kit-Ido1 axis *in vivo*, we treated mice bearing PS or PSP syngeneic tumors with imatinib or 1-MT and observed that both inhibitors significantly impeded PS tumors but not PSP tumors (**Fig. 5h-i**). At the TME level, imatinib and 1-MT treatments of PS tumors increased CD8^+^ and CD4^+^ T cells and decreased the T_reg_ fraction of CD4^+^ T cells (**Extended Data Fig. 5c**). However, CD8^+^ T cell infiltration was unaltered by the treatments in PSP tumors (**Extended Data Fig. 5d**). In conclusion, Kit-Ido1 cascade is the underlying mechanism for Pygo2-mediated CTL exclusion in PCa.

### Deletion of Pygo2 enhances efficacy from ICB, adoptive T cell therapy and CXCR2 inhibitor

Since Pygo2 extinction in PCa cells stimulated CTL infiltration *in vivo* and enhanced OT-I T-cell killing *in vitro*, Pygo2 ablation may enhance the efficacy of immunotherapies. We tested this hypothesis in different therapeutic contexts. To investigate the effect of Pygo2 knockout on ICB therapy, tumor-bearing C57BL/6 mice injected with RM9 control or Pygo2-knockout sublines were treated with isotype IgG or ICB antibodies (anti-PD1 plus anti-CTLA4). ICB decelerated but failed to eradicate any tumors. Pygo2 knockout led to 50% remission. Strikingly, the combination of ICB and Pygo2 knockout eliminated all RM9 tumors (**Fig. 6a-b**). In an ACT immunotherapy setting, stimulated OT-I CD8^+^ cells were infused into mice bearing tumors formed by OVA-expressed RM9 control or Pygo2 knockout sublines. A single dose of ACT slowed down Pygo2-knockout-OVA tumors more dramatically than control-OVA tumors (**Fig. 6c**). The ACT experiment was conducted in nude mice that lack T cells; therefore, the tumor-infiltrating CD8^+^ T cells should represent the infused OT-I CTLs. We observed a 4-fold higher CD8^+^ T cell infiltration in Pygo2-knockout-OVA tumors than in control-OVA tumors (**Fig. 6d**). Therefore, targeting Pygo2 has the potential to enhance both ICB and ACT immunotherapies in PCa.

**Fig. 6.**
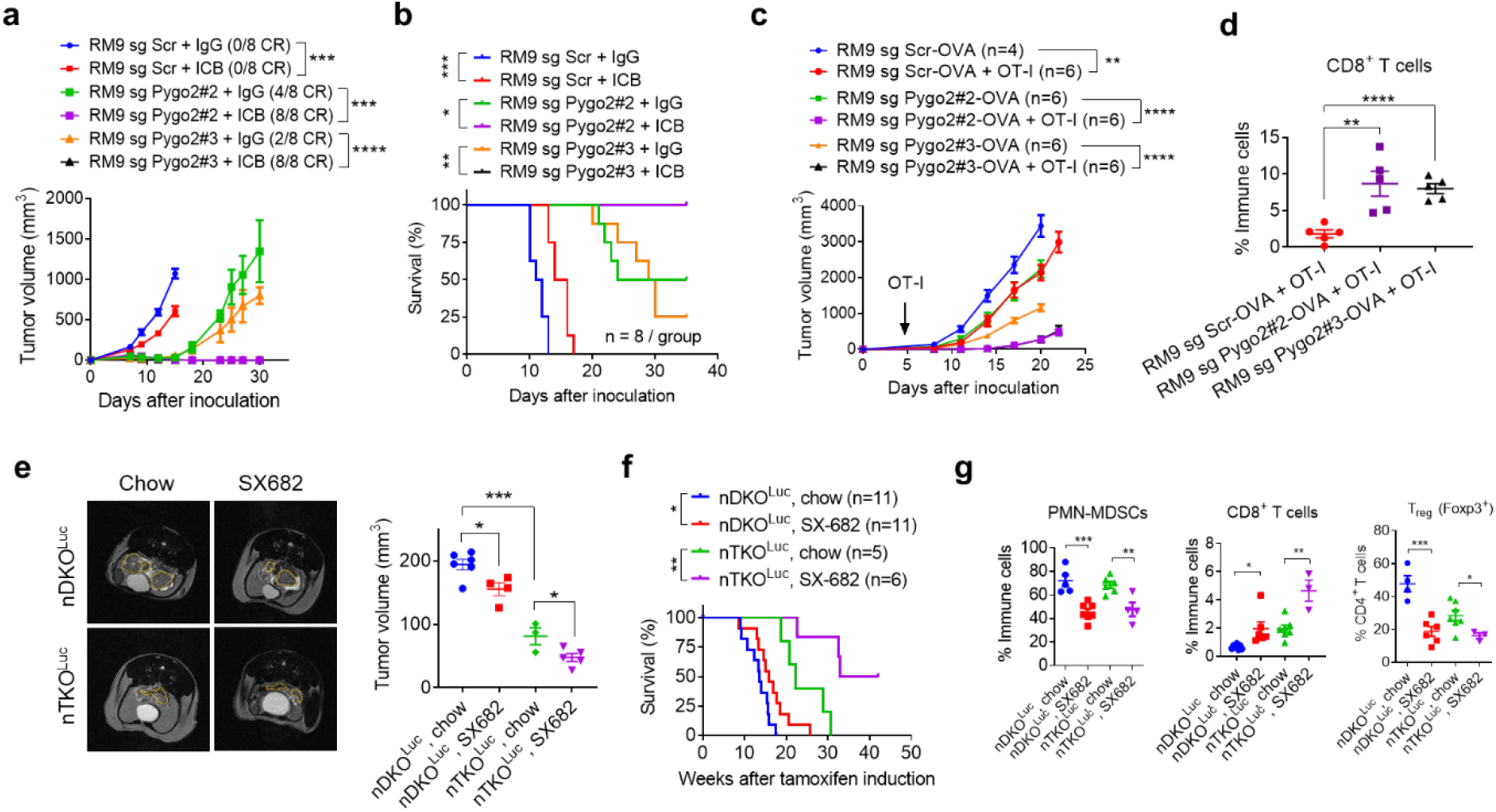
Deletion of Pygo2 enhances efficacy from ICB, adoptive T cell therapy and CXCR2 inhibitor. **(a-b)** Tumor growth curves and survival analysis for C57BL/6 mice inoculated with RM9 sgScr or sgPygo2 sublines and treated with IgG or ICB (anti-PD1 plus anti-CTLA4). n=8 for each group, CR, complete regression. **(c)** Tumor growth curves for ACT experiment, where nude mice inoculated with OVA-expressing RM9 sgScr or sgPygo2 sublines were infused with OVA-stimulated OT-I CD8^+^ T cells (1 × 10^7^) through tail vein at the arrow-indicated timepoint (n = 4 -6). **(d)** Flow cytometry quantification of tumor-infiltrating CD8^+^ T cells for nude mice bearing indicated tumors one week of OT-I T cell infusion. **(e)** Representative MRI images and tumor volume quantification of nDKO^Luc^ and nTKO^Luc^ mice fed with with standard chow or SX-682-admixed diet. Mice were imaged 12 weeks after tamoxifen induction, n = 3 - 6. Prostate regions were demarcated in yellow. **(f)** Kaplan-Meier curves for standard or SX-682 diet treated nDKO^Luc^ and nTKO^Luc^ mice (n = 5 - 11). **(g)** Flow cytometry quantification of tumor-infiltrating PMN-MDSCs, CD8^+^ T, T_reg_ (as fractions of CD4^+^ T cells) for standard or SX-682 diet treated nDKO^Luc^ and nTKO^Luc^ mice. In (a)(c)(d)(e)(g), error bars represent SEM. *P<0.05, **P<0.01, ***P<0.001, ****P<0.0001, Student’s t-test. In (b)(f), *P<0.05, **P<0.01, ***P<0.001, log-rank test.

PMN-MDSCs constitute a formidable barrier to anti-tumor T cell immunity in PCa. Blocking CXCR2 attenuated PMN-MDSC infiltration and decelerated pDKO tumor progression ^15^. Pygo2 loss in PCa cells did not affect PMN-MDSC abundance (**Extended Data Fig.2a**) or their immunosuppressive ability (**Extended Data Fig.6a**). Therefore, Pygo2-controlled and PMN-MDSC-mediated immunosuppressions were distinct, prompting a co-targeting strategy. CXCR1/2 antagonist SX-682 reduced PMN-MDSC infiltration in PCa ^49^, and the combination of SX-682 and anti-PD1 therapy are under evaluation in clinical trials (NCT03161431, NCT04599140). We induced spontaneous prostate tumors in nDKO^Luc^ and nTKO^Luc^ mice with tamoxifen, followed by administration of SX-682 medicated chow. SX-682-fed mice contained substantial SX-682 in circulation (**Extended Data Fig.6b**). Quantification of tumor volume with MRI indicated that although SX-682 was not as strong as Pygo2 loss to attenuate Pten/Smad4 tumor progression, nTKO^Luc^ mice fed the SX-682 diet developed the smallest tumors (**Fig. 6e**). Concordantly, SX-682-treated nTKO^Luc^ mice showed the most extended survival (**Fig. 6f**). SX-682 significantly reduced PMN-MDSCs and increased CD8^+^ T cells in the TME of nDKO^Luc^ and nTKO^Luc^ tumors (**Fig. 6g**). Although SX-682 did not affect the infiltration of total CD4^+^ T cells (**Extended Data Fig. 6c**), SX-682 significantly downregulated the fraction of T_reg_ in CD4^+^ T cells (**Fig. 6g**), which should also contribute to the overall reduced immunosuppression in the TME. In conclusion, Pygo2 extinction in PCa cells and PMN-MDSC blockade by targeting CXCR2 signaling alter the TME through complementary mechanisms, together eliciting stronger anti-tumor immunity.

### Pygo2 inhibitors antagonize PCa progression and enhance immunotherapy

Ali et al. identified a Pygo2 small-molecule inhibitor, JBC117, based on virtual screening for agents targeting the PHD domain of Pygo2, and showed the anti-tumor effect of JBC117 on colon and lung cancer xenografts ^50^. It was unclear whether the anti-tumor activity of JBC117 was dependent on Pygo2 expression. Here, we synthesized JBC117 and its analog JBC117ana, which lacks one hydroxyl group (**Fig. 7a, Extended Data Fig.7a-b**). The ability of JBC117 and JBC117ana to interrupt Pygo2 binding with H3K4me2 was verified by enzyme-linked immunosorbent assay (ELISA) (**Extended Data Fig.7c-e**). PS and PSP syngeneic tumors were treated with JBC117 or JBC117ana, and only the PS tumors were attenuated (**Fig. 7b-c**), supporting the on-target activity of JBC117. JBC117 or JBC117ana attenuated Kit and Ido1 expression in PS tumors but not in PSP tumors (**Fig. 7d**). We tested whether Pygo2 inhibitors sensitized PCa to immunotherapy. In the RM9 model, ICB and JBC117 exhibited single-agent anti-tumor activity, but the combination showed significantly more potent efficacy (**Fig. 7e**). Infiltration of CD8^+^ T cells, but not CD4^+^ T cells, was increased significantly by ICB or JBC117 treatment, and JBC117 enhanced the ability of ICB to downregulate T_reg_ (**Fig. 7f**).

**Fig. 7.**
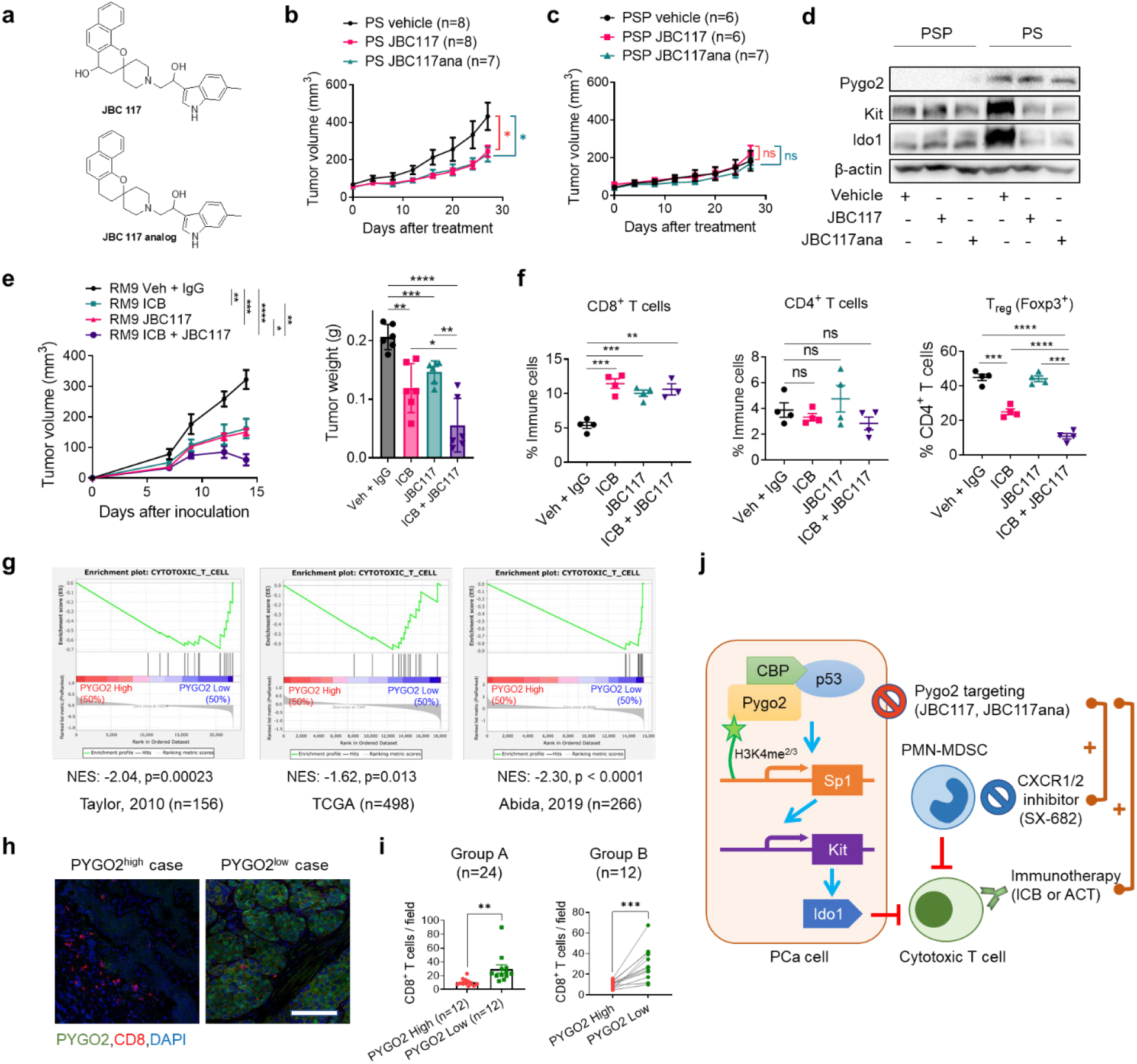
Pygo2 inhibitors antagonize PCa progression and enhance immunotherapy. **(a)** The structure of prototype Pygo2 inhibitors JBC117 and JBC117ana. **(b)** Growth curves for syngeneic tumors formed by PS treated with vehicle, JBC117 or JBC117ana (n = 7 - 8). **(c)** Growth curves for syngeneic tumors formed by PSP treated with vehicle, JBC117 or JBC117ana (n = 6 - 7). **(d)** Western blot to detect Pygo2, Kit and Ido1 in tumor lysates from the treatment groups. **(e)** Growth curves (left) and endpoint weight (right) for syngeneic tumors formed by RM9 and treated with control (vehicle plus IgG), ICB (anti-PD1 plus anti-CTLA4), JBC117, or ICB + JBC117 (n=6 per group). **(f)** Flow cytometry quantification of CD8^+^, CD4^+^, and T_reg_ cells from RM9 tumors treated with vehicle + IgG, ICB, JBC117, or ICB + JBC117 (n = 3 - 4). **(g)** GSEA result showing the enrichment of a CTL gene signature to PCa cases with low PYGO2 level than the ones with high PYGO2 level across 3 transcriptome databases. **(h)** Representative co-IF staining result of PYGO2 and CD8 in primary PCa with high PYGO2 (left) and low PYGO2 (right). Scale bar, 100µm. **(i)** Quantification of CD8 staining in primary PCa samples with relatively homogeneous PYGO2 staining pattern (Group A) or relatively heterogeneous PYGO2 staining pattern (Group B). In Group A (n=24), cases were stratified as high (intense) and low (weak) PYGO2 staining subgroups. In Group B (n=12), on the tissue sections, areas with high and low PYGO2 staining were demarcated for quantifying CD8^+^ T cells. **(j)** Schematic illustration of Pygo2 immuno-modulatory function and therapeutic opportunity in PCa. Pygo2 binds to H3K4me2/3 and engages CBP/p300 and p53 to activate Sp1 expression. Sp1 in turn upregulates Kit which induces Ido1 to impair CTLs and augment T_reg_ (not drawn). Therapeutically, Pygo2 inhibition decelerates prostate tumor growth and synergizes with different classes of immunotherapeutics to eradicate PCa. In (b)(c)(e)(f)(i), error bars represent SEM; ns, not significant, *P<0.05, **P<0.01, ***P<0.001, ****P<0.0001, Student’s t-test (paired for I, group B; unpaired for all others).

To explore the correlation between PYGO2 and CTL infiltration in human PCa, we performed GSEA of the CTL gene signature (**Supplementary Table 7**) on several transcriptome datasets of primary or metastatic human PCa ^43,44,51^. The CTL signature was enriched in patient samples with low *PYGO2* expression (**Fig. 7g**). Experimentally, immunofluorescence staining for PYGO2 and CD8 in archived samples of human primary PCa was performed (**Fig. 7h**). PYGO2 expression appeared heterogeneous in the tumor areas of some tumor samples, possibly due to the histological heterogeneity of human PCa. Therefore, we classified the samples into group A, composed of more homogeneous PYGO2 staining, and group B, composed of more heterogeneous PYGO2 expression. In Group A, samples with higher PYGO2 expression contained significantly fewer CD8^+^ T cells, whereas in Group B, tumor areas with higher PYGO2 expression also showed much lower CD8^+^ T cell infiltration (**Fig. 7i**). Therefore, clinical evidence supports the role of PYGO2 in excluding CTL infiltration.

## DISCUSSION

A combinatorial approach that fosters a TME conducive to immunotherapy is likely required to treat advanced PCa effectively. Recent studies highlighted the potential of sensitizing PCa to ICB therapy by inhibiting immunosuppressive myeloid cells, especially PMN-MDSCs, using CXCR1/2 antagonists, IL-23 blockade, and kinase inhibitors such as cabozantinib ^52^. Another powerful approach is to target cancer-cell-intrinsic mechanisms that simultaneously control cancer cell malignant functions and the formation of an immunosuppressive TME ^12,13^. This second approach has not been investigated in PCa. Here, we show that Pygo2, a PCa oncoprotein encoded by the 1q21.3 amplicon, initiates a signaling cascade that involves p53, Sp1, Kit, and Ido1 to reduce CTL infiltration and promote T_reg_ infiltration in the TME of PCa (**Fig. 7j**). Expression ablation or pharmacological inhibition of Pygo2 not only generated single-agent anti-tumor activity and prolonged the survival of PCa-bearing animals but also significantly enhanced the efficacy of ICB and ACT therapies. This newly revealed function of Pygo2, in addition to the cell-autonomous function of Pygo2 in promoting PCa cell proliferation ^17,30^, strongly supports the clinical development of Pygo2 inhibitors as a new avenue for the treatment of PCa and possibly other malignancies with high Pygo2 expression ^19-21,23-25^.

Previous studies focused on the cell-autonomous function of Pygo2 in cancer cell proliferation and stemness. Our study elucidates a Pygo2-controlled signaling axis that fosters a metabolically immunosuppressive TME in which CTL infiltration and activity are constrained. This finding echoes the therapeutically relevant theme that various oncogenic pathways play master regulatory roles in shaping the tumor immune landscape and dictating tumor resistance to immunotherapy ^12,13^. Consequently, pharmacological inhibition of Pygo2 can have two effects: thwarting tumor cell proliferation and enhancing CTL infiltration and activity. Because the histone code reader function of Pygo2 through binding between the PHD domain and H3K4me2/3 is required for Pygo2 to promote Kit expression (**Fig. 4i**), pharmacological targeting the PHD domain is a valid approach to diminish the revealed signaling mechanism.

Although the reader function of Pygo2 through binding to H3K4me2/3 appears invariant in Pygo2-involved gene regulation, the co-factors interacting with Pygo2 vary in a context-dependent manner. Pygo2 formed a complex with CBP/p300 and p53 (**Extended Data Fig. 4e-f**), and loss of Pygo2 reduced p53 acetylation, phosphorylation, and transcriptional activity (**Fig. 4e-f**). This result is consistent with the function of Pygo2 in inducing p53 accumulation and acetylation in hair follicle early progenitor cells ^28^. However, Pygo2-p53 regulation in the hair follicle context depended on β-catenin activation, whereas Pygo2 promoted the p53/Sp1/Kit/Ido1 axis in PCa cells independent of canonical Wnt/β-catenin signaling. This mechanism provides an example of the Wnt/β-catenin-independent mechanism of Pygo2 action, suggesting that the therapeutic inhibition of canonical Wnt signaling will not abolish the tumorigenic activity of Pygo2.

Both mouse and human PCa genetic data suggest p53 dependence of Pygo2 function (**Fig. 4c, 4g-h**). This result adds to the known complexity of p53 function in cancer progression ^53,54^. Pten/p53 mice developed PCa more rapidly than *PB-Cre4*^*+*^ *Pten*^*L/L*^ mice due to removing the p53-dependent cellular senescence ^55^. However, this tumor suppressor function of p53 was not manifested in the Pten/Smad4 background because the median survival for pDKO (Pten/Smad4) and Pten/Smad4/p53 mice was 16.4 and 17.9 weeks, respectively (**Fig. 1d, 4c**). This may be explained by the convergent function of p53 and TGFβ/Smad signaling to restrict early tumorigenesis ^56^. The p53 activity regulated by Pygo2 is insufficient to cause cell cycle arrest or apoptosis, and Pygo2 does not participate in p53-dependent cell cycle regulation.

Our results showed the co-occupancy of Pygo2 and p53 to the *Sp1* promoter (**Fig. 4j-k**). Previous studies using ChIP-seq identified Sp1 as a direct target and signaling intermediate effector of p53 ^57,58^. Our finding of Sp1 binding to *the Kit* promoter region (**Fig. 4m**) is consistent with reports showing the recruitment of Sp1 to the G-quadruplex-forming sites in the *KIT* promoter ^59^. More work is needed to identify the components of the TF complexes that control Sp1 and Kit expression in a Pygo2-dependent manner. It should be noted that Pygo2 may modulate immunosuppression through mechanisms in addition to the Kit/Ido1 pathway because PCa cells in pTKO mice were also enriched for innate immune pathways, such as interferon-α response, interferon-γ response, and JAK/STAT signaling (**Fig. 3b**). Our follow-up studies investigating other Pygo2 immune-modulatory mechanisms will help provide a complete picture. Because Pygo2 function in immunosuppression may go beyond the Kit/Ido1 pathway, it is reasonable to believe that targeting Pygo2 is more effective than inhibiting Kit or Ido1 in treating human PCa.

A significant contribution of this study is the translational implications. First, through genetic ablation of Pygo2, we demonstrated that all three immunotherapy approaches (ICB, ACT, and CXCR2 inhibitor) were significantly enhanced when Pygo2 was co-targeted (**Fig. 6**). Second, we synthesized JBC117 and JBC117ana as prototype Pygo2 inhibitors and showed that they largely phenocopied Pygo2 genetic deletion to generate both single-agent and combinatorial efficacy with ICB (**Fig. 7**). Nevertheless, the potency of JBC117 and JBC117ana to inhibit Pygo2-H3K3me2 interaction was moderate (**Extended Data Fig.7e**), prompting ongoing studies in our laboratory to perform virtual screening followed by ELISA-based validation to identify Pygo2 inhibitors with better drug-like properties.

## METHODS

### Mice

All animal work performed in this study was approved by the Institutional Animal Care and Use Committee at University of Notre Dame. All animals were maintained under pathogen-free conditions and cared for in accordance with the International Association for Assessment and Accreditation of Laboratory Animal Care policies and certification. *PB-Cre4, Pten*^*L/L*^, *Smad4*^*L/L*^, *p53*^*L/L*^, and *Rosa26-LSL-Luc*^*L/L*^ alleles have been previously described ^49^. *Pygo2*^*L/L*^ allele has been previously reported ^32^. *Nkx3*.*1*^*CreERT2*^ allele was obtained from the NCI mouse repository (strain number 01XBQ). All GEM mouse models were backcrossed to a C57BL/6 background for at least four generations. C57BL/6J (RRID: IMSR_JAX:000664) and OT-I (C57BL/6-Tg (TcraTcrb)1100Mjb/J, RRID: IMSR_JAX:003831) mice were purchased from Jackson Laboratory. Nude mice (RRID: IMSR_TAC:ncrnu) were purchased from Taconics.

### Cell Lines

TS3132 was previously isolated from Pten/Smad4 mice on a mixed background ^33^ and cultured in DMEM (GE Healthcare, SH30243.FS) supplemented with 10% fetal bovine serum (FBS, GE Healthcare, SH30396.03) and 100U/ml penicillin-streptomycin (Caisson Labs, PSL01). PS and PSP cell lines were established in this study from pDKO and pTKO tumors, respectively, and cultured in an optimized mouse prostate primary cell medium composed of DMEM/F12 (VWR, 45000-344), 10% FBS, 100U/ml penicillin-streptomycin, 10ng/ml EGF (Sigma-Aldrich, E4127), 20μg/ml adenine (Sigma-Aldrich, A3159),15mM HEPES (VWR, 16777-032), 5μg/ml insulin (Sigma-Aldrich, I-1882), 0.32μg/ml hydrocortisone (Sigma-Aldrich, H0888), and 10μM Y-27632 (ApexBio, B1293). PPS and PPSP cell lines established in this study were derived from prostate tumors of Pten/Smad4/p53 and Pten/Smad4/p53/Pygo2, respectively, and cultured in DMEM supplemented with 10% FBS and 100U/ml penicillin-streptomycin. All genotypes of these newly established cell lines were verified by genotyping. RM9 was purchased from ATCC (CRL-3312, RRID: CVCL_B461) and cultured in DMEM/F12 supplemented with 10% FBS and 100U/ml penicillin-streptomycin. All the cell lines were cultured at 37°C in a humidified incubator with 5% CO_2_. All cells were tested for mycoplasma-free status using a Mycoplasma Assay Kit (Agilent Technologies, 302109).

### Animal Experiments

For tamoxifen-inducible PCa models, nDKO^Luc^ and nTKO^Luc^ mice between 1-5 months were intraperitoneally (i. p.) injected with tamoxifen (Sigma-Aldrich, T5648) at 1mg in 100µl corn oil for 5 consecutive days to induce Cre activity and tumorigenesis. For SX-682 treatment experiments, nDKO^Luc^ mice 4-6 weeks post-tamoxifen and nTKO^Luc^ mice 8-10 weeks post-tamoxifen had similar tumor volumes; thus, they were fed with SX-682 medicated chow (Syntrix Pharmaceuticals) prepared at 1428.5 mg/kg (equivalent to 200 mg/kg mouse body weight/day) until the survival endpoint.

For syngeneic or allogenic primary tumor experiments, 1×10^6^ tumor cells were injected subcutaneously into C57BL/6 or nude male mice. For experimental metastasis experiments, 2×10^5^ tumor cells were injected intracardially into nude male mice. Metastatic tumors were monitored by bioluminescence imaging at the indicated time points. Mice were sacrificed eight weeks post-injection for necropsy.

For CD8^+^ T cell depletion experiments, mice with tumors reaching 50-100 mm^3^ were randomized to receive i.p. injection of an initial 400μg followed by 200μg anti-CD8 (BioXCell, BE0061) twice weekly or an equivalent dose of isotype IgG control. For targeted therapeutic experiments, mice with tumors reaching 50-100 mm^3^ were randomized to receive the following therapies at the reported doses: imatinib (MedChem Express. HY-50946) at 50 mg/kg, i.p., twice/day ^39^, 1-MT (Sigma-Aldrich, 452483) at 400 mg/kg, oral, twice/day ^39^, JBC117, and JBC117ana (synthesized in-house, see below) at 20 mg/kg, subcutaneously, daily ^50^. For ICB therapy using the RM9 model, 2×10^6^ RM9 sublines were inoculated into both the flanks of C57BL/6 male mice. Three days after inoculation, the mice were treated with anti-PD1 (BioLegend, 114116) and anti-CTLA4 (BioLegend, 106207) at 10mg/kg each, i.p., twice a week, or an equivalent dose of isotype IgG control. All treatments were continued until the specified experimental endpoints were reached.

### Adoptive OT-I T Cell Transfer

Splenocytes were isolated from the spleen of 6-10-week-old OT-I male mice and pulsed with 2ug/ml of OVA peptide SIINFEKL (VWR, H-4866.0001BA) for 4 h in T cell culture media composed of RPMI1640 (GE Healthcare, SH30027.01) supplemented with 10% FBS, 100U/ml penicillin-streptomycin, and 50μM 2-mercaptoethanol (VWR, 97064-880). Splenocytes were washed three times with PBS and seeded 1×10^7^ cells/well in 6-well plates. Cells were propagated every 1-2 days and T cells proliferated to form clusters. After 3-5 days, T cells were washed three times with PBS, and 1×10^7^ cells in 100ul PBS were intravenously injected into nude mice inoculated with RM9 sublines 3 days before. Tumor-bearing mice were monitored until the specified endpoint.

### Non-invasive Animal Imaging

MRI imaging with 1T ICON (Bruker) and bioluminescence imaging with a Spectral Ami HT Advanced Molecular Imager (Spectral Instruments Imaging) were performed at the Notre Dame Integrated Imaging Facility, following our previous report ^49^. MRI image sequences were loaded into ImageJ (RRID: SCR_003070) to manually demarcate the contour of the prostate on each plane and to calculate the total volume by integration.

### SX-682 Measurement in Mice Plasma

Plasma was isolated from the peripheral blood of mice treated with the standard or SX-682 diet (Syntrix Pharmaceuticals) for one month. The calibration sample was prepared by diluting a stock solution containing a known amount of SX-682 in C57BL/6 mouse plasma. A 20ul aliquot of each sample was then diluted 1/4 into an internal standard solution (acetonitrile + 50 ng/ml SX-517). The resulting suspension was briefly vortexed and centrifuged at 10,000 rpm for 10 min. The supernatants were then transferred to HPLC vials for analysis. The peak areas for SX-682 were integrated, and the SX-682 concentrations were calculated using a formula derived from the calibration curve.

### Immunohistochemistry (IHC), Immunofluorescence (IF), and Western Blot

Animal tissues were fixed overnight in 10% formalin and embedded in paraffin. IHC and IF staining were performed as previously described ^49^. Antigen retrieval was performed by heating in a pressure cooker at 95°C for 30 min, followed by 115°C for 1 min in citrate-unmasking buffer (pH 6.0). The IHC slides were scanned using an Aperio ScanScope (Leica). For IF staining of human FFPE specimens, the tumor areas were demarcated based on pathological inspection of H&E staining. IF slides were imaged with an A1R confocal laser microscope (Nikon), and CD8^+^ T cells were counted manually. Primary antibodies used included Pygo2 (clone S3I4, previously reported ^30^), Ki67 (Fisher Scientific, RM-9106-S1), cleaved caspase 3 (Cell Signaling Technology, 9661), Kit (Cell Signaling Technology, 3074), Ido1 (Santa Cruz Biotechnology, sc137012), CD3 (DAKO, A0452), β-catenin (Cell Signaling Technology, 8480), mouse CD8a (Cell Signaling Technology, 98941), and human CD8a (Biolegend, 372902).

For western blot, cells or fresh tissues were lysed on ice using RIPA buffer supplemented with protease inhibitors (Bimake, B14012) and phosphatase inhibitors (Roche, 04906845001). Immunoblotting was performed as described previously ^49^. The following primary antibodies were used: Pygo2 (clone S3I4), β-actin (Santa Cruz Biotechnology, sc-47778), kit (Santa Cruz Biotechnology, sc-13508), Erk (Cell Signaling Technology, 4695), phospho-Erk1/2 (Cell Signaling Technology, 4370), p53 (Cell Signaling Technology, 2524), Akt (Cell Signaling Technology, 2920), phospho-Akt (Cell Signaling Technology, 4060), Ido1 (Santa Cruz Biotechnology, sc137012), CBP (Cell Signaling Technology, 7389), phospho-p53 (Cell Signaling Technology, 9284), acetyl-p53 (Cell Signaling Technology, 2525), and Sp1 (Santa Cruz Biotechnology, sc-420).

### CyTOF and Flow Cytometry for Intratumoral Immunocytes

Tumors were minced into homogenate and rotated at 37°C in dissociation media, DMEM with 10% FBS and 1 mg/ml collagenase IV (STEMCELL Technologies, 07427) for 1 h, followed by passing through 40μm strainers. Erythrocytes were depleted via hypotonic lysis. The CyTOF procedure and antibody panel have been described previously ^15^. The samples were analyzed with Helios CyTOF mass cytometer (Fluidigm) in the Flow Cytometry and Cellular Imaging Core Facility at the MD Anderson Cancer Center. Flow cytometry samples were prepared as described previously ^49^ and run on CytoFLEX S (Beckman Coulter). CyTOF and flow cytometry data were analyzed using FlowJo v10.8 (FlowJo, RRID: SCR_008520) or CytExpert (Beckman Coulter). Fluorochrome-conjugated antibodies included CD8a (Tonbo Biosciences, 65-0081), CD4 (Tonbo Biosciences, 35-0042), Foxp3 (Tonbo Biosciences, 20-5773), CD3 (Tonbo Biosciences, 25-0032), CD45 (Tonbo Biosciences, 60-0451), CD45 (Tonbo Biosciences, 35-0451), CD11b (Tonbo Biosciences, 65-0112), Gr-1 (Tonbo Biosciences, 60-5931), F4/80 (Tonbo Biosciences, 25-4801), kit (Tonbo Biosciences, 20-1172), and EpCAM (BioLegend, 118215).

### T Cell-PMN-MDSC Co-culture Assay and T Cell Killing Assay

T cell and PMN-MDSC co-culture assays were used to assess the immunosuppressive activity of PMN-MDSCs following our previous method ^49^. CD3^+^ T cells were isolated from the spleens of wild-type C57BL/6 mice, whereas PMN-MDSCs were isolated from nDKO^Luc^ and nTKO^Luc^ tumors. PMN-MDSCs and T cells were co-cultured in a 2:1 ratio. For the antigen-dependent T-cell killing assay, OVA-overexpressing cancer cells were seeded at 5,000 cells/well in 96-well plates. After the cells were attached to the plate, OT-I T cells stimulated in the same manner as in the ACT experiment (see above) were added to the cancer cells at the specified E:T ratios. After 24-48h of co-culture, T cells were washed away and viable cancer cells were measured using the resazurin assay.

### Tumor Cell Proliferation Assays

For the 2D colony formation assay, cancer cells were seeded at 100/well into 24-well plates and cultured for 5-7 days before fixation and crystal violet staining. The colonies were counted manually. For the 3D spheroid assay, cancer cells at a density of 2,000/10μl culture media were mixed with 20μl Matrigel (Corning, 354230), then dropped to the center of the wells of a 24-well plate, followed by flipping over the plate and incubation for 15-30 min at 37°C. The plate was flipped back, and prewarmed mouse prostate primary cell medium (see recipe above) was added. Spheroids were formed within 5-7 days and imaged for manual counting.

### Cell Migration Assay

Cells were seeded at 5×10^5^ cells in 200μl serum-free DMEM in the upper chamber of the inserts (Celltreat, 230639). DMSO or imatinib (4µM) was then added to the cells. The inserts were placed in 24-well plates containing DMEM with 10% FBS as chemoattractant. After 24 h, cells were fixed and stained with crystal violet. Cells on the top of the insert membranes were wiped away, and cells at the bottom of the membranes were imaged and counted.

### Reporter Assay

For the p53 reporter assay, PG13-Luc (Addgene, 16442) was transfected into PS and PSP cells with jetOPTIMUS (Polyplus, 101000051) followed by 10μM nutlin-3 (Cayman Chemical Company, 10004372), 500ng/ml CPT (Chem-Impex, 22069), or 2μM doxorubicin (LC Laboratories, D-4000) treatment for 24h. Luciferase activity was measured using a SpectraMax Gemini EM microplate reader (Molecular Devices).

### CRISPR/cas9, shRNA and Gene Overexpression

To generate Pygo2 CRPSR/cas9-knockout cells, three different CRISPR/Cas9 sgRNA designs in an all-in-one lentiviral vector (ABM, 382541140595) were purchased. Lentivirus was packaged to infect target cells following a previous report ^15^. After puromycin selection, single-cell clones were expanded and screened by western blot to validate the Pygo2 knockout. For shRNA knockdown, all lentiviral shRNA clones targeting *Sp1, Kit*, and non-targeting shRNA control were obtained from Sigma-Aldrich in the pLKO vector and were prepared as previously described ^15^. Cell lines stably overexpressing EβC (Addgene, 24312) were generated by sorting mCherry^+^ cells. Mouse *Pygo2* and *Kit* were subcloned from the original vectors (OriGene Technologies, MR206368; Sino Biological, MG50530-CH) into the pMSCV-puro retroviral backbone (Addgene, 68469). The HA tag was added to the Pygo2 C-terminus by adding the HA coding sequence to the PCR primer. Mouse Pygo2 mutants W351A and Y326A, corresponding to human PYGO2 W352 and Y327, respectively, were generated using the Phusion Site-Directed Mutagenesis Kit (Thermo Fisher Scientific, F541). All stable cell lines were selected using 2μg/ml of puromycin (Goldbio, P-600). For OVA overexpression, full-length OVA was subcloned from the original vector (Addgene, 64599) to the EF1a-FOXA1-P2A-Hygro lentiviral vector (Addgene, 120438) with FOXA1 replaced with OVA. Stable cells were selected using 200μg/ml hygromycin B.

### Quantitative RT-PCR (qRT-PCR)

RNA was isolated using the RNeasy Kit (Bio Basic, BS1361) and reverse transcribed using the All-in-One cDNA Synthesis Kit (Bimake, B24403). qRT-PCR was performed using SYBR Green qPCR Master Mix (Bimake, B21202). *Gapdh* was used for normalization. Student’s t-test was performed based on the ΔΔC_T_ values. Unless otherwise specified, n=3 biological replicates per group were used for all qRT-PCR experiments. Primer sequences are listed in **Supplementary Table 8**.

### CUT&RUN Assay Followed by Sequencing or qPCR

CUT&RUN experiments were conducted using 200,000 PS or PSP cells using the CUT&RUN assay kit (Cell Signaling Technology, 86652). Briefly, the cells were washed and bound to concanavalin A-coated magnetic beads. Next, permeabilized cells were incubated with IgG (Cell Signaling Technology, 66362), antibodies against H3K4me3 (Cell Signaling Technology, 9751), Sp1 (Santa Cruz Biotechnology, sc-420), p53 (Cell Signaling Technology, 2524), or Pygo2 (clone S3I4) for 2h at 4°C with rotation. The cell-bead slurry was washed and incubated with protein A-MNase for 1h at 4°C with rotation. CaCl_2_ was added to the cell-bead slurry to initiate protein A-MNase digestion, and the reaction was incubated at 4°C for 30 min. The reaction was stopped with stop buffer containing 50pg of spike-in DNA. Digested DNA was extracted and purified using DNA purification spin columns (Cell Signaling Technology, 14209). The input samples were sonicated for 12 min with a Covaris S220 Ultrasonicator System to obtain a fragment size between 150-300bp. For sequencing, we prepared a library using the SimpleChIP ChIP-seq DNA Library Prep Kit for Illumina (Cell Signaling Technology, 56795). The library was sequenced using MiSeq (Illumina) in the Genomic & Bioinformatics Core Facility at the University of Notre Dame. Data were analyzed using Galaxy (https://usegalaxy.org/). The reads were aligned to the mm10 reference genome. Peak calling was performed using MACS2 software. To validate individual binding, qPCR was performed using SYBR Green qPCR master mix (Bimake, B21202). The enrichment of the *Kit* and *Sp1* promoter regions was calculated relative to the IgG control. For *Sp1* promoter region, forward primer 5’-TAATTGGCTGTTCGTTCACGTC-3’; reverse primer 5’-GGAGCAAGCTTCCTAAACCA-3’. For *Kit* promoter region, forward primer 5’-AGCGTCCTCTCTCCGA-3’; reverse primer 5’-CCGCAAGAAAAGGCTCT-3’.

### Microarray and Genomic Data Analysis

Single cells from spontaneous prostate tumors of pDKO and pTKO mice were isolated by digesting the tumors with 10mg/ml of *Bacillus Licheniformis* protease (Creative Enzymes, NATE0633) and 0.2 mg/ml DNase I (Sigma-Aldrich, 10104159001) on ice for 40 min, followed by passing through 70μm cell strainers. Cancer cells were purified using the MojoSort Mouse CD326/EpCAM Selection Kit (BioLegend, 480141), followed by RNA extraction using RNeasy Kit (Qiagen). RNA samples were profiled on the Mouse Genome 430 2.0 Array (Affymetrix) at the Genomics Core Facility at the MD Anderson Cancer Center. The data were analyzed using Transcriptome Analysis Console (TAC) software (Thermo Fisher) to generate a list of differentially expressed genes with a fold-change cutoff of over 1.5-fold and P-value < 0.05. Pathway enrichment was performed using GSEA software (UC San Diego and Broad Institute, RRID:SCR_003199). The transcription factor enrichment was performed using Enrichr (RRID: SCR_001575). Upstream regulator prediction and gene regulation connections for Kit and Pygo2 were conducted with Ingenuity Pathway Analysis (IPA) software (QIAGEN, RRID:SCR_008653).

To analyze the correlation between PYGO2 expression and CTL gene signature using published human PCa transcriptomic data, we downloaded transcriptome data of three studies ^43,44,51^ from cBioPortal. For each dataset, the samples were grouped as high and low based on the normalized *PYGO2* level. Differential analyses between the PYGO2 high group and the PYGO2 low group were performed using limma ^60^. Log2(fold change) of gene expression between the two groups was used to run GSEA of a CTL gene signature (Supplementary Table 7), curated based on Szabo et al ^61^.

### Tryptophan and Kynurenine Detection

To detect metabolites in prostate tumor tissues, 30mg of tissue was homogenized in 600μl dissolving solution (40:40:20 methanol: acetonitrile: water with 0.5% formic acid). The homogenate was centrifuged at 16,000 × g and 4°C for 10 min. The supernatant was collected, neutralized with 30ul of 15% NH_4_HCO_3_, and ready for detection. To detect metabolites in the cancer cells and conditioned medium, PS-OVA and PSP-OVA cell lines were first co-cultured with OVA-pulsed OT-I T cells at a 1:1 ratio for 24h. Cells were washed with cold PBS three times to remove T cells, and then 1ml dissolving solution was added and incubated on ice for 5 min, followed by adding 50ul 15% NH_4_HCO_3_. The cells were then scraped from the plate and transferred to a 1.5ml tube, followed by centrifugation at 16,000 × g at 4°C for 10 min. The supernatant was then collected for detection. For metabolite extraction from the conditioned medium, after co-culture with T cells, the medium was collected and centrifuged at 1000 g for 5 min and passed through a 0.22μm filter. From the medium, 50ul was transferred to a 1.5ml tube followed by adding 200μl ice-cold methanol and incubation at -20°C for 20 min. Next, the samples were centrifuged at 16,000 × g at 4°C for 10 min, and the supernatant was collected and set aside. The pellet was dissolved with 1ml dissolving solution without formic acid and incubated for 10 min on ice. After centrifugation at 16,000 × g at 4°C for 10 min, the supernatant was collected and combined with the methanol-extracted supernatant as the final sample for detection. To detect tryptophan and kynurenine, all samples were run on a Thermo Q-Exactive MS/MS coupled with a Thermo UPLC system at the Metabolomics Core Facility at the Rutgers-Robert Wood Johnson Medical School. The relative intensities of tryptophan and kynurenine were calculated by normalizing the intensity of the individual samples to the mean intensity of all samples.

### ELISA for Pygo2

An ELISA for Pygo2 was designed by coating a streptavidin-coated 96-well plate (Thermo Fisher, 15125) with biotinylated 21-mer H3K4me2 peptide (Active Motif, 81041) at 0.5µg/ml of 2h, room temperature. Next, the wells were incubated with recombinant human PYGO2 (LSBio, LS-G26167) at 2µg/ml with compounds the test compounds (DMSO, JBC117, JBC117ana) at different concentrations. One hour later, free-floating rhPYGO2 was washed off. PYGO2 antibody (R&D, MAB3616) and HRP-conjugated secondary antibody were added sequentially, with washing between the steps. TMB substrate (BioLegend, 421101) was added after secondary antibody incubation for signal development detected at 450nm using an Epoch 2 microplate spectrophotometer (BioTek Instruments). If the compound interferes with PYGO2-H3K4me2 binding, the reading at 450 nm is expected to be reduced.

### Co-Immunoprecipitation (Co-IP)

PS Cells were extracted in IP buffer (25 mM Tris-HCl pH 7.4, 150mM NaCl, 1mM EDTA, 1% NP-40 and 5% glycerol) supplemented with protease inhibitors (Bimake, Cat# B14012) and phosphatase inhibitors (Roche, Cat# 04906845001) and sonicated for 30 sec. The cell extracts were incubated with IgG (Cell Signaling Technology, 3900), p53 (Cell Signaling Technology, 2524S), Pygo2 (S3I4), or CBP (Cell Signaling Technology, 7389) antibody at 1:200 dilution, 4°C overnight with rotation. Next, the samples were pulled down with protein A agarose beads (Cell Signaling Technology, 9863) and washed with 1x loading buffer for western blot detection.

### Proximity Ligation Assay (PLA)

PLA was carried out on 4% paraformaldehyde-fixed PS and PSP cells using the Duolink PLA Kit (Sigma-Aldrich, DUO92101-1KT) following the manufacturer protocol. Pygo2 (S3I4) and p53 (Cell Signaling Technology, 2524) antibodies were used in this assay, rabbit IgG (Cell Signaling Technology, 3900) and mouse IgG (Santa Cruz Biotechnology, sc-2025) were used as control. The slides were imaged with an A1R confocal laser microscope (Nikon). The positive dots in cell nuclei were quantified manually.

### Clinical Samples

Formalin-fixed paraffin-embedded (FFPE) slides of primary prostate tumors harvested by transurethral resection of the prostate (TURP) or radical prostatectomy were obtained from the tissue bank at the Indiana University School of Medicine. All specimens were de-identified. The experiments were approved by the IRB protocols of Indiana University School of Medicine (IRB#1808872882) and University of Notre Dame (20-03-5926). The clinical characteristics of the samples are summarized in **Supplementary Table 9**.

### JBC117 and JBC117ana Synthesis

The putative Pygo2 small-molecule inhibitor, JBC117, was first discovered by Ali et al. ^50^, but its synthesis method was not described. Our in-house syntheses of JBC117 and JBC117ana were conducted by the Warren Center for Drug Discovery and Development at the University of Notre Dame. The synthetic routes are illustrated in Extended Data Fig. 7a-b.

JBC117 was purified using a reverse-phase reverse phase Yamazen column (MPLC) using 30% acetonitrile: H_2_O (likely using a neutral mobile phase). The purity (>95%) of JBC117 was determined using HPLC analysis. ^1^H NMR (400 MHz, MeOD) δ 8.17 – 8.15 (m, 1H), 7.65 (ddt, *J* = 7.9, 5.3, 2.4 Hz, 1H), 7.48 (dt, *J* = 8.2, 2.8 Hz, 1H), 7.41 (d, *J* = 8.5 Hz, 1H), 7.39 – 7.29 (m, 3H), 7.07 (d, *J* = 14.8 Hz, 2H), 6.78 (dt, *J* = 8.1, 1.6 Hz, 1H), 5.19 (dt, *J* = 9.9, 3.4 Hz, 1H), 4.84 (t, *J* = 6.4 Hz, 1H), 3.20 – 2.87 (m, 6H), 2.31 (d, *J* = 2.8 Hz, 3H), 2.21 – 2.09 (m, 2H), 2.02 – 1.81 (m, 4H).; ^13^C NMR (101 MHz, MeOD) δ 147.16, 137.44, 134.28, 130.81, 127.11, 125.90, 125.41, 125.39, 124.89, 123.75, 121.66, 121.11, 120.28, 119.39, 118.48, 118.26, 116.56, 110.87, 73.75, 64.94, 64.65, 62.25, 50.12, 49.99, 41.32, 35.44, 35.35, 33.23, 33.15, 20.42.; HRMS: Calcd. for C28H31N2O3+ [M+H]+ 443.2319 found 427.2320.

JBC117ana was purified using a reverse-phase reverse phase Yamazen column (MPLC) using 30% acetonitrile: H_2_O (likely using a neutral mobile phase), followed by recrystallization. The purity of JBC117ana was determined by HPLC analysis (The compound was partially soluble and formed an eliminated product in the mobile phase system (AcCN: H_2_O and MeOH: H_2_O containing 0.1%TFA). ^1^H NMR (400 MHz, DMSO) δ 10.72 (s, 1H), 8.42 – 8.00 (m, 1H), 7.80 (dd, *J* = 7.2, 2.1 Hz, 1H), 7.55 – 7.41 (m, 3H), 7.36 (d, *J* = 8.4 Hz, 1H), 7.21 (d, *J* = 8.3 Hz, 1H), 7.14 (d, *J* = 11.9 Hz, 2H), 6.80 (dd, *J* = 8.1, 1.5 Hz, 1H), 5.02 (s, 1H), 2.96 – 2.53 (m, 8H), 2.38 (s, 3H), 1.95 – 1.67 (m, 6H). ^13^C NMR (101 MHz, DMSO) δ 147.71, 137.33, 137.17, 133.35, 130.27, 128.35, 127.86, 126.00, 125.63, 125.59, 124.31, 124.28, 121.92, 121.76, 121.43, 120.52, 119.55, 119.38, 118.04, 115.64, 111.69, 111.63, 73.28, 65.70, 64.52, 50.28, 49.23, 34.51, 31.63, 21.88, 21.72.; HRMS: Calcd. for C28H31N2O2+ [M+H]+ 427.2384 found 427.2380.

### Statistical Analysis

Statistical analyses were performed using GraphPad Prism v8.0 (RRID: SCR_002798). Unless otherwise mentioned, all data are presented as mean ± SEM (standard error of the mean). Sample sizes, error bars, P values, and statistical methods are shown in the Figures. legends. Statistical significance was defined as P < 0.05.

### Data Availability

Microarray data are available in the Gene Expression Omnibus (GEO) (RRID: SCR_005012) with accession number GSE195948. CUT&RUN-seq data are available at GEO with accession number GSE196486. Other data generated in this study are available in the article and its data files. Further information and requests for resources and reagents should be directed towards lead contact.

## Author Contributions

**Y. Zhu:** Conceptualization, investigation, methodology, data curation, formal analysis, validation, visualization, project administration, writing – original draft, writing – review & editing. **Y. Zhao:** investigation, methodology. **J. Wen**: investigation. **S. Liu**: software, visualization, formal analysis, writing – original draft. **T. Huang**: investigation. **I. Hatial**: resources, methodology, writing – original draft. **X. Peng**: investigation. **HA. Janabi**: investigation. **G. Huang**: investigation. **J. Mittlesteadt**: investigation. **M. Cheng**: resources. **B. Ashfeld**: resources. **A. Bhardwaj**: investigation. **KR. Kao**: resources. **DY. Maeda**: resources. **X. Dai**: resources. **O. Wiest**: supervision. **B. Blagg**: supervision. **Xuemin Lu**: supervision. **L. Cheng**: resources, data curation, supervision, funding acquisition. **J. Wan**: formal analysis, supervision, funding acquisition. **Xin Lu**: Conceptualization, investigation, resources, supervision, funding acquisition, writing – original draft, writing – review & editing.

## Acknowledgments

We would like to thank the Lu lab members, Siyuan Zhang, Mary Ann McDowell, Jun Li, Crislyn D’Souza-Schorey, Zach Schafer, Sharon Stack, and Kasturi Haldar, for their essential comments and suggestions during this work. We are grateful for the support from various core facilities used in this study, including the Freimann Life Science Center (Teri Highbaugh), Genomics and Bioinformatics Core Facility (Michael Pfrender, Melissa Stephens, Jacqueline Lopez, Brent Harker), Integrated Imaging Facility (Sara Cole, Sarah Chapman), and Metabolomics Core Facility (Xiaoyang Su, Yujue Wang). This work was supported by the National Institutes of Health grant R01CA248033 (X. Lu). Other support included the Department of Defense CDMRP PCRP grants W81XWH2010312 (X. Lu) and W81XWH2010332 (X. Lu), Elsa U. Pardee Foundation grant (X. Lu), Indiana CTSI pilot grant (X. Lu) through NIH NCATS CTSA grant UL1TR002529 (A. Shekhar, PI), National Institutes of Health grant P30CA082709 (J. Wan), and Boler Family Foundation endowment (X. Lu) at University of Notre Dame.

## EXTENDED DATA

**Extended Data Fig. 1.**
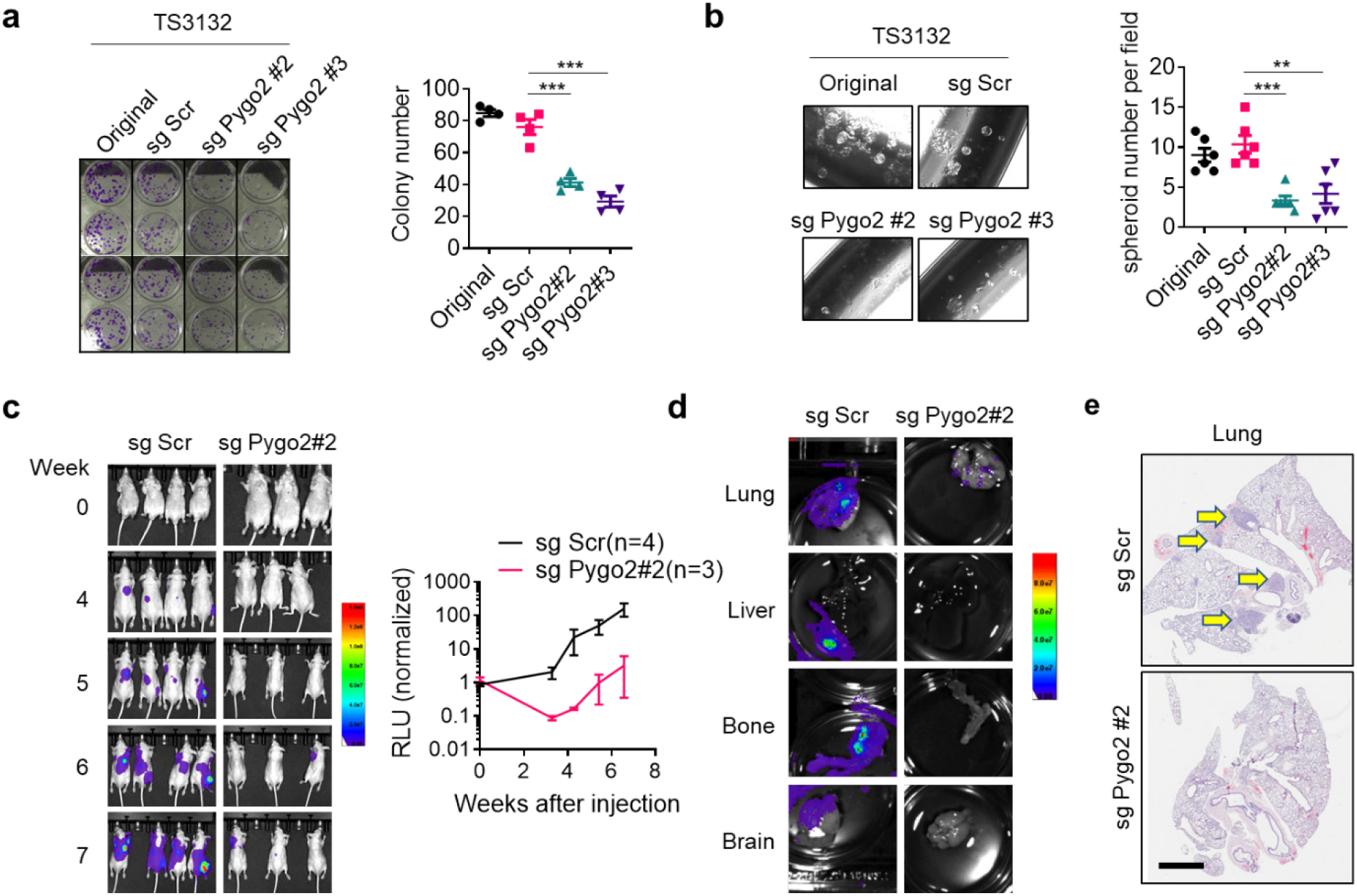
Pygo2 promotes prostate cancer proliferation and metastasis in xenograft models. **(a)** Two-dimensional colony formation by TS3132 control and Pygo2-knockout sublines. ***P<0.001, Student’s t-test (n=4). **(b)** Tumor spheroid assay for TS3132 control and Pygo2-knockout sublines. **P<0.01, ***P<0.001, Student’s t-test (n=6). **(c)** Bioluminescence images and quantification of metastasis signals in nude mice after intracardiac injection of TS3132-sgScr (n=4) or TS3132-sgPygo2#2 (n=3). **(d)** Representative bioluminescence images of various organs from nude mice injected with TS3132-sgScr or TS3132-sgPygo2#2. **(e)** H&E staining for lungs from nude mice injected with TS3132-sgScr or TS3132-sgPygo2#2. Yellow arrows denote metastasis nodules. Scale bar, 2mm.

**Extended Data Fig. 2.**
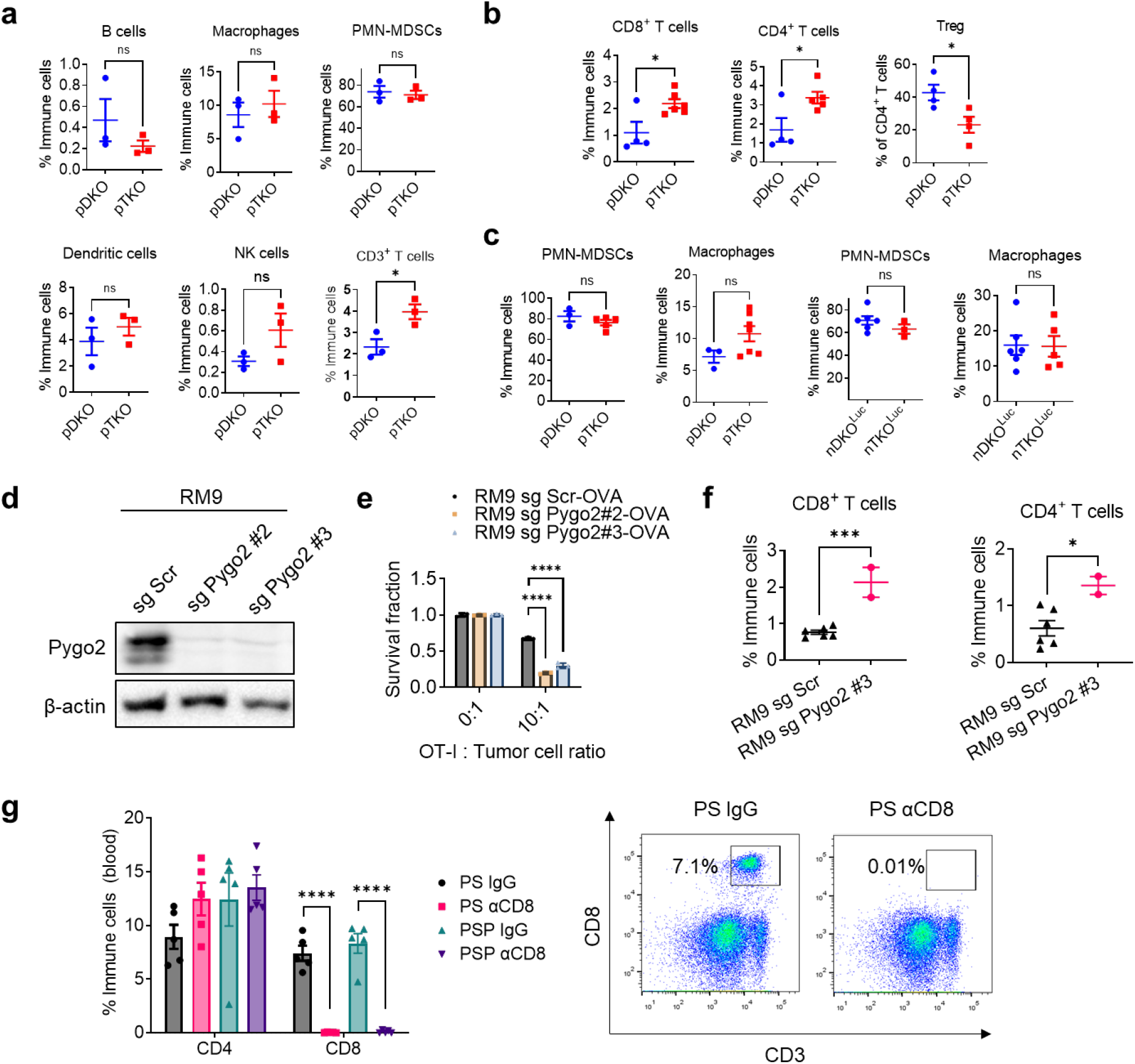
Pygo2 inhibits CTL infiltration and attenuates OT-I T cell killing of RM9-OVA cells. **(a)** Immune cell infiltration quantified with CyTOF for pDKO and pTKO tumors (n=3). **(b)** Flow cytometry quantification of tumor-infiltrating T cell subsets in pDKO and pTKO tumors (n = 4-6). **(c)** Flow cytometry quantification of tumor-infiltrating PMN-MDSCs (CD11b^+^ Gr1^high^) and macrophages (CD11b^+^ F4/80^+^) in pDKO and pTKO tumors as well as nDKO^LUC^ and nTKO^LUC^ tumors (n = 3-7). **(d)** Pygo2 expression in RM9 sgScr and sgPygo2 sublines, detected by western blot. **(e)** T cell cytotoxicity assay to compare the killing of RM9 sgScr and sgPygo2 sublines by antigen-stimulated OT-I T-cells. Viable cancer cells were detected by resazurin (n=3). **(f)** Flow cytometry quantification of CD8^+^ and CD4^+^ T cells in syngeneic tumors formed by RM9 sgScr (n=6) and sgPygo2 sublines (n=2). **(g)** Flow cytometry to confirm the depletion of CD8^+^ T cells (but not CD4^+^ T cells) in the blood by anti-CD8 antibody treatment (n=5). In all panels, error bars represent SEM; ns, not significant; *P<0.05, ***P<0.001, ****P<0.0001, Student’s t-test.

**Extended Data Fig. 3.**
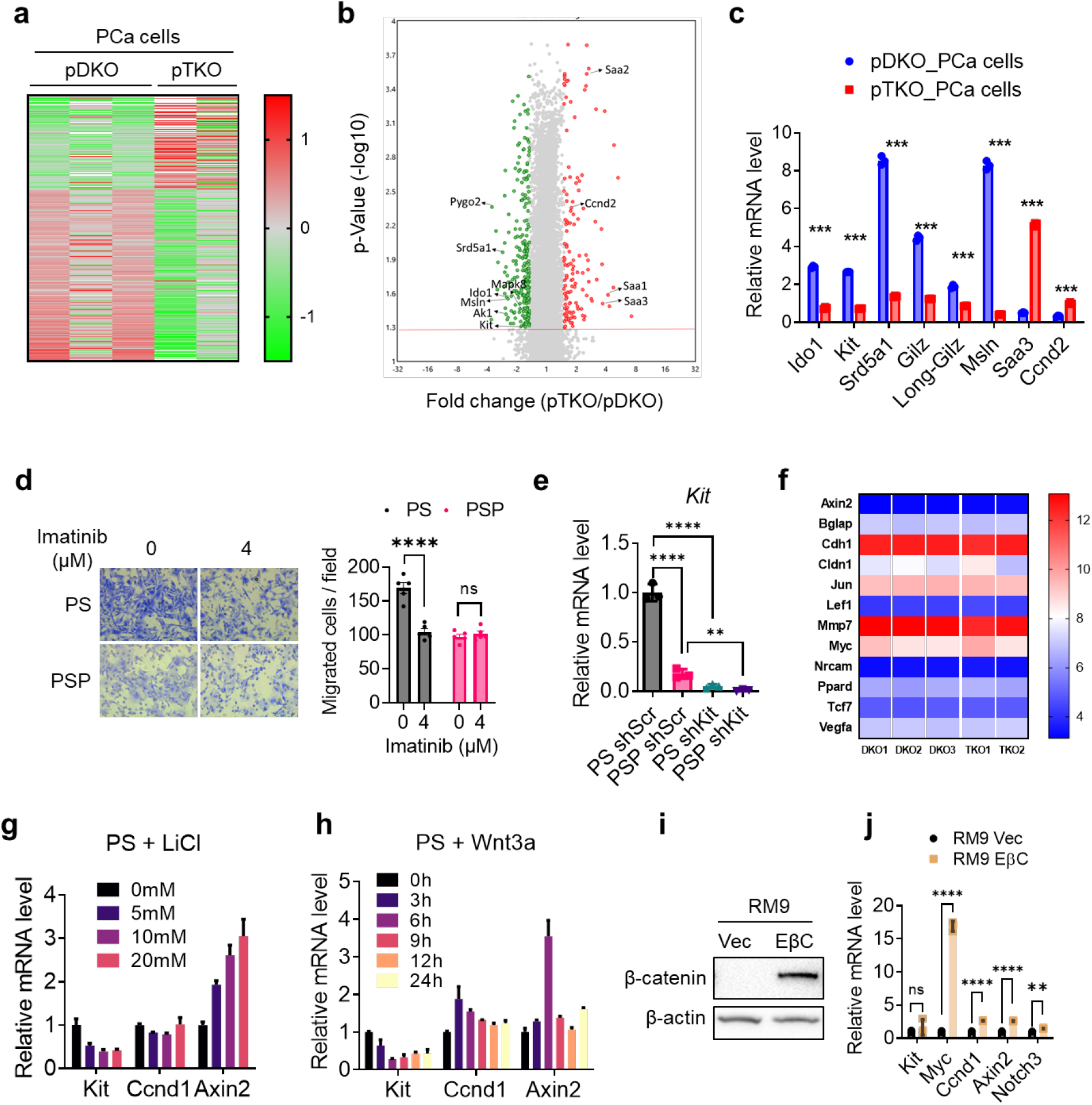
Pygo2 upregulates kit independently on Wnt signaling. **(a)** Heatmap of 379 differentially expressed gene probes between pDKO and pTKO microbead-sorted tumor cells (P<0.05, fold change >1.5). **(b)** Volcano plot showing the differential gene expression. Red line indicates the P value of 0.05. **(c)** qRT-PCR validating a short list of DE genes using PCa cells purified from pDKO and pTKO tumors, independently from the samples used in the microarray (n=3). **(d)** Representative images and quantification of transwell assay to measure the migration of PS and PSP cells treated with vehicle or imatinib (n=5). **(e)** qRT-PCR validating the shRNA-knockdown of Kit in PS and PSP (n=3). **(f)** Heatmap of Wnt downstream targets expressed in pDKO and pTKO tumor cells based on microarray data. **(g-h)** qRT-PCR to measure expression of Kit and Wnt downstream targets Ccnd1 and Axin2 in PS cells treated with LiCl or Wnt3a conditioned medium (n=3). **(i)** Western blot validating the overexpression of EβC in RM9. **(j)** qRT-PCR detecting the effect of EβC on the expression of *Kit* and Wnt targets (*Myc, Ccnd1, Axin2, Notch3*) in RM9 (n=3). In (c)(d)(e)(f)(h)(j), error bars represent SEM; ns, no*t* significant; **P<0.01, ***P<0.001, ****P<0.0001, Student’s t-test.

**Extended Data Fig. 4.**
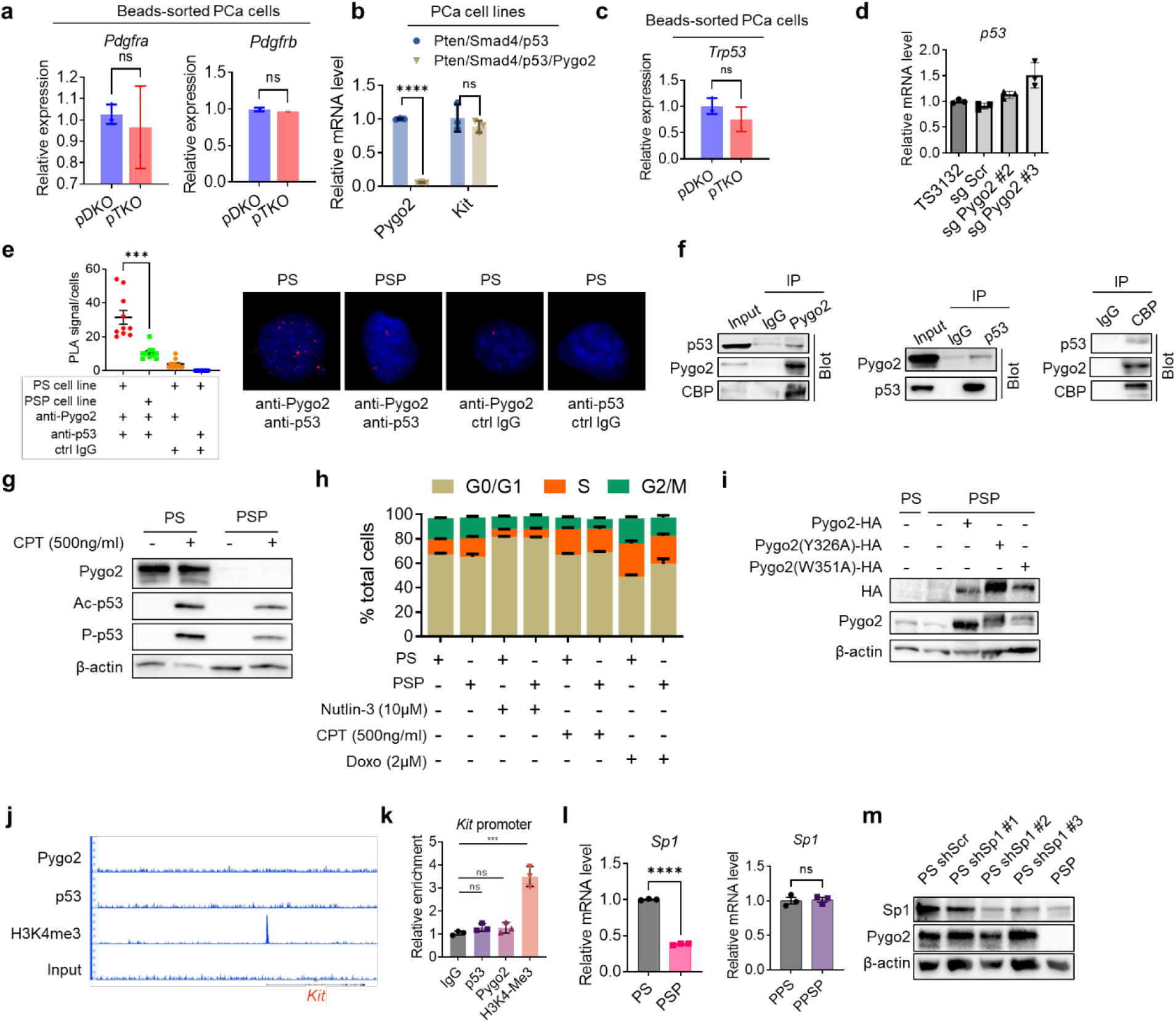
Pygo2 cooperates with p53 to upregulate the Sp1/Kit axis. **(a)** *Pdgfra* and *Pdgfrb* expression in pDKO (n=3) and pTKO (n=2) PCa cells based on normalized microarray data. **(b)** qRT-PCR for *Pygo2* and *Kit* expression in murine PCa cell lines PPS (n=3) and PPSP (n=3) established from Pten/Smad4/p53 and Pten/Smad4/p53/Pygo2 tumors, respectively. **(c)** *p53* expression in pDKO (n=3) and pTKO (n=2) PCa cells based on normalized microarray data. qRT-PCR for *p53* expression in TS3132 and its sublines (sgScr, sgPygo2) (n=3). **(e)** PLA assay to assess the proximity of Pygo2 and p53 in PS and PSP cell lines, shown in quantification plots and representative images (n=10). **(f)** co-IP/immunoblot to detect protein-protein interactions between Pygo2, p53 and p300/CBP. **(g)** Western blot to detect Pygo2, acetyl-p53 (Lys382), phospho-p53 (Ser15) in PS and PSP cell lines treated with vehicle or CPT. **(h)** Cell cycle analysis by flow cytometry for PS and PSP cell lines treated with DMSO (control), nutlin-3, CPT or doxorubicin for 24h. **(i)** Western blot confirming the overexpression of HA-tagged WT, Y326A or W351A mutant Pygo2 in PSP cell line. **(j)** IGB genomic views showing the lack of association of Pygo2 or p53 to the *Kit* promoter region, based on CUT&RUN-seq of PS cell line. **(k)** CUT&RUN-qPCR to show the lack of association of Pygo2 or p53 to the *Kit* promoter region. IgG was the negative control (n=3). (**l**) qRT-PCR to detect *Sp1* expression between PS and PSP cell lines, and between Pten/Smad4/p53 (PPS) and Pten/Smad4/p53/Pygo2 (PPSP) cell lines (n=3). **(m)** Western blot validating the shRNA knockdown of Sp1 in PS cell line. In (a)(b)(c)(d)(e)(k)(l)(m), error bars represent SEM; ns, not significant, ***P<0.001, ****P<0.0001, Student’s t-test.

**Extended Data Fig. 5.**
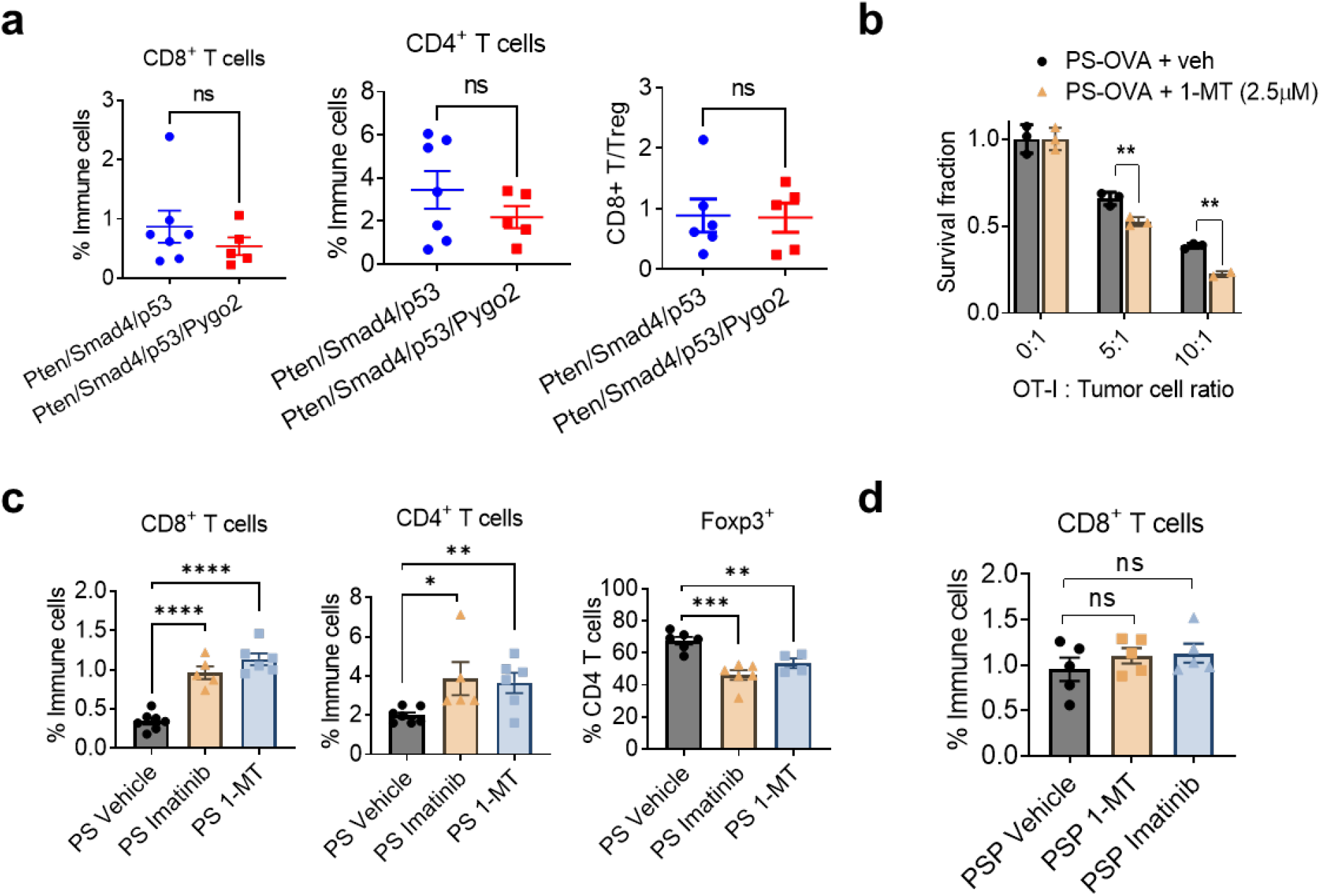
Pygo2 downregulates T cell infiltration through the Kit-Ido1 pathway. **(a)** Flow cytometry quantification of CD8^+^, CD4^+^, and CD8^+^-T/T_reg_ ratio in fully developed prostate tumors of Pten/Smad4/p53 and Pten/Smad4/p53/Pygo2 mice (n = 5 -7). **(b)** T cell cytotoxicity assay to compare the killing of PS-OVA in the presence of vehicle or 1-MT by antigen-stimulated OT-I T-cells at different E:T ratios (n=3). Viable cancer cells were quantified with resazurin. **(c)** Flow cytometry quantification of tumor-infiltrating CD8^+^, CD4^+^, and Foxp3^+^ T_reg_ cells in PS syngeneic tumors (n = 4 -7) treated with vehicle, imatinib (50mg/kg, twice/daily), or 1-MT (400mg/kg, twice/daily). **(d)** Flow cytometry quantification of tumor-infiltrating CD8^+^ T cells in PSP syngeneic tumors (n=5) treated with vehicle, imatinib (50mg/kg, twice/daily), or 1-MT (400mg/kg, twice/daily). In all panels, error bars represent SEM; ns, not significant; *P<0.05, **P<0.01, ***P<0.001, ****P<0.0001, Student’s t-test.

**Extended Data Fig. 6.**
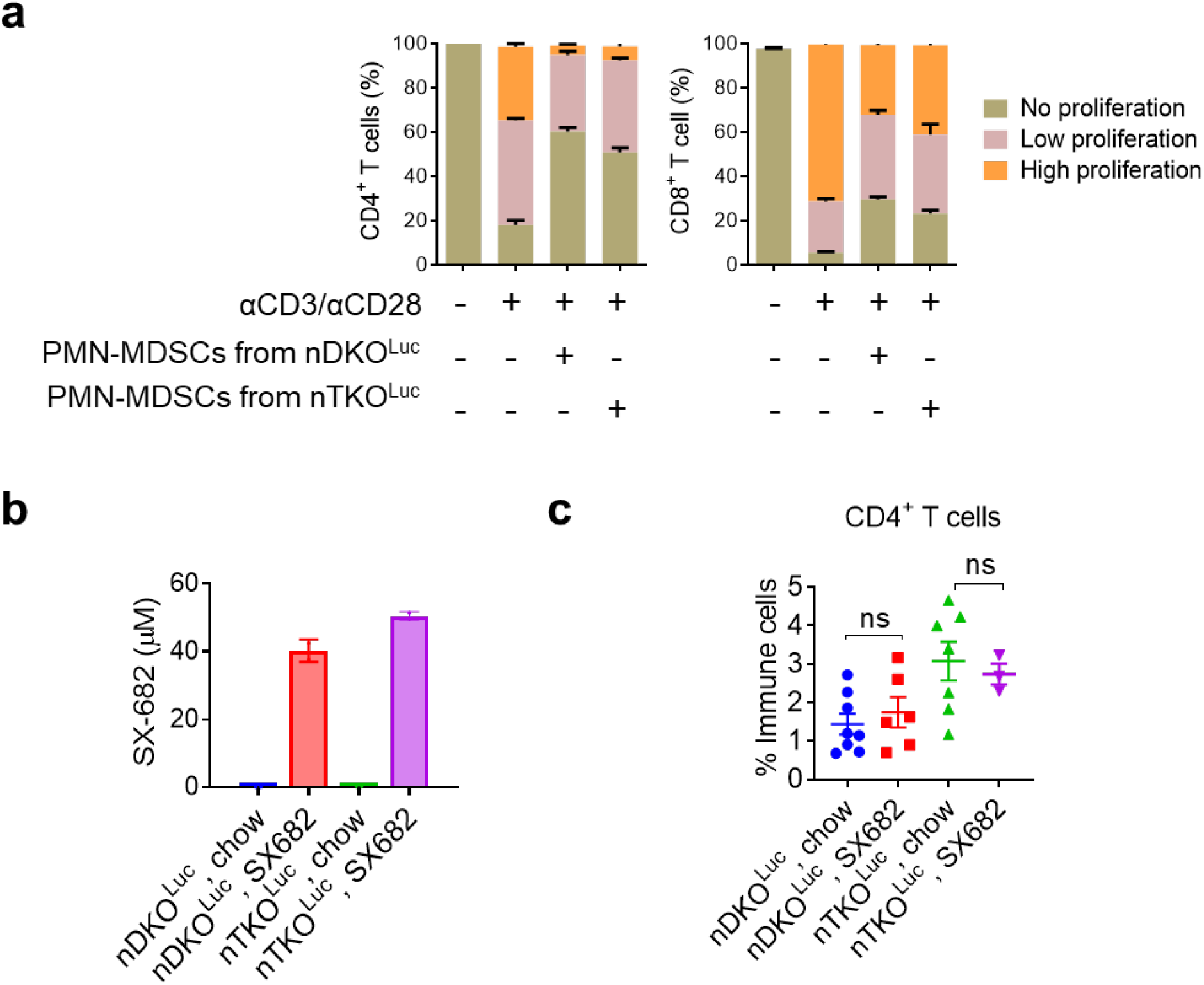
Deletion of Pygo2 enhances efficacy of CXCR2 inhibitor. **(a)** CD4^+^ (left) and CD8^+^ (right) T cell proliferation assay with and without anti-CD3/anti-CD28 stimulation and co-cultured 1:2 with PMN-MDSCs isolated from nDKO^Luc^ or nTKO^Luc^ tumors. High, moderate, and no proliferation was defined as T-cell division ≥ 2, 1, and 0, respectively based on CFSE peaks (n = 3). **(b)** SX-682 concentration measured with HPLC in the plasma of mice treated with standard or SX-682 diet for one month. **(c)** Flow cytometry quantification of CD4^+^ T cells in tumors of indicated groups. In all panels, error bars represent SEM; ns, not significant, Student’s t-test.

**Extended Data Fig. 7.**
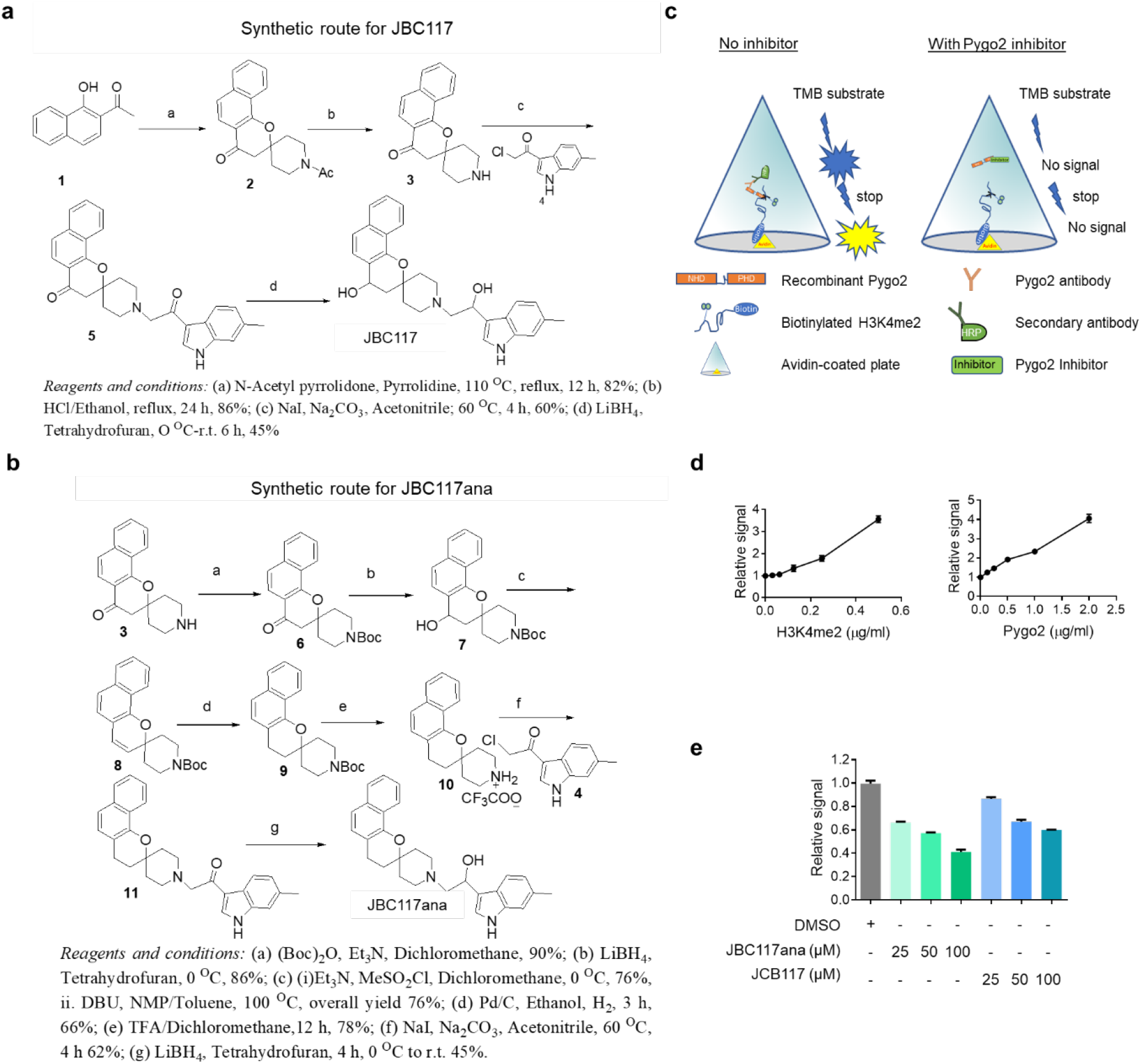
Pygo2 inhibitors antagonize PCa progression and enhance immunotherapy. **(a)** Synthetic route for JBC117. Synthesis was started from commercially available 1-Hydroxy-2-Acetonaphthone (**1**). At first 1-Hydroxy-2-Acetonaphthone was treated with 1-Acetyl-4-piperidinone at refluxing condition to get the tetra cyclic intermediate (**2**). Then acid-catalyzed acetyl group deprotection produced the corresponding amine (**3**), the amine was then coupled with 2-Chloro-1-(6-methyl-1H-indol-3-yl)-ethanone in presence of Na_2_CO_3_ to get the intermediate (**5**), The latter LiBH_4_ reduction of the coupling product (**5**) generated the desired final product JBC117. **(b)** Synthetic route for JBC117ana. For the synthesis, the tetracyclic amine (**3**) was protected by (Boc)_2_O, followed by LiBH_4_ reduction generated the alcohol (**7**), the alkene tetracyclic intermediate (**8**) was synthesized from the alcohol (**7**) via the following sequence of steps: mesylation (NEt_3_, MsCl), elimination at higher temperature (DBU, NMP). The intermediate (**8**) was treated with Pd/C under hydrogen atmosphere to get the *boc*-protected saturated intermediate (**9**). Then compound (**9**) was deprotected with TFA and the free amine was obtained *in situ* in presence of excess Na_2_CO_3_ in the reaction mixture. Similarly, the free amine (generated *in situ* in the reaction mixture) was coupled with 2-Chloro-1-(6-methyl-1H-indol-3-yl)-ethanone (**4**) to generate the intermediate (**11**). Then LiBH_4_ mediated reduction was done to get the final product JBC117ana. **(c)**The schematic of the ELISA assay. **(d)** ELISA signals increasing proportionally to the concentrations of H3K4me2 (left) or Pygo2 (right) in the absence of inhibitors. **(e)** Inhibition of Pygo2-H3K4me2 interaction by JBC117 and JBC117ana in a dose-dependent manner as measured by ELISA. In (d) and (e), error bars represent SEM.

