## Supplemental information for "Targeting Chromatin Effector Pygo2 to Enhance Immunotherapy in Prostate Cancer"

### Supplementary Table 1. Differential expression gene list bewteen purified pDKO and pTKO tumor cells (P<0.05)

| ID | TKO Ave (l | DKO Ave ( | Fold Chan | P-val | Gene Symbol | Description |
|---|---|---|---|---|---|---|
| 1419681_a_at | 5.67 | 4.33 | 2.53 | 0.0002 | Prok2 | prokineticin 2 |
| 1430602_at | 6.29 | 5.57 | 1.64 | 0.0002 | | |
| 1416122_at | 8.68 | 7.27 | 2.65 | 0.0003 | Ccnd2 | cyclin D2 |
| 1449326_x_at | 5.52 | 4.18 | 2.53 | 0.0003 | Saa2 | serum amyloid A 2 |
| 1420331_at | 4.78 | 4.05 | 1.65 | 0.0003 | Clec4e | C-type lectin domain family 4, member e |
| 1437188_at | 5.81 | 5.14 | 1.59 | 0.0003 | Gabbr1 | gamma-aminobutyric acid (GABA) B receptor, 1 |
| 1421638_at | 6.02 | 5.4 | 1.53 | 0.0003 | Psg17 | pregnancy specific glycoprotein 17 |
| 1424958_at | 5.48 | 4.87 | 1.52 | 0.0003 | Car8 | carbonic anhydrase 8 |
| 1456333_a_at | 5.37 | 4.77 | 1.51 | 0.0003 | Arhgap6 | Rho GTPase activating protein 6 |
| 1415893_at | 4.77 | 5.37 | -1.52 | 0.0003 | Sgpl1 | sphingosine phosphate lyase 1 |
| 1419740_at | 6.17 | 4.84 | 2.51 | 0.0004 | Pde6b | phosphodiesterase 6B, cGMP, rod receptor, beta polypeptide |
| 1441863_x_at | 4.28 | 3 | 2.43 | 0.0004 | Krt13 | keratin 13 |
| 1448737_at | 7.81 | 7.2 | 1.53 | 0.0004 | Tspan7 | tetraspanin 7 |
| 1447460_at | 6.35 | 5.75 | 1.51 | 0.0004 | | |
| 1438658_a_at | 5.67 | 4.85 | 1.76 | 0.0005 | S1pr3 | sphingosine-1-phosphate receptor 3 |
| 1449227_at | 4.78 | 4.12 | 1.57 | 0.0005 | Ch25h | cholesterol 25-hydroxylase |
| 1420119_s_at | 4.97 | 4.38 | 1.51 | 0.0005 | Phf3 | PHD finger protein 3 |
| 1430185_at | 4.31 | 5.07 | -1.69 | 0.0005 | | |
| 1450295_s_at | 5.5 | 6.46 | -1.94 | 0.0005 | Pvr | poliovirus receptor |
| 1416250_at | 6.6 | 7.74 | -2.2 | 0.0005 | Btg2 | B cell translocation gene 2, anti-proliferative |
| 1435588_at | 8 | 6.05 | 3.86 | 0.0006 | Wdfy1 | WD repeat and FYVE domain containing 1 |
| 1437358_at | 9.26 | 7.6 | 3.18 | 0.0006 | Wdfy1 | WD repeat and FYVE domain containing 1 |
| 1436714_at | 9.24 | 9.88 | -1.56 | 0.0006 | Lpp | LIM domain containing preferred translocation partner in lipoma |
| 1437543_at | 6.15 | 6.86 | -1.63 | 0.0006 | Fubp1 | far upstream element (FUSE) binding protein 1 |
| 1451537_at | 5.82 | 4.87 | 1.93 | 0.0007 | Chil1 | chitinase-like 1 |
| 1424907_a_at | 7.3 | 7.93 | -1.55 | 0.0007 | Farsa | phenylalanyl-tRNA synthetase, alpha subunit |
| 1425701_a_at | 5.44 | 6.35 | -1.88 | 0.0007 | Rgs3 | regulator of G-protein signaling 3 |
| 1450398_at | 5.34 | 6.42 | -2.11 | 0.0008 | Kcnk5 | potassium channel, subfamily K, member 5 |
| 1453201_at | 6.02 | 5.35 | 1.59 | 0.0009 | Rassf10 | Ras association (RalGDS/AF-6) domain family (N-terminal) member 10 |
| 1426720_at | 5.3 | 5.98 | -1.6 | 0.0009 | Apbb2 | amyloid beta (A4) precursor protein-binding, family B, member 2 |
| 1420841_at | 7.92 | 9.01 | -2.13 | 0.0009 | Ptprf | protein tyrosine phosphatase, receptor type, F |
| 1425249_a_at | 7.12 | 7.77 | -1.57 | 0.0011 | Tyro3 | TYRO3 protein tyrosine kinase 3 |
| 1421143_at | 6.8 | 7.54 | -1.67 | 0.0011 | Diaph1 | diaphanous related formin 1 |
| 1453164_a_at | 6.01 | 6.75 | -1.67 | 0.0011 | Ptdss2 | phosphatidylserine synthase 2 |
| 1422481_at | 8.29 | 6.03 | 4.8 | 0.0012 | Krt1 | keratin 1 |
| 1418014_a_at | 10.1 | 10.69 | -1.51 | 0.0012 | B4galt1 | UDP-Gal:betaGlcNAc beta 1,4- galactosyltransferase, polypeptide 1 |
| 1455026_at | 6.27 | 7.19 | -1.9 | 0.0012 | Sbno1 | sno, strawberry notch homolog 1 (Drosophila) |
| 1420836_at | 8.11 | 9.08 | -1.96 | 0.0012 | Slc25a30 | solute carrier family 25, member 30 |
| 1451403_at | 8.58 | 9.16 | -1.5 | 0.0013 | Ppp1r37 | protein phosphatase 1, regulatory subunit 37 |
| 1426542_at | 7.18 | 7.96 | -1.71 | 0.0013 | Endod1 | endonuclease domain containing 1 |
| 1435965_at | 7.34 | 7.92 | -1.5 | 0.0014 | Cnot3 | CCR4-NOT transcription complex, subunit 3 |
| 1417701_at | 5.6 | 6.39 | -1.73 | 0.0014 | Ppp1r14c | protein phosphatase 1, regulatory (inhibitor) subunit 14c |
| 1448161_a_at | 7.99 | 8.59 | -1.52 | 0.0015 | Clcn4 | chloride channel, voltage-sensitive 4 |
| 1456610_at | 6.96 | 7.58 | -1.53 | 0.0015 | Kdm6b | KDM1 lysine (K)-specific demethylase 6B |
| 1424357_at | 8.3 | 9.23 | -1.9 | 0.0016 | Tmem45b | transmembrane protein 45b |
| 1429574_at | 5.73 | 6.38 | -1.57 | 0.0018 | Clic3 | chloride intracellular channel 3 |
| 1419098_at | 4.95 | 5.61 | -1.58 | 0.0018 | Stom | stomatin |
| 1431892_a_at | 7.61 | 8.57 | -1.95 | 0.0018 | Plcd3 | phospholipase C, delta 3 |
| 1433459_x_at | 6.37 | 5.62 | 1.68 | 0.0019 | Prss2 | protease, serine 2 |
| 1430717_at | 7.49 | 8.39 | -1.86 | 0.002 | Fam83e | family with sequence similarity 83, member E |
| 1425281_a_at | 7.53 | 8.57 | -2.06 | 0.002 | Tsc22d3 | TSC22 domain family, member 3 |
| 1423946_at | 8.42 | 9.09 | -1.59 | 0.0021 | Pdlim2 | PDZ and LIM domain 2 |
| 1424609_a_at | 7.71 | 8.45 | -1.66 | 0.0021 | Cwc22; Xdh | CWC22 spliceosome-associated protein homolog (S. cerevisiae); xanthine dehydrogenase |
| 1454649_at | 8.74 | 9.7 | -1.93 | 0.0021 | Srd5a1 | steroid 5 alpha-reductase 1 |
| 1448231_at | 7.77 | 9.16 | -2.61 | 0.0021 | Fkbp5 | FK506 binding protein 5 |
| 1426541_a_at | 7.71 | 8.48 | -1.71 | 0.0022 | Endod1 | endonuclease domain containing 1 |
| 1438148_at | 7.98 | 6.79 | 2.28 | 0.0023 | Cxcl3 | chemokine (C-X-C motif) ligand 3 |
| 1436746_at | 7.77 | 8.37 | -1.51 | 0.0023 | Wnk1 | WNK lysine deficient protein kinase 1 |
| 1417561_at | 9.22 | 6.82 | 5.27 | 0.0024 | Apoc1 | apolipoprotein C-I |
| 1452239_at | 9.11 | 8.09 | 2.04 | 0.0024 | Gt(ROSA)26Sor | gene trap ROSA 26, Philippe Soriano |
| 1456391_at | 4.73 | 3.94 | 1.72 | 0.0024 | Tdrd5 | tudor domain containing 5 |
| 1421040_a_at | 3.63 | 3.01 | 1.53 | 0.0024 | Gsta2 | glutathione S-transferase, alpha 2 (Yc2) |
| 1436080_at | 6.97 | 7.64 | -1.59 | 0.0024 | | |
| 1457718_at | 6.08 | 6.83 | -1.69 | 0.0024 | Srcap; Tmem265 | Snf2-related CREBBP activator protein; transmembrane protein 265 |
| 1440920_at | 6.97 | 7.62 | -1.57 | 0.0025 | Mmp14 | matrix metallopeptidase 14 (membrane-inserted) |
| 1442439_at | 5.98 | 6.67 | -1.62 | 0.0025 | W91776 | expressed sequence W91776 |
| 1428409_at | 9.58 | 10.56 | -1.98 | 0.0025 | Naa50 | N(alpha)-acetyltransferase 50, NatE catalytic subunit |
| 1437052_s_at | 6.6 | 5.49 | 2.15 | 0.0026 | Slc2a3 | solute carrier family 2 (facilitated glucose transporter), member 3 |
| 1419697_at | 7.61 | 6.62 | 1.99 | 0.0027 | Cxcl11 | chemokine (C-X-C motif) ligand 11 |
| 1451739_at | 6.56 | 7.39 | -1.77 | 0.0027 | Klf5 | Kruppel-like factor 5 |
| 1452097_a_at | 5.95 | 6.58 | -1.55 | 0.0029 | Dusp7 | dual specificity phosphatase 7 |
| 1431211_s_at | 5.98 | 4.9 | 2.12 | 0.0031 | Them5 | thioesterase superfamily member 5 |
| 1416345_at | 8.54 | 7.95 | 1.5 | 0.0031 | Timm8a1 | translocase of inner mitochondrial membrane 8A1 |
| 1419157_at | 4.58 | 5.68 | -2.14 | 0.0031 | Sox4 | SRY (sex determining region Y)-box 4 |
| 1423912_at | 8.04 | 8.65 | -1.52 | 0.0032 | Aspscr1 | alveolar soft part sarcoma chromosome region, candidate 1 (human) |
| 1425233_at | 7.54 | 6.87 | 1.59 | 0.0034 | 2210407C18Rik | RIKEN cDNA 2210407C18 gene |
| 1423071_x_at | 9.19 | 9.9 | -1.64 | 0.0034 | 6720475J19Rik | RIKEN cDNA 6720475J19 gene |
| 1425530_a_at | 6.07 | 6.88 | -1.75 | 0.0034 | Stx3 | syntaxin 3 |
| 1416041_at | 11.65 | 13.07 | -2.66 | 0.0035 | Sgk1 | serum/glucocorticoid regulated kinase 1 |
| 1456888_at | 7.28 | 8.01 | -1.66 | 0.0036 | Pfkfb4 | 6-phosphofructo-2-kinase/fructose-2,6-biphosphatase 4 |
| 1421142_s_at | 6.61 | 7.23 | -1.54 | 0.0037 | Foxp1 | forkhead box P1 |
| 1416501_at | 6.06 | 6.84 | -1.71 | 0.0037 | Pdpk1 | 3-phosphoinositide dependent protein kinase 1 |
| 1455144_s_at | 5.99 | 5.31 | 1.6 | 0.0038 | Arhgap44 | Rho GTPase activating protein 44 |
| 1436858_at | 6.94 | 7.78 | -1.79 | 0.0038 | Mbnl2 | muscleblind-like 2 |
| 1417752_at | 6.1 | 6.96 | -1.81 | 0.0038 | Coro1c | coronin, actin binding protein 1C |
| 1429285_at | 5.07 | 4.2 | 1.83 | 0.0039 | Serpina9 | serine (or cysteine) peptidase inhibitor, clade A (alpha-1 antiproteinase, antitrypsin), member 9 |
| 1438299_at | 5.37 | 5.96 | -1.51 | 0.004 | Gm36348; Gm39 | predicted gene, 36348; predicted gene, 39564 |
| 1421313_s_at | 8.67 | 9.36 | -1.61 | 0.004 | Cttn | cortactin |
| 1448229_s_at | 7.08 | 6.26 | 1.77 | 0.0044 | Ccnd2 | cyclin D2 |
| 1419415_a_at | 7.9 | 8.64 | -1.67 | 0.0044 | Rarg | retinoic acid receptor, gamma |
| 1428249_at | 6.82 | 8.65 | -3.55 | 0.0044 | Pygo2 | pygopus 2 |
| 1425137_a_at | 7.08 | 5.85 | 2.34 | 0.0045 | H2-Q10 | histocompatibility 2, Q region locus 10 |
| 1416523_at | 7.02 | 6.21 | 1.75 | 0.0047 | Rnase1 | ribonuclease, RNase A family, 1 (pancreatic) |
| 1442017_at | 3.59 | 4.23 | -1.55 | 0.0047 | Nfs1 | nitrogen fixation gene 1 (S. cerevisiae) |
| 1437433_at | 3.51 | 2.62 | 1.85 | 0.0051 | B3galt2 | UDP-Gal:betaGlcNAc beta 1,3-galactosyltransferase, polypeptide 2 |
| 1450185_a_at | 5.31 | 6.6 | -2.43 | 0.0051 | Kcnj15 | potassium inwardly-rectifying channel, subfamily J, member 15 |
| 1416846_a_at | 5.03 | 3.99 | 2.06 | 0.0056 | Pdzrn3 | PDZ domain containing RING finger 3 |
| 1442580_at | 3.99 | 3.37 | 1.54 | 0.0056 | A530020G20Rik | RIKEN cDNA A530020G20 gene |
| 1422540_at | 5.32 | 4.66 | 1.59 | 0.0057 | Fbln1 | fibulin 1 |
| 1448328_at | 7.73 | 8.4 | -1.59 | 0.0057 | Sh3bp2 | SH3-domain binding protein 2 |
| 1420771_at | 8.02 | 8.77 | -1.68 | 0.0058 | Sprr2d | small proline-rich protein 2D |
| 1450368_a_at | 4.42 | 5.43 | -2.01 | 0.0058 | Ppp3r1; Wdr92 | protein phosphatase 3, regulatory subunit B, alpha isoform (calcineurin B, type I); WD repeat domain 92 |
| 1424749_at | 7.83 | 6.19 | 3.13 | 0.0059 | Wdfy1 | WD repeat and FYVE domain containing 1 |
| 1458662_at | 4.71 | 3.91 | 1.74 | 0.006 | 1700008O03Rik | RIKEN cDNA 1700008O03 gene |
| 1427837_at | 4.12 | 3.39 | 1.66 | 0.006 | Igkv15-103 | immunoglobulin kappa chain variable 15-103 |
| 1450808_at | 5.46 | 4.54 | 1.89 | 0.0062 | Fpr1 | formyl peptide receptor 1 |
| 1426112_a_at | 6.22 | 5.58 | 1.56 | 0.0062 | Cd72 | CD72 antigen |
| 1431464_a_at | 5.84 | 6.46 | -1.54 | 0.0062 | Pmm2 | phosphomannomutase 2 |
| 1448868_at | 8.91 | 9.52 | -1.53 | 0.0066 | Scand1 | SCAN domain-containing 1 |
| 1419709_at | 6.47 | 4.63 | 3.6 | 0.0068 | Stfa3 | stefin A3 |
| 1454686_at | 6.93 | 7.82 | -1.86 | 0.0068 | 6430706D22Rik; A | RIKEN cDNA 6430706D22 gene; RIKEN cDNA A730008H23 gene; Holliday junction recognition protein |
| 1456189_x_at | 4.64 | 5.68 | -2.06 | 0.0068 | Ltbp3 | latent transforming growth factor beta binding protein 3 |
| 1433722_at | 9.71 | 10.38 | -1.59 | 0.0069 | Akap13 | A kinase (PRKA) anchor protein 13 |
| 1425779_a_at | 5.05 | 4.37 | 1.6 | 0.0071 | Tbx1 | T-box 1 |
| 1416191_at | 7.8 | 9.5 | -3.25 | 0.0071 | Sec61a1 | Sec61 alpha 1 subunit (S. cerevisiae) |
| 1422341_s_at | 4.39 | 5.68 | -2.44 | 0.0073 | Pla2g15 | phospholipase A2, group XV |
| 1427440_a_at | 5.05 | 5.86 | -1.75 | 0.0076 | Afm | afamin |
| 1419591_at | 10 | 10.67 | -1.59 | 0.0077 | Gsdmc | gasdermin C |
| 1431382_a_at | 5.67 | 6.65 | -1.97 | 0.0079 | Clip4 | CAP-GLY domain containing linker protein family, member 4 |
| 1425273_s_at | 7.14 | 8 | -1.82 | 0.008 | Emp2 | epithelial membrane protein 2 |
| 1448538_a_at | 9.58 | 10.62 | -2.06 | 0.0081 | Rsrp1 | arginine/serine rich protein 1 |
| 1424607_a_at | 7.69 | 8.43 | -1.67 | 0.0087 | Cwc22; Xdh | CWC22 spliceosome-associated protein homolog (S. cerevisiae); xanthine dehydrogenase |
| 1420772_a_at | 9.03 | 9.62 | -1.5 | 0.0089 | Tsc22d3 | TSC22 domain family, member 3 |
| 1421786_at | 4.97 | 5.78 | -1.75 | 0.0089 | Ppp3r1 | protein phosphatase 3, regulatory subunit B, alpha isoform (calcineurin B, type I) |
| 1449554_at | 5.98 | 7.38 | -2.66 | 0.009 | Tle3 | transducin-like enhancer of split 3, homolog of Drosophila E(spl) |
| 1419099_x_at | 5.39 | 6.18 | -1.73 | 0.0091 | Stom | stomatin |
| 1436544_at | 6.62 | 6.02 | 1.52 | 0.0092 | Atp10d | ATPase, class V, type 10D |
| 1436301_at | 6.58 | 7.26 | -1.6 | 0.0092 | Dstyk | dual serine/threonine and tyrosine protein kinase |
| 1428136_at | 5.39 | 4.72 | 1.6 | 0.0094 | Sfrp1 | secreted frizzled-related protein 1 |
| 1447263_at | 3.89 | 4.55 | -1.58 | 0.0095 | | |
| 1429859_a_at | 8.06 | 8.75 | -1.61 | 0.0095 | Arl2bp | ADP-ribosylation factor-like 2 binding protein |
| 1426332_a_at | 10.45 | 11.14 | -1.62 | 0.0096 | Cldn3 | claudin 3 |
| 1421324_a_at | 5.38 | 6.39 | -2.02 | 0.0097 | Akt2 | thymoma viral proto-oncogene 2 |
| 1455735_at | 6.76 | 6.08 | 1.6 | 0.0098 | Ap1s3 | adaptor-related protein complex AP-1, sigma 3 |
| 1439915_at | 3.17 | 3.86 | -1.61 | 0.0099 | | |
| 1430641_at | 11.08 | 11.99 | -1.88 | 0.0101 | Gsdmc2; Gsdmc4 | gasdermin C2; gasdermin C4 |
| 1450007_at | 7.7 | 8.34 | -1.57 | 0.0102 | Chp1 | calcineurin-like EF hand protein 1 |
| 1416125_at | 10.15 | 11.05 | -1.88 | 0.0104 | Fkbp5 | FK506 binding protein 5 |
| 1451456_at | 5.35 | 6.02 | -1.6 | 0.0105 | A730008H23Rik; | RIKEN cDNA A730008H23 gene; Holliday junction recognition protein |
| 1437165_a_at | 7.65 | 8.4 | -1.68 | 0.0105 | Pcolce | procollagen C-endopeptidase enhancer protein |
| 1445618_at | 4.27 | 4.93 | -1.58 | 0.0106 | | |
| 1460344_at | 10.19 | 9.42 | 1.7 | 0.0107 | Pbxip1 | pre B cell leukemia transcription factor interacting protein 1 |
| 1448793_a_at | 8.59 | 9.34 | -1.69 | 0.0107 | Sdc4 | syndecan 4 |
| 1438699_at | 8.08 | 9.55 | -2.77 | 0.0109 | Srd5a1 | steroid 5 alpha-reductase 1 |
| 1443260_at | 6.71 | 7.38 | -1.59 | 0.0112 | Meis1 | Meis homeobox 1 |
| 1430127_a_at | 8.19 | 6.86 | 2.52 | 0.0115 | Ccnd2 | cyclin D2 |
| 1428738_a_at | 10.61 | 11.29 | -1.61 | 0.0115 | Tmem254a; Tmer | transmembrane protein 254a; transmembrane protein 254b; transmembrane protein 254c |
| 1424130_a_at | 7.54 | 8.22 | -1.6 | 0.0116 | Ptrf | polymerase I and transcript release factor |
| 1417613_at | 5.39 | 6.35 | -1.95 | 0.0116 | Ier5 | immediate early response 5 |
| 1439341_at | 6.77 | 5.66 | 2.16 | 0.0117 | Greb1l | growth regulation by estrogen in breast cancer-like |
| 1439795_at | 6.74 | 7.47 | -1.65 | 0.0119 | Adgrg2 | adhesion G protein-coupled receptor G2 |
| 1456398_at | 6.5 | 7.27 | -1.71 | 0.0119 | | |
| 1422017_s_at | 6.43 | 7.32 | -1.86 | 0.0119 | 4833439L19Rik | RIKEN cDNA 4833439L19 gene |
| 1424096_at | 11.43 | 10.84 | 1.5 | 0.0121 | Krt5 | keratin 5 |
| 1419449_a_at | 8.91 | 9.6 | -1.61 | 0.0122 | Gnai2 | guanine nucleotide binding protein (G protein), alpha inhibiting 2 |
| 1421026_at | 4.71 | 5.51 | -1.74 | 0.0123 | Gna12 | guanine nucleotide binding protein, alpha 12 |
| 1452911_at | 5.13 | 4.51 | 1.53 | 0.0125 | Spred1 | sprouty protein with EVH-1 domain 1, related sequence |
| 1435660_at | 8.93 | 7.54 | 2.62 | 0.0127 | C130026I21Rik; G | RIKEN cDNA C130026I21 gene; predicted gene 15753; predicted pseudogene 16026; sp110 nuclear body protein-like; Sp110 nuclear body protein pseudogene; Sp110 nuclear body protein |
| 1415834_at | 9.86 | 8.96 | 1.86 | 0.0127 | Dusp6 | dual specificity phosphatase 6 |
| 1435521_at | 6.25 | 7.28 | -2.04 | 0.0129 | Msi2 | musashi RNA-binding protein 2 |
| 1416237_at | 7.88 | 8.49 | -1.53 | 0.013 | Mpzl2 | myelin protein zero-like 2 |
| 1420835_at | 7.33 | 8.45 | -2.17 | 0.013 | Slc25a30 | solute carrier family 25, member 30 |
| 1448529_at | 5.86 | 5.27 | 1.5 | 0.0131 | Thbd | thrombomodulin |
| 1416188_at | 7.45 | 8.1 | -1.57 | 0.0133 | Gm2a | GM2 ganglioside activator protein |
| 1416190_a_at | 6.74 | 8.36 | -3.09 | 0.0133 | Sec61a1 | Sec61 alpha 1 subunit (S. cerevisiae) |
| 1422603_at | 7.79 | 6.99 | 1.74 | 0.0136 | Rnase4 | ribonuclease, RNase A family 4 |
| 1431415_a_at | 10.01 | 10.74 | -1.66 | 0.0137 | Tbpl1 | TATA box binding protein-like 1 |
| 1434484_at | 6.72 | 4.85 | 3.64 | 0.0139 | Wfdc21 | WAP four-disulfide core domain 21 |
| 1457799_at | 6.51 | 7.1 | -1.51 | 0.0139 | Kcns3 | potassium voltage-gated channel, delayed-rectifier, subfamily S, member 3 |
| 1460191_at | 7.23 | 8.25 | -2.03 | 0.0139 | Ykt6 | YKT6 homolog (S. Cerevisiae) |
| 1437517_x_at | 4.05 | 3.32 | 1.66 | 0.014 | Serpinb3a | serine (or cysteine) peptidase inhibitor, clade B (ovalbumin), member 3A |
| 1456327_at | 6.31 | 5.69 | 1.54 | 0.0141 | A530020G20Rik | RIKEN cDNA A530020G20 gene |
| 1427012_at | 5.6 | 4.91 | 1.61 | 0.0144 | Lancl1 | LanC (bacterial lantibiotic synthetase component C)-like 1 |
| 1417461_at | 7.99 | 8.73 | -1.67 | 0.0144 | Cap1 | CAP, adenylate cyclase-associated protein 1 (yeast) |
| 1421687_at | 4.5 | 3.6 | 1.86 | 0.0145 | Msmb | beta-microseminoprotein |
| 1427773_a_at | 10.18 | 10.96 | -1.71 | 0.0145 | Rabac1 | Rab acceptor 1 (prenylated) |
| 1452730_at | 7.49 | 8.86 | -2.59 | 0.0147 | Rps4l | ribosomal protein S4-like |
| 1449028_at | 8.65 | 9.27 | -1.53 | 0.015 | Rhou | ras homolog gene family, member U |
| 1448214_at | 5.56 | 6.19 | -1.55 | 0.015 | Pdhb | pyruvate dehydrogenase (lipoamide) beta |
| 1453801_at | 4.57 | 3.18 | 2.62 | 0.0154 | Them5 | thioesterase superfamily member 5 |
| 1428142_at | 5.2 | 4.32 | 1.83 | 0.0154 | Etv5 | ets variant 5 |
| 1420975_at | 4.26 | 4.96 | -1.63 | 0.0156 | Baz1b | bromodomain adjacent to zinc finger domain, 1B |
| 1425789_s_at | 9.27 | 10.04 | -1.71 | 0.0156 | Anxa8 | annexin A8 |
| 1440851_at | 4.19 | 4.99 | -1.74 | 0.0157 | Erlec1 | endoplasmic reticulum lectin 1 |
| 1458089_at | 4.88 | 5.87 | -1.98 | 0.0157 | | |
| 1451021_a_at | 11.1 | 11.7 | -1.52 | 0.016 | Klf5 | Kruppel-like factor 5 |
| 1448433_a_at | 6.57 | 7.24 | -1.59 | 0.0161 | Pcolce | procollagen C-endopeptidase enhancer protein |
| 1452731_x_at | 4.42 | 3.39 | 2.04 | 0.0163 | Gm10409; Gm28 | predicted pseudogene 8348; predicted gene 2897; predicted gene 2956; alpha-takusan pseudogene; predicted gene 3095; predicted gene 3164; predicted gene 3173; predicted gene 3317; predicted gene 3373; predicted gene 3383; predicted gene, 33933; predicted gene, 3488; predicted gene 3500; predicted gene 3558; predicted gene 3636; predicted gene 3696; predicted gene 5796; predicted gene LOC100040870; alpha takusan-like; uncharacterized LOC101055754; uncharacterized LOC102638 |
| 1431214_at | 10.79 | 12.2 | -2.66 | 0.0164 | Gm3579; Gm405 | predicted gene 3579; predicted gene, 40514; predicted gene, 40991; predicted gene, 42035 |
| 1448640_at | 5.77 | 4.55 | 2.32 | 0.0165 | Slc14a1 | solute carrier family 14 (urea transporter), member 1 |
| 1417496_at | 9.2 | 9.79 | -1.5 | 0.0166 | Cp | ceruloplasmin |
| 1455577_at | 9.31 | 8.43 | 1.84 | 0.0168 | Ccl28 | chemokine (C-C motif) ligand 28 |
| 1458512_at | 5.71 | 6.33 | -1.54 | 0.0173 | Tle3 | transducin-like enhancer of split 3, homolog of Drosophila E(spl) |
| 1435913_at | 5.52 | 4.85 | 1.59 | 0.0174 | B4galnt4 | beta-1,4-N-acetyl-galactosaminyl transferase 4 |
| 1422959_s_at | 8.35 | 9.67 | -2.51 | 0.0174 | Rnf114 | ring finger protein 114 |
| 1436967_at | 6.15 | 6.76 | -1.52 | 0.0177 | Ankrd11 | ankyrin repeat domain 11 |
| 1454883_at | 11.04 | 11.77 | -1.66 | 0.0179 | Gsdmc2 | gasdermin C2 |
| 1450018_s_at | 7.28 | 8.26 | -1.97 | 0.018 | Slc25a30 | solute carrier family 25, member 30 |
| 1450716_at | 7.66 | 8.26 | -1.52 | 0.0182 | Adamts1 | a disintegrin-like and metallopeptidase (reprolysin type) with thrombospondin type 1 motif, 1 |
| 1456539_at | 4.69 | 4.01 | 1.6 | 0.0183 | Klk13 | kallikrein related-peptidase 13 |
| 1431818_at | 7.19 | 8.26 | -2.1 | 0.0183 | Rab10os | RAB10, member RAS oncogene family, opposite strand |
| 1451956_a_at | 6.85 | 7.48 | -1.55 | 0.0184 | Sigmar1 | sigma non-opioid intracellular receptor 1 |
| 1426313_at | 8.58 | 9.25 | -1.6 | 0.0186 | Bre | brain and reproductive organ-expressed protein |
| 1437938_x_at | 6.48 | 7.18 | -1.62 | 0.0188 | Dnm2 | dynamin 2 |
| 1436932_at | 5.72 | 4.04 | 3.21 | 0.0189 | Grhl3 | grainyhead-like 3 (Drosophila) |
| 1450834_at | 4.79 | 5.47 | -1.6 | 0.0189 | Fut4 | fucosyltransferase 4 |
| 1428720_s_at | 5.29 | 4.43 | 1.82 | 0.019 | Iglc2; Iglc3; Iglv1 | immunoglobulin lambda constant 2; immunoglobulin lambda constant 3; immunoglobulin lambda variable 1 |
| 1449452_a_at | 7.72 | 8.87 | -2.22 | 0.019 | Gp2 | glycoprotein 2 (zymogen granule membrane) |
| 1433582_at | 7.47 | 6.64 | 1.78 | 0.0191 | 1190002N15Rik | RIKEN cDNA 1190002N15 gene |
| 1428834_at | 8.19 | 7.57 | 1.53 | 0.0192 | | |
| 1425811_a_at | 9.36 | 10.15 | -1.73 | 0.0194 | Csrp1 | cysteine and glycine-rich protein 1 |
| 1435792_at | 10.7 | 9.3 | 2.64 | 0.0196 | Csprs; Gm15433; | component of Sp100-rs; predicted pseudogene 15433; predicted gene 2666; predicted pseudogene 7609; proteinase-activated receptor 1-like |
| 1421140_a_at | 7.4 | 7.99 | -1.51 | 0.0197 | Foxp1 | forkhead box P1 |
| 1437073_x_at | 9.66 | 10.28 | -1.55 | 0.0199 | | |
| 1416629_at | 9.27 | 10.02 | -1.68 | 0.0199 | Slc1a5 | solute carrier family 1 (neutral amino acid transporter), member 5 |
| 1456080_a_at | 10.85 | 11.59 | -1.67 | 0.02 | Serinc3 | serine incorporator 3 |
| 1450378_at | 9.03 | 10.27 | -2.37 | 0.0201 | Tapbp | TAP binding protein |
| 1440055_at | 6.14 | 7.44 | -2.47 | 0.0201 | | |
| 1416123_at | 5.14 | 4.17 | 1.95 | 0.0205 | Ccnd2 | cyclin D2 |
| 1444552_at | 5.95 | 6.58 | -1.54 | 0.0205 | | |
| 1438069_a_at | 6.33 | 6.98 | -1.57 | 0.0214 | Rbm5 | RNA binding motif protein 5 |
| 1416432_at | 8.16 | 7.54 | 1.53 | 0.0215 | Pfkfb3 | 6-phosphofructo-2-kinase/fructose-2,6-biphosphatase 3 |
| 1444734_at | 4.02 | 5.17 | -2.23 | 0.0216 | LOC102632594 | uncharacterized LOC102632594 |
| 1448470_at | 4.88 | 4.04 | 1.79 | 0.0217 | Fbp1 | fructose bisphosphatase 1 |
| 1425606_at | 10.21 | 9.27 | 1.91 | 0.0219 | Slc5a8 | solute carrier family 5 (iodide transporter), member 8 |
| 1456956_at | 7.14 | 7.9 | -1.69 | 0.0219 | Ikzf2 | IKAROS family zinc finger 2 |
| 1450747_at | 6.1 | 7.52 | -2.68 | 0.0221 | Keap1 | kelch-like ECH-associated protein 1 |
| 1429968_at | 4.66 | 5.28 | -1.54 | 0.0222 | | |
| 1419075_s_at | 10.53 | 8.29 | 4.74 | 0.0223 | Saa1 | serum amyloid A 1 |
| 1437591_a_at | 9.59 | 10.19 | -1.51 | 0.0223 | Wdr1 | WD repeat domain 1 |
| 1416933_at | 8.75 | 9.44 | -1.61 | 0.0227 | Por | P450 (cytochrome) oxidoreductase |
| 1418832_at | 6.67 | 7.52 | -1.8 | 0.0229 | Pkp3 | plakophilin 3 |
| 1429413_at | 7.87 | 7.28 | 1.5 | 0.023 | Cpm | carboxypeptidase M |
| 1452092_at | 6.8 | 5.59 | 2.32 | 0.0232 | Chst15 | carbohydrate (N-acetylgalactosamine 4-sulfate 6-O) sulfotransferase 15 |
| 1444016_at | 7.04 | 8.7 | -3.17 | 0.0233 | Anxa1 | annexin A1 |
| 1443233_at | 5.04 | 5.68 | -1.56 | 0.0234 | | |
| 1423446_at | 6.45 | 7.3 | -1.8 | 0.0237 | Dapk3 | death-associated protein kinase 3 |
| 1447040_at | 4.68 | 5.78 | -2.13 | 0.0239 | | |
| 1432558_a_at | 8.01 | 9.05 | -2.05 | 0.024 | Mal | myelin and lymphocyte protein, T cell differentiation protein |
| 1440990_at | 5.61 | 6.23 | -1.53 | 0.0242 | Kif26b | kinesin family member 26B |
| 1421944_a_at | 3.63 | 2.83 | 1.74 | 0.0243 | Asgr1 | asialoglycoprotein receptor 1 |
| 1423089_at | 7.57 | 8.55 | -1.97 | 0.0245 | Tmod3 | tropomodulin 3 |
| 1420723_at | 7.3 | 6.53 | 1.71 | 0.0246 | Vnn3 | vanin 3 |
| 1417704_a_at | 3.93 | 3.22 | 1.63 | 0.0246 | Arhgap6 | Rho GTPase activating protein 6 |

| | | | | | | |
|---|---|---|---|---|---|---|
| 1418187_at | 6 | 6.95 | -1.92 | 0.0248 | Ramp2 | receptor (calcitonin) activity modifying protein 2 |
| 1419654_at | 6.88 | 7.5 | -1.54 | 0.0249 | Tle3 | transducin-like enhancer of split 3, homolog of Drosophila E(spl) |
| 1430980_a_at | 11.1 | 11.94 | -1.79 | 0.025 | Eif4a1 | eukaryotic translation initiation factor 4A1 |
| 1435308_at | 7.53 | 8.55 | -2.02 | 0.025 | Fut9 | fucosyltransferase 9 |
| 1421156_a_at | 9.23 | 8.63 | 1.51 | 0.0251 | Dsc2 | desmocollin 2 |
| 1420913_at | 8.17 | 9 | -1.77 | 0.0252 | Slco2a1 | solute carrier organic anion transporter family, member 2a1 |
| 1450407_a_at | 8.92 | 9.89 | -1.96 | 0.0254 | Anp32a | acidic (leucine-rich) nuclear phosphoprotein 32 family, member A |
| 1455843_at | 7.63 | 9.03 | -2.64 | 0.0254 | Fut4 | fucosyltransferase 4 |
| 1453367_a_at | 7.91 | 8.55 | -1.56 | 0.0255 | Abhd12 | abhydrolase domain containing 12 |
| 1450788_at | 11.22 | 9.16 | 4.18 | 0.0258 | Saa1 | serum amyloid A 1 |
| 1447706_at | 3.58 | 4.35 | -1.7 | 0.0259 | | |
| 1456689_at | 5.13 | 5.76 | -1.54 | 0.026 | Rnf10 | ring finger protein 10 |
| 1438667_at | 4.72 | 5.36 | -1.56 | 0.0262 | Sybu | syntabulin (syntaxin-interacting) |
| 1416159_at | 9.83 | 10.64 | -1.76 | 0.0262 | Nr2f2 | nuclear receptor subfamily 2, group F, member 2 |
| 1434745_at | 8.44 | 7.08 | 2.56 | 0.0263 | Ccnd2 | cyclin D2 |
| 1418210_at | 9.29 | 8.64 | 1.57 | 0.0265 | Pfn2 | profilin 2 |
| 1446720_at | 5.32 | 6.01 | -1.61 | 0.0265 | | |
| 1420768_a_at | 6.03 | 7.21 | -2.27 | 0.0265 | Dhx58 | DEXH (Asp-Glu-X-His) box polypeptide 58 |
| 1434610_at | 7.83 | 8.58 | -1.68 | 0.0268 | Plec | plectin |
| 1420437_at | 6.79 | 8.14 | -2.55 | 0.027 | Ido1 | indoleamine 2,3-dioxygenase 1 |
| 1432543_a_at | 5.94 | 6.61 | -1.6 | 0.0271 | Klf13 | Kruppel-like factor 13 |
| 1425405_a_at | 6.53 | 7.47 | -1.91 | 0.0272 | Adar | adenosine deaminase, RNA-specific |
| 1420378_at | 9.88 | 9.2 | 1.6 | 0.0274 | Sftpd | surfactant associated protein D |
| 1416572_at | 5.79 | 6.52 | -1.66 | 0.0275 | Mmp14 | matrix metallopeptidase 14 (membrane-inserted) |
| 1457233_at | 5.27 | 5.87 | -1.52 | 0.0276 | | |
| 1429834_a_at | 5.6 | 6.25 | -1.56 | 0.0277 | Vps51 | vacuolar protein sorting 51 homolog (S. cerevisiae) |
| 1455394_at | 6.89 | 7.48 | -1.5 | 0.0279 | Pias4 | protein inhibitor of activated STAT 4 |
| 1425394_at | 6.31 | 4.81 | 2.82 | 0.028 | BC023105 | cDNA sequence BC023105 |
| 1452684_at | 6.49 | 7.12 | -1.55 | 0.0281 | Akt1s1 | AKT1 substrate 1 (proline-rich) |
| 1456857_at | 7.53 | 6.88 | 1.56 | 0.0282 | 1500011B03Rik | RIKEN cDNA 1500011B03 gene |
| 1460238_at | 6.17 | 7.48 | -2.48 | 0.0282 | Msln | mesothelin |
| 1438683_at | 7.14 | 7.75 | -1.53 | 0.0283 | Wasf2 | WAS protein family, member 2 |
| 1453604_a_at | 4.85 | 5.45 | -1.51 | 0.0289 | Hbs1l | Hbs1-like (S. cerevisiae) |
| 1453605_s_at | 5.88 | 6.5 | -1.53 | 0.029 | Ccdc91 | coiled-coil domain containing 91 |
| 1435031_at | 6.16 | 6.83 | -1.59 | 0.029 | Tmem120a | transmembrane protein 120A |
| 1449833_at | 5.7 | 4.17 | 2.89 | 0.0291 | Sprr2f | small proline-rich protein 2F |
| 1439651_at | 6.67 | 7.43 | -1.69 | 0.0293 | | |
| 1420363_at | 6.75 | 7.39 | -1.56 | 0.0295 | Bik | BCL2-interacting killer |
| 1417732_at | 9.23 | 9.84 | -1.53 | 0.0296 | Anxa8 | annexin A8 |
| 1430152_at | 3.97 | 4.61 | -1.56 | 0.0297 | | |
| 1438497_at | 4.02 | 4.62 | -1.51 | 0.0299 | Mfsd8 | major facilitator superfamily domain containing 8 |
| 1424524_at | 9.64 | 10.39 | -1.69 | 0.0303 | Dram1 | DNA-damage regulated autophagy modulator 1 |
| 1418723_at | 5.78 | 4.8 | 1.98 | 0.0305 | Lpar3 | lysophosphatidic acid receptor 3 |
| 1449984_at | 10.56 | 9.77 | 1.73 | 0.0305 | Cxcl2 | chemokine (C-X-C motif) ligand 2 |
| 1417275_at | 9.23 | 9.9 | -1.59 | 0.0306 | Mal | myelin and lymphocyte protein, T cell differentiation protein |
| 1452408_at | 4.39 | 3.64 | 1.68 | 0.0307 | | |
| 1450826_a_at | 10.86 | 8.97 | 3.69 | 0.0308 | Saa3 | serum amyloid A 3 |
| 1435227_at | 6.69 | 5.94 | 1.69 | 0.0309 | Bcl11b | B cell leukemia/lymphoma 11B |
| 1426888_at | 6.34 | 7.24 | -1.87 | 0.0309 | Ehmt2 | euchromatic histone lysine N-methyltransferase 2 |
| 1455793_at | 6.77 | 6.15 | 1.54 | 0.0315 | Fam149a | family with sequence similarity 149, member A |
| 1429463_at | 5.64 | 6.68 | -2.06 | 0.0317 | Prkaa2 | protein kinase, AMP-activated, alpha 2 catalytic subunit |
| 1436265_at | 6.78 | 4.68 | 4.28 | 0.0324 | Gm10419 | predicted gene 10419 |
| 1449184_at | 11.86 | 10.83 | 2.03 | 0.0326 | Pglyrp1 | peptidoglycan recognition protein 1 |
| 1432436_a_at | 7.42 | 8.11 | -1.6 | 0.0334 | Ak3 | adenylate kinase 3 |
| 1426089_a_at | 7.87 | 8.52 | -1.56 | 0.0335 | BC003331 | cDNA sequence BC003331 |
| 1447831_s_at | 6.92 | 6.23 | 1.61 | 0.0336 | Mtmr7 | myotubularin related protein 7 |
| 1425810_a_at | 8.2 | 9.6 | -2.64 | 0.0337 | Csrp1 | cysteine and glycine-rich protein 1 |
| 1459897_a_at | 10.08 | 10.79 | -1.63 | 0.0338 | Sbsn | suprabasin |
| 1439630_x_at | 9.87 | 10.63 | -1.7 | 0.034 | Sbsn | suprabasin |
| 1439045_x_at | 6.7 | 6.09 | 1.53 | 0.0341 | Tc2n | tandem C2 domains, nuclear |
| 1422141_s_at | 8.85 | 7.68 | 2.25 | 0.0344 | Csprs; Gm15433; | component of Sp100-rs; predicted pseudogene 15433; predicted gene 2666; predicted pseudogene 7609; proteinase-activated receptor 1-like |
| 1443509_at | 5.69 | 6.59 | -1.87 | 0.0344 | | |
| 1423331_a_at | 3.99 | 2.94 | 2.06 | 0.0345 | Pvrl3 | poliovirus receptor-related 3 |
| 1460431_at | 6.33 | 6.99 | -1.58 | 0.0347 | Gcnt1 | glucosaminyl (N-acetyl) transferase 1, core 2 |
| 1457150_at | 6.79 | 7.38 | -1.5 | 0.0358 | AI428301 | expressed sequence AI428301 |
| 1454268_a_at | 10.19 | 10.8 | -1.52 | 0.0358 | Cyba | cytochrome b-245, alpha polypeptide |
| 1415798_at | 6.45 | 7.63 | -2.28 | 0.0359 | Ddr1 | discoidin domain receptor family, member 1 |
| 1437621_x_at | 9.81 | 9.15 | 1.58 | 0.036 | Phgdh | 3-phosphoglycerate dehydrogenase |
| 1428372_at | 8.96 | 9.63 | -1.59 | 0.0362 | St5 | suppression of tumorigenicity 5 |
| 1437045_at | 4.98 | 5.75 | -1.71 | 0.0362 | Mapk8 | mitogen-activated protein kinase 8 |
| 1456881_at | 5.25 | 5.94 | -1.62 | 0.0364 | Nbeal2 | neurobeachin-like 2 |
| 1415859_at | 9.29 | 9.89 | -1.51 | 0.0365 | Eif3c | eukaryotic translation initiation factor 3, subunit C |
| 1451858_at | 6.89 | 6 | 1.85 | 0.0366 | Mrgpra2a; Mrgpr | MAS-related GPR, member A2A; MAS-related GPR, member A2B |
| 1440841_at | 5.5 | 6.41 | -1.89 | 0.0366 | | |
| 1428547_at | 6.02 | 7.1 | -2.11 | 0.0366 | Nt5e | 5' nucleotidase, ecto |
| 1427034_at | 7.87 | 7.09 | 1.72 | 0.037 | Ace | angiotensin I converting enzyme (peptidyl-dipeptidase A) 1 |
| 1425645_s_at | 7.99 | 9.1 | -2.15 | 0.0374 | Cyp2b10 | cytochrome P450, family 2, subfamily b, polypeptide 10 |
| 1451344_at | 6.04 | 5.38 | 1.58 | 0.0376 | Tmem119 | transmembrane protein 119 |
| 1449366_at | 5.2 | 4.37 | 1.78 | 0.0377 | Mmp8 | matrix metallopeptidase 8 |
| 1426543_x_at | 8.09 | 8.67 | -1.5 | 0.0377 | Endod1 | endonuclease domain containing 1 |
| 1425603_at | 11.05 | 10.45 | 1.51 | 0.0381 | Tmem176a | transmembrane protein 176A |
| 1449682_s_at | 4.48 | 5.11 | -1.54 | 0.0382 | Tubb2a-ps2; Tubb | tubulin, beta 2a, pseudogene 2; tubulin, beta 2B class IIB |
| 1425711_a_at | 6.07 | 7.43 | -2.57 | 0.0385 | Akt1 | thymoma viral proto-oncogene 1 |
| 1457020_at | 5.82 | 6.43 | -1.52 | 0.0387 | | |
| 1455956_x_at | 7.79 | 6.58 | 2.31 | 0.0394 | Ccnd2 | cyclin D2 |
| 1430724_at | 5.8 | 5.18 | 1.54 | 0.0394 | Ephx3 | epoxide hydrolase 3 |
| 1416388_at | 6.82 | 7.62 | -1.74 | 0.0394 | Pip4k2c | phosphatidylinositol-5-phosphate 4-kinase, type II, gamma |
| 1429262_at | 8.68 | 8.06 | 1.53 | 0.0395 | Rassf6 | Ras association (RalGDS/AF-6) domain family member 6 |
| 1420470_at | 6.57 | 3.72 | 7.21 | 0.0397 | Sult1c1 | sulfotransferase family, cytosolic, 1C, member 1 |
| 1418186_at | 9.94 | 8.66 | 2.43 | 0.0397 | Gstt1 | glutathione S-transferase, theta 1 |
| 1455531_at | 10.91 | 10.23 | 1.6 | 0.0399 | Mfsd4 | major facilitator superfamily domain containing 4 |
| 1431230_a_at | 6.52 | 7.24 | -1.65 | 0.04 | Btbd9 | BTB (POZ) domain containing 9 |
| 1428127_at | 7.71 | 8.35 | -1.56 | 0.0405 | Atxn7l3b | ataxin 7-like 3B |
| 1451867_x_at | 7.55 | 6.66 | 1.85 | 0.0406 | Arhgap6 | Rho GTPase activating protein 6 |
| 1420867_at | 9.75 | 10.35 | -1.52 | 0.0407 | Gm21540; Tmed2 | predicted gene, 21540; transmembrane emp24 domain trafficking protein 2 |
| 1431101_a_at | 7.44 | 8.73 | -2.45 | 0.0407 | Srd5a1 | steroid 5 alpha-reductase 1 |
| 1454106_a_at | 4.1 | 4.84 | -1.67 | 0.0408 | Cxxc1 | CXXC finger 1 (PHD domain) |
| 1455898_x_at | 5.01 | 4.07 | 1.92 | 0.0409 | Slc2a3 | solute carrier family 2 (facilitated glucose transporter), member 3 |
| 1448404_at | 8.05 | 8.65 | -1.52 | 0.0413 | Scamp2 | secretory carrier membrane protein 2 |
| 1433610_at | 8.18 | 9.22 | -2.06 | 0.0414 | AA986860 | expressed sequence AA986860 |
| 1420400_at | 3.66 | 2.85 | 1.76 | 0.0416 | Sval1 | seminal vesicle antigen-like 1 |
| 1449166_at | 9.25 | 9.99 | -1.67 | 0.0417 | S100a14 | S100 calcium binding protein A14 |
| 1437303_at | 6.3 | 5.59 | 1.64 | 0.0419 | Il6st | interleukin 6 signal transducer |
| 1457700_at | 4.85 | 4.19 | 1.57 | 0.042 | Cyp4f39 | cytochrome P450, family 4, subfamily f, polypeptide 39 |
| 1424770_at | 5.55 | 6.19 | -1.55 | 0.042 | Cald1 | caldesmon 1 |
| 1419152_at | 4.68 | 6.55 | -3.64 | 0.0425 | 2810417H13Rik | RIKEN cDNA 2810417H13 gene |
| 1427430_at | 7.98 | 8.62 | -1.56 | 0.0427 | Suco | SUN domain containing ossification factor |
| 1449586_at | 10.18 | 10.84 | -1.58 | 0.0428 | Pkp1 | plakophilin 1 |
| 1429365_at | 6.15 | 7.48 | -2.51 | 0.0428 | Mroh4 | maestro heat-like repeat family member 4 |
| 1439528_at | 7.73 | 6.46 | 2.4 | 0.0429 | 4833423E24Rik | RIKEN cDNA 4833423E24 gene |
| 1423153_x_at | 5.8 | 5.07 | 1.66 | 0.0432 | Cfh | complement component factor h |
| 1421346_a_at | 6.2 | 6.84 | -1.55 | 0.0433 | Slc6a6 | solute carrier family 6 (neurotransmitter transporter, taurine), member 6 |
| 1451787_at | 7.73 | 8.61 | -1.84 | 0.0442 | Cyp2b10 | cytochrome P450, family 2, subfamily b, polypeptide 10 |
| 1421833_at | 6.24 | 5.64 | 1.51 | 0.0447 | Pip5k1b | phosphatidylinositol-4-phosphate 5-kinase, type 1 beta |
| 1426632_at | 9.15 | 9.85 | -1.62 | 0.0449 | Kctd14 | potassium channel tetramerisation domain containing 14 |
| 1418209_a_at | 9.44 | 8.58 | 1.81 | 0.0455 | Pfn2 | profilin 2 |
| 1457572_at | 3.98 | 3.17 | 1.75 | 0.0457 | | |
| 1420891_at | 5.35 | 6.06 | -1.64 | 0.0459 | Wnt7b | wingless-type MMTV integration site family, member 7B |
| 1423072_at | 7.05 | 8.17 | -2.18 | 0.0461 | | |
| 1433101_at | 6.63 | 7.23 | -1.51 | 0.0462 | | |
| 1432004_a_at | 5.55 | 6.75 | -2.3 | 0.0463 | Dnm2 | dynamin 2 |
| 1437896_at | 4.82 | 5.43 | -1.53 | 0.0467 | | |
| 1437923_at | 7.21 | 7.9 | -1.61 | 0.0467 | Csnk1a1 | casein kinase 1, alpha 1 |
| 1426301_at | 9.5 | 10.25 | -1.68 | 0.047 | Alcam | activated leukocyte cell adhesion molecule |
| 1421141_a_at | 6.56 | 7.32 | -1.69 | 0.0472 | Foxp1 | forkhead box P1 |
| 1443137_at | 4.84 | 5.42 | -1.5 | 0.048 | | |
| 1448155_at | 7.98 | 8.57 | -1.5 | 0.0481 | Pdcd6ip | programmed cell death 6 interacting protein |
| 1419481_at | 6.29 | 5.42 | 1.83 | 0.0487 | Sell | selectin, lymphocyte |
| 1436993_x_at | 8.69 | 8.06 | 1.55 | 0.0487 | Pfn2 | profilin 2 |
| 1454714_x_at | 10.14 | 9.51 | 1.55 | 0.0488 | Phgdh | 3-phosphoglycerate dehydrogenase |
| 1452081_a_at | 7.04 | 7.88 | -1.8 | 0.0488 | Misp | mitotic spindle positioning |
| 1452968_at | 5.28 | 4.65 | 1.55 | 0.0489 | Cthrc1 | collagen triple helix repeat containing 1 |
| 1415900_a_at | 3.94 | 4.67 | -1.66 | 0.0493 | Kit | kit oncogene |
| 1444064_at | 3.98 | 4.71 | -1.65 | 0.0495 | Samhd1 | SAM domain and HD domain, 1 |
| 1430629_at | 4.86 | 4.24 | 1.53 | 0.0499 | Slc16a14 | solute carrier family 16 (monocarboxylic acid transporters), member 14 |

#### Supplementary Table 2. Hallmark gene sets enriched in pDKO based on GSEA analysis

| NAME | GS<br> foll | GS DETAILS | SIZE | ES | NES | NOM p-val | FDR q-val | FWER p-va | RANK AT M | LEADING EDGE |
|---|---|---|---|---|---|---|---|---|---|---|
| HALLMARK_EPITHELIAL_MESENCHYMAL_TRANSITION | HALLMARK | Details ... | 196 | 0.348611 | 1.532583 | 0 | 0.190087 | 0.062 | 1864 | tags=29%, list=11%, signal=32% |
| HALLMARK_P53_PATHWAY | HALLMARK | Details ... | 199 | 0.303562 | 1.326126 | 0.019608 | 0.373049 | 0.24 | 2779 | tags=33%, list=17%, signal=39% |
| HALLMARK_ESTROGEN_RESPONSE_LATE | HALLMARK | Details ... | 197 | 0.302837 | 1.305205 | 0.011364 | 0.297652 | 0.288 | 2716 | tags=31%, list=17%, signal=37% |
| HALLMARK_UV_RESPONSE_DN | HALLMARK | Details ... | 143 | 0.314028 | 1.304627 | 0.006944 | 0.22426 | 0.289 | 2174 | tags=27%, list=13%, signal=31% |
| HALLMARK_ESTROGEN_RESPONSE_EARLY | HALLMARK | Details ... | 197 | 0.29619 | 1.298922 | 0 | 0.192806 | 0.304 | 2716 | tags=32%, list=17%, signal=39% |
| HALLMARK_CHOLESTEROL_HOMEOSTASIS | HALLMARK | Details ... | 73 | 0.329549 | 1.245239 | 0.114428 | 0.245502 | 0.414 | 2126 | tags=34%, list=13%, signal=39% |
| HALLMARK_MYOGENESIS | HALLMARK | Details ... | 199 | 0.285263 | 1.224582 | 0.066667 | 0.248581 | 0.465 | 1786 | tags=19%, list=11%, signal=21% |
| HALLMARK_PEROXISOME | HALLMARK | Details ... | 102 | 0.302745 | 1.203828 | 0.102857 | 0.265056 | 0.533 | 3125 | tags=28%, list=19%, signal=35% |
| HALLMARK_ADIPOGENESIS | HALLMARK | Details ... | 199 | 0.254104 | 1.11473 | 0.117647 | 0.478184 | 0.789 | 1343 | tags=13%, list=8%, signal=14% |
| HALLMARK_APICAL_JUNCTION | HALLMARK | Details ... | 198 | 0.253884 | 1.104451 | 0.15 | 0.466385 | 0.822 | 2540 | tags=26%, list=16%, signal=30% |
| HALLMARK_NOTCH_SIGNALING | HALLMARK | Details ... | 32 | 0.3402 | 1.080863 | 0.33557 | 0.502087 | 0.869 | 2912 | tags=44%, list=18%, signal=53% |
| HALLMARK_TGF_BETA_SIGNALING | HALLMARK | Details ... | 54 | 0.296586 | 1.070154 | 0.292887 | 0.497865 | 0.888 | 3451 | tags=30%, list=21%, signal=37% |
| HALLMARK_ANGIOGENESIS | HALLMARK | Details ... | 36 | 0.323203 | 1.048438 | 0.379061 | 0.530283 | 0.925 | 745 | tags=19%, list=5%, signal=20% |
| HALLMARK_ANDROGEN_RESPONSE | HALLMARK | Details ... | 98 | 0.257397 | 1.00991 | 0.419753 | 0.637822 | 0.964 | 3090 | tags=34%, list=19%, signal=41% |
| HALLMARK_MITOTIC_SPINDLE | HALLMARK | Details ... | 199 | 0.213093 | 0.91685 | 0.804124 | 0.97006 | 0.997 | 3416 | tags=27%, list=21%, signal=34% |
| HALLMARK_FATTY_ACID_METABOLISM | HALLMARK | Details ... | 155 | 0.217838 | 0.902674 | 0.786207 | 0.959949 | 0.999 | 2576 | tags=22%, list=16%, signal=26% |
| HALLMARK_HEDGEHOG_SIGNALING | HALLMARK | Details ... | 36 | 0.267271 | 0.878707 | 0.699634 | 0.976193 | 1 | 1974 | tags=25%, list=12%, signal=28% |
| HALLMARK_PI3K_AKT_MTOR_SIGNALING | HALLMARK | Details ... | 105 | 0.204001 | 0.81685 | 0.889503 | 1 | 1 | 3376 | tags=28%, list=21%, signal=35% |
| HALLMARK_GLYCOLYSIS | HALLMARK | Details ... | 198 | 0.185428 | 0.813963 | 0.979381 | 0.993998 | 1 | 2575 | tags=18%, list=16%, signal=21% |
| HALLMARK_DNA_REPAIR | HALLMARK | Details ... | 149 | 0.132624 | 0.564831 | 1 | 1 | 1 | 1404 | tags=6%, list=9%, signal=7% |
| HALLMARK_OXIDATIVE_PHOSPHORYLATION | HALLMARK_OXIDATIVI | | 196 | 0.114629 | 0.499664 | 1 | 0.999842 | 1 | 2898 | tags=11%, list=18%, signal=13% |

**Supplementary Table 3. Hallmark gene sets enriched in pTKO based on GSEA analysis**

| NAME | GS<br> foll | GS DETAILS | SIZE | ES | NES | NOM p-val | FDR q-val | FWER p-va | RANK AT M | LEADING EDGE |
|---|---|---|---|---|---|---|---|---|---|---|
| HALLMARK | HALLMARK | Details ... | 87 | -0.541085 | -1.757537 | 0 | 0.004714 | 0.007 | 3042 | tags=36%, list=19%, signal=44% |
| HALLMARK | HALLMARK | Details ... | 138 | -0.485082 | -1.679708 | 0.002378 | 0.01075 | 0.032 | 1791 | tags=25%, list=11%, signal=28% |
| HALLMARK | HALLMARK | Details ... | 196 | -0.440753 | -1.565313 | 0 | 0.030251 | 0.127 | 3146 | tags=33%, list=19%, signal=40% |
| HALLMARK | HALLMARK | Details ... | 192 | -0.437684 | -1.555462 | 0.001114 | 0.025084 | 0.137 | 3530 | tags=35%, list=22%, signal=44% |
| HALLMARK | HALLMARK | Details ... | 197 | -0.431651 | -1.536424 | 0 | 0.025047 | 0.172 | 2831 | tags=34%, list=17%, signal=41% |
| HALLMARK | HALLMARK | Details ... | 195 | -0.420839 | -1.492809 | 0.003326 | 0.034789 | 0.268 | 2853 | tags=27%, list=18%, signal=32% |
| HALLMARK | HALLMARK | Details ... | 134 | -0.43052 | -1.46266 | 0.005917 | 0.042722 | 0.363 | 3398 | tags=31%, list=21%, signal=38% |
| HALLMARK | HALLMARK | Details ... | 94 | -0.446384 | -1.43757 | 0.022059 | 0.050104 | 0.456 | 4812 | tags=39%, list=30%, signal=56% |
| HALLMARK | HALLMARK | Details ... | 193 | -0.368706 | -1.314121 | 0.035714 | 0.163173 | 0.906 | 3146 | tags=31%, list=19%, signal=38% |
| HALLMARK | HALLMARK | Details ... | 195 | -0.369042 | -1.312462 | 0.03456 | 0.149034 | 0.908 | 3111 | tags=33%, list=19%, signal=41% |
| HALLMARK | HALLMARK | Details ... | 199 | -0.362296 | -1.272883 | 0.04918 | 0.198066 | 0.967 | 2411 | tags=24%, list=15%, signal=27% |
| HALLMARK | HALLMARK | Details ... | 196 | -0.337563 | -1.204735 | 0.108259 | 0.328919 | 0.997 | 2201 | tags=19%, list=14%, signal=22% |
| HALLMARK | HALLMARK | Details ... | 198 | -0.334753 | -1.184469 | 0.132675 | 0.355031 | 1 | 3278 | tags=31%, list=20%, signal=38% |
| HALLMARK | HALLMARK | Details ... | 44 | -0.390157 | -1.145252 | 0.269863 | 0.435107 | 1 | 3146 | tags=30%, list=19%, signal=37% |
| HALLMARK | HALLMARK | Details ... | 40 | -0.383709 | -1.10225 | 0.323651 | 0.536166 | 1 | 4029 | tags=38%, list=25%, signal=50% |
| HALLMARK | HALLMARK | Details ... | 160 | -0.311283 | -1.092749 | 0.287356 | 0.534745 | 1 | 2062 | tags=16%, list=13%, signal=18% |
| HALLMARK | HALLMARK | Details ... | 197 | -0.268418 | -0.956253 | 0.561521 | 0.964411 | 1 | 2168 | tags=18%, list=13%, signal=20% |
| HALLMARK | HALLMARK | Details ... | 41 | -0.297249 | -0.856756 | 0.718232 | 1 | 1 | 1911 | tags=15%, list=12%, signal=17% |
| HALLMARK | HALLMARK | Details ... | 197 | -0.236648 | -0.845567 | 0.815556 | 1 | 1 | 3086 | tags=17%, list=19%, signal=21% |
| HALLMARK | HALLMARK | Details ... | 156 | -0.242522 | -0.838767 | 0.820069 | 1 | 1 | 3383 | tags=21%, list=21%, signal=26% |
| HALLMARK | HALLMARK_E2F_TARG | | 199 | -0.227294 | -0.808022 | 0.886541 | 1 | 1 | 5471 | tags=35%, list=34%, signal=52% |
| HALLMARK | HALLMARK_UNFOLDEI | | 112 | -0.241049 | -0.807488 | 0.839853 | 1 | 1 | 5014 | tags=34%, list=31%, signal=49% |
| HALLMARK | HALLMARK_BILE_ACID | | 112 | -0.225903 | -0.756654 | 0.915194 | 1 | 1 | 3341 | tags=21%, list=21%, signal=27% |
| HALLMARK | HALLMARK_REACTIVE_ | | 49 | -0.234716 | -0.700648 | 0.916216 | 1 | 1 | 1919 | tags=12%, list=12%, signal=14% |
| HALLMARK | HALLMARK_MYC_TAR | | 57 | -0.217081 | -0.666315 | 0.961178 | 1 | 1 | 6605 | tags=46%, list=41%, signal=77% |
| HALLMARK | HALLMARK_HEME_ME | | 189 | -0.180501 | -0.63861 | 0.998915 | 1 | 1 | 3516 | tags=18%, list=22%, signal=23% |
| HALLMARK | HALLMARK_PROTEIN_ | | 94 | -0.168866 | -0.551678 | 0.998805 | 1 | 1 | 4264 | tags=18%, list=26%, signal=24% |
| HALLMARK | HALLMARK_G2M_CHE | | 195 | -0.1462 | -0.516155 | 1 | 1 | 1 | 4324 | tags=21%, list=27%, signal=28% |
| HALLMARK | HALLMARK_MYC_TAR | | 199 | -0.111296 | -0.398058 | 1 | 0.999954 | 1 | 8218 | tags=47%, list=51%, signal=94% |

**Supplementary Table 4. The transciptome upstream analysis with IPA between pDKO and pTKO**

| Upstream F | Expr Fold C | Molecule Type | Predicted A | Activation : | p-value of overlap |
|---|---|---|---|---|---|
| ANXA1 | -3.17 | enzyme | | | 0.00352 |
| DNM2 | -2.3 | enzyme | | | 0.031 |
| EFNA4 | -2.05 | kinase | | -1 | 0.000324 |
| FUT9 | -2.02 | enzyme | | | 0.0156 |
| Pvr | -1.94 | other | | | 0.00264 |
| PTEN | -1.84 | phosphatase | | 0.123 | 0.00000621 |
| MAPK8 | -1.71 | kinase | | 1 | 0.0435 |
| MMP14 | -1.66 | peptidase | | | 0.00218 |
| KIT | -1.66 | transmembrane rec | | | 0.0452 |
| S100A4 | -1.64 | other | | | 0.0357 |
| POR | -1.61 | enzyme | | -0.152 | 0.0031 |
| PDLIM2 | -1.59 | other | | 0 | 0.00644 |
| RBM5 | -1.57 | other | | | 0.00528 |
| VCAN | -1.57 | other | | -0.243 | 0.00162 |
| RBM39 | -1.55 | transcription regula | | | 0.00784 |
| MARK2 | -1.49 | kinase | | | 0.0291 |
| MYO1C | -1.48 | enzyme | | | 0.0156 |
| CAV1 | -1.47 | transmembrane rec | | | 0.0246 |
| S100A6 | -1.45 | transporter | | | 0.0216 |
| COL18A1 | -1.44 | other | | | 0.033 |
| JAG2 | -1.44 | growth factor | | | 0.0375 |
| SCARB1 | -1.42 | transporter | | | 0.0411 |
| TP53 | -1.42 | transcription regula | Inhibited | -2.255 | 0.000225 |
| STAT3 | -1.41 | transcription regula | | -0.152 | 0.0164 |
| LPCAT3 | -1.39 | enzyme | | | 0.031 |
| Muc1 | -1.39 | transmembrane rec | | | 0.0386 |
| ZKSCAN3 | -1.39 | transcription regula | | | 0.031 |
| MBD3 | -1.37 | other | | | 0.00397 |
| TGFB1 | -1.37 | growth factor | | -1.521 | 0.000000353 |
| PHIP | -1.36 | other | | | 0.00784 |
| SOCS6 | -1.36 | other | | | 0.0116 |
| FKBP1A | -1.36 | enzyme | | | 0.0386 |
| IL33 | -1.35 | cytokine | | | 0.042 |
| CTSD | -1.35 | peptidase | | | 0.031 |
| FN1 | -1.35 | enzyme | | 0.35 | 0.00607 |
| KITLG | -1.35 | growth factor | | | 0.0369 |
| CTR9 | -1.33 | other | | | 0.0139 |
| CD44 | -1.31 | other | | 0.577 | 0.0000374 |
| LMNA | -1.31 | other | | | 0.0442 |
| STAT1 | -1.3 | transcription regula | | 0.97 | 0.00642 |
| KLF6 | -1.3 | transcription regula | | | 0.0184 |
| TAP1 | -1.29 | transporter | | | 0.031 |
| EPAS1 | -1.28 | transcription regula | | | 0.0225 |
| JUNB | -1.28 | transcription regula | | | 0.017 |
| SCIN | -1.27 | other | | | 0.031 |
| HMGB2 | -1.27 | transcription regula | | | 0.0176 |
| FKBP8 | -1.26 | other | | | 0.0461 |
| AHNAK | -1.26 | other | | | 0.0233 |
| NSD2 | -1.25 | enzyme | | | 0.0357 |
| PRKCD | -1.25 | kinase | | -0.555 | 0.0172 |
| IKBKB | -1.24 | kinase | | 0.847 | 0.0000832 |
| CLDN7 | -1.24 | other | | 0 | 0.0161 |

| | | | | | |
|---|---|---|---|---|---|
| LDLR | -1.24 | transporter | | | 0.00799 |
| KLF3 | -1.24 | transcription regula | | 0.378 | 0.0168 |
| RAE1 | -1.24 | other | | | 0.00958 |
| EEF2K | -1.24 | kinase | | | 0.031 |
| JUN | -1.23 | transcription regula | | -0.563 | 0.00101 |
| CYTH1 | -1.23 | other | | | 0.0233 |
| TBC1D10A | -1.23 | other | | | 0.0233 |
| TNC | -1.23 | other | | | 0.0486 |
| ZDHHC7 | -1.22 | enzyme | | | 0.0156 |
| GLIS2 | -1.22 | transcription regula | | | 0.00861 |
| APP | -1.22 | other | Activated | 2.411 | 0.00445 |
| SFTPA1 | -1.22 | transporter | | | 0.0243 |
| G6PC3 | -1.21 | phosphatase | | | 0.0386 |
| ACAT2 | -1.21 | enzyme | | | 0.00784 |
| ERO1A | -1.21 | enzyme | | | 0.0233 |
| MTPN | -1.21 | transcription regula | | | 0.0267 |
| HMG20A | -1.2 | transcription regula | | | 0.0461 |
| STAT4 | -1.2 | transcription regula | | 1.798 | 0.00878 |
| STARD3 | -1.2 | transporter | | | 0.00784 |
| PHF2 | -1.19 | enzyme | | | 0.00784 |
| SOX2 | -1.19 | transcription regula | | -1.151 | 0.0156 |
| PURB | -1.19 | transcription regula | | | 0.0233 |
| GNA14 | -1.17 | enzyme | | | 0.00224 |
| INHBA | -1.17 | growth factor | | 0.385 | 0.0186 |
| INHA | -1.17 | growth factor | Inhibited | -2.201 | 0.00144 |
| BCL6 | -1.17 | transcription regula | | | 0.0261 |
| NRAS | -1.17 | enzyme | | | 0.00424 |
| CIC | -1.16 | transcription regula | | | 0.00321 |
| KCNE3 | -1.16 | ion channel | | | 0.00021 |
| ARNT2 | -1.16 | transcription regula | | -1.342 | 0.0238 |
| MADD | -1.15 | other | | | 0.0233 |
| ABL1 | -1.15 | kinase | | | 0.0393 |
| PCYT1A | -1.15 | enzyme | | | 0.00096 |
| GNAQ | -1.15 | enzyme | | | 0.014 |
| MTTP | -1.14 | transporter | | | 0.0291 |
| GAS2L3 | -1.13 | other | | | 0.00683 |
| NAB1 | -1.13 | transcription regula | | | 0.0233 |
| CHUK | -1.13 | kinase | | 0.679 | 0.000269 |
| TERT | -1.13 | enzyme | | | 0.00081 |
| CSF3 | -1.13 | cytokine | | 0.277 | 0.0127 |
| ERCC1 | -1.13 | enzyme | | | 0.0461 |
| CDH1 | -1.13 | other | | | 0.0259 |
| GTF2B | -1.13 | transcription regula | | | 0.0324 |
| WWP1 | -1.13 | enzyme | | | 0.0233 |
| HRAS | -1.13 | enzyme | | -1.633 | 0.000619 |
| TFAP2A | -1.12 | transcription regula | | 0.931 | 0.00132 |
| CSF1 | -1.12 | cytokine | | 1.413 | 0.0193 |
| MACROH2/ | -1.12 | other | | | 0.00969 |
| PRKACA | -1.12 | kinase | | | 0.0486 |
| EFNA2 | -1.12 | kinase | | -1 | 0.00104 |
| MXD1 | -1.11 | transcription regula | | | 0.00321 |
| SCGB1A1 | -1.11 | cytokine | | | 0.0116 |
| CREM | -1.1 | transcription regula | | -1.432 | 0.00137 |
| PELI2 | -1.1 | enzyme | | | 0.031 |
| ZAP70 | -1.1 | kinase | | | 0.0102 |

| | | | | |
|---|---|---|---|---|
| TCF7L2 | -1.1 | transcription regula | -1.664 | 0.0473 |
| CTNNB1 | -1.1 | transcription regula | -0.666 | 0.0000102 |
| IGF1 | -1.09 | growth factor | -0.614 | 0.000107 |
| ACSL1 | -1.09 | enzyme | | 0.00784 |
| IFNAR1 | -1.09 | transmembrane rec | | 0.0287 |
| SIM1 | -1.09 | transcription regula | -1.342 | 0.0265 |
| ALOX15 | -1.09 | enzyme | | 0.0291 |
| NR3C2 | -1.09 | ligand-dependent n | | 0.0129 |
| PSMC5 | -1.08 | transcription regula | | 0.0233 |
| XDH | -1.07 | enzyme | | 0.0324 |
| CFB | -1.07 | peptidase | | 0.0077 |
| PRKCZ | -1.07 | kinase | | 0.026 |
| PKD1 | -1.07 | ion channel | -0.447 | 0.0161 |
| LPAR1 | -1.07 | G-protein coupled r | | 0.00451 |
| ERBB2 | -1.07 | kinase | 1.195 | 0.0161 |
| GSTP1 | -1.06 | enzyme | | 0.0113 |
| TJP1 | -1.06 | other | | 0.0156 |
| BMP4 | -1.06 | growth factor | -0.447 | 0.0225 |
| DUSP1 | -1.06 | phosphatase | | 0.0452 |
| KCTD11 | -1.06 | other | | 0.0386 |
| APOE | -1.06 | transporter | | 0.0429 |
| LHCGR | -1.05 | G-protein coupled r | | 0.00327 |
| Nppb | -1.05 | other | | 0.031 |
| NR1H2 | -1.05 | ligand-dependent n | | 0.0062 |
| TP73 | -1.05 | transcription regula | -1.528 | 0.0133 |
| RFXANK | -1.05 | transcription regula | | 0.0386 |
| TTC39B | -1.04 | other | | 0.0156 |
| CHRNA7 | -1.04 | transmembrane rec | | 0.0291 |
| TOR1A | -1.04 | enzyme | | 0.0077 |
| AR | -1.04 | ligand-dependent n | -1.452 | 0.0157 |
| ZYX | -1.04 | other | | 0.0461 |
| SUMO2 | -1.03 | enzyme | | 0.00435 |
| PSMC4 | -1.03 | peptidase | | 0.00784 |
| ARHGEF11 | -1.03 | other | | 0.0233 |
| THRB | -1.02 | ligand-dependent n | | 0.0284 |
| IL22 | -1.02 | cytokine | 0.927 | 0.0133 |
| MYD88 | -1.02 | other | 0.009 | 0.0000481 |
| SYVN1 | -1.02 | transporter | 0 | 0.0224 |
| CRH | -1.02 | cytokine | | 0.0151 |
| TAL1 | -1.02 | transcription regula | | 0.00878 |
| CLEC4M | -1.02 | other | | 0.0461 |
| AGT | -1.01 | growth factor | -0.127 | 0.00025 |
| ESR2 | -1.01 | ligand-dependent n | 0.371 | 0.0207 |
| TLR2 | -1.01 | transmembrane rec | 1.013 | 0.0459 |
| SUMO3 | -1.01 | other | | 0.0357 |
| TRPC1 | -1.01 | ion channel | | 0.0127 |
| CDR2 | -1.01 | other | | 0.0386 |
| EZH2 | -1.01 | transcription regula | 1.671 | 0.00708 |
| F7 | -1.01 | peptidase | | 0.026 |
| NR1I2 | -1.01 | ligand-dependent n | | 0.0397 |
| GPS1 | -1.01 | other | | 0.0233 |
| MPO | -1.01 | enzyme | | 0.0233 |
| FSHB | -1.01 | other | | 0.0116 |
| RETNLB | -1 | other | | 0.0227 |
| SOAT2 | -1 | enzyme | | 0.0386 |

| | | | | | |
|---|---|---|---|---|---|
| TWIST1 | -1 | transcription regula | | -0.113 | 0.000236 |
| NKX2-1 | -1 | transcription regula | | | 0.0148 |
| WIF1 | -1 | other | | | 0.0461 |
| CYP8B1 | -1 | enzyme | | | 0.031 |
| IFIH1 | -1 | enzyme | | | 0.0375 |
| FGF19 | -1 | growth factor | | | 0.0293 |
| RC3H1 | -1 | enzyme | | | 0.0202 |
| WNT3A | -1 | cytokine | | -1.673 | 0.00443 |
| BUD23 | 1 | enzyme | | | 0.0386 |
| WNT1 | 1 | cytokine | | -0.749 | 0.00364 |
| FGF2 | 1 | growth factor | | 1.428 | 0.0000191 |
| IL13 | 1 | cytokine | | -1.786 | 0.000261 |
| CRABP2 | 1 | transporter | | | 0.031 |
| NPC1 | 1.01 | transporter | | | 0.00465 |
| LEO1 | 1.01 | other | | | 0.031 |
| LRP6 | 1.01 | transmembrane rec | | | 0.0357 |
| ALDH1A2 | 1.01 | enzyme | | | 0.00199 |
| TSC2 | 1.01 | other | | | 0.0166 |
| CANX | 1.01 | other | | | 0.031 |
| TFF2 | 1.01 | other | | | 0.0461 |
| DSPP | 1.01 | other | | | 0.0461 |
| BMP6 | 1.01 | growth factor | | | 0.00719 |
| WT1 | 1.01 | transcription regula | | -0.564 | 0.000566 |
| GNA15 | 1.01 | enzyme | | | 0.0121 |
| KDM4C | 1.02 | enzyme | | | 0.026 |
| AMH | 1.02 | growth factor | | | 0.0189 |
| LEP | 1.02 | growth factor | | -1.901 | 0.47 |
| LUM | 1.02 | other | | | 0.00264 |
| MED24 | 1.02 | transcription regula | | | 0.0233 |
| KRT18 | 1.02 | other | | | 0.00784 |
| FOXO3 | 1.02 | transcription regula | | | 0.014 |
| TNFRSF1A | 1.02 | transmembrane rec | | | 0.0367 |
| OSM | 1.02 | cytokine | | 1.242 | 0.0000885 |
| PDX1 | 1.02 | transcription regula | Activated | 2 | 0.00373 |
| FOXP3 | 1.03 | transcription regula | | -1.067 | 0.000181 |
| NEUROG1 | 1.03 | transcription regula | | | 0.0145 |
| ANGPT1 | 1.03 | growth factor | | | 0.0393 |
| HNF1A | 1.03 | transcription regula | | 0.555 | 0.0245 |
| FOXH1 | 1.03 | transcription regula | | | 0.031 |
| ARHGAP21 | 1.03 | other | | | 0.0448 |
| TXN2 | 1.03 | enzyme | | | 0.031 |
| Prl7d1 | 1.04 | cytokine | | | 0.00784 |
| USP22 | 1.04 | peptidase | | | 0.031 |
| SMAD1 | 1.04 | transcription regula | | | 0.0411 |
| FGF10 | 1.04 | growth factor | | | 0.00922 |
| DAB1 | 1.04 | other | | | 0.0386 |
| CCL22 | 1.04 | cytokine | | | 0.0386 |
| NLRP12 | 1.04 | other | | | 0.0028 |
| COMMD1 | 1.04 | transporter | | | 0.0163 |
| CFD | 1.04 | peptidase | | | 0.00784 |
| CYP1A1 | 1.04 | enzyme | | | 0.0441 |
| SNAI1 | 1.04 | transcription regula | | -1.131 | 0.0000481 |
| PSMD6 | 1.04 | enzyme | | | 0.00784 |
| IFI30 | 1.05 | enzyme | | | 0.0156 |
| PGR | 1.05 | ligand-dependent n | | -1.484 | 0.00166 |

| | | | | |
|---|---|---|---|---|
| KDM1A | 1.05 | enzyme | | 0.043 |
| HNRNPA2B | 1.05 | other | | 0.0287 |
| LSS | 1.05 | enzyme | | 0.0233 |
| PRDM1 | 1.05 | transcription regula | -1.387 | 0.0358 |
| IFNG | 1.05 | cytokine | 0.564 | 0.0000217 |
| ZMPSTE24 | 1.06 | peptidase | | 0.0486 |
| INPP5D | 1.06 | phosphatase | | 0.0357 |
| STX12 | 1.06 | other | | 0.00784 |
| CEBPA | 1.06 | transcription regula | -0.726 | 0.00295 |
| TNF | 1.06 | cytokine | 0.303 | 9.27E-10 |
| FOXM1 | 1.06 | transcription regula | -0.072 | 0.000642 |
| KDR | 1.06 | kinase | | 0.0467 |
| SERPIND1 | 1.06 | other | | 0.0461 |
| RHOJ | 1.07 | enzyme | | 0.00958 |
| ZNF24 | 1.07 | transcription regula | | 0.031 |
| PSMA2 | 1.07 | peptidase | | 0.00784 |
| PDCD6 | 1.07 | other | | 0.00784 |
| KMT5B | 1.08 | enzyme | | 0.00125 |
| OSCAR | 1.08 | other | | 0.0291 |
| SRF | 1.08 | transcription regula | -1.98 | 0.283 |
| AP1M1 | 1.08 | transporter | | 0.0156 |
| PAPPA | 1.08 | peptidase | | 0.0386 |
| ELAVL1 | 1.08 | other | | 0.0497 |
| H60a | 1.08 | other | | 0.0233 |
| INSIG1 | 1.08 | other | | 0.0463 |
| MEOX2 | 1.08 | transcription regula | | 0.00352 |
| SFN | 1.08 | other | | 0.00134 |
| Hbb-b2 | 1.08 | other | | 0.0393 |
| ASGR2 | 1.08 | transmembrane rec | | 0.0156 |
| ADAMTS5 | 1.08 | peptidase | | 0.0156 |
| SMO | 1.08 | G-protein coupled r | | 0.0259 |
| DYSF | 1.08 | other | | 0.00347 |
| NEDD9 | 1.08 | other | | 0.0375 |
| MED4 | 1.08 | transcription regula | | 0.00784 |
| CUL4A | 1.09 | other | | 0.0233 |
| SPRY2 | 1.09 | other | | 0.0357 |
| TAF10 | 1.09 | transcription regula | | 0.0461 |
| KLF4 | 1.09 | transcription regula | -1.364 | 0.000214 |
| S100A8 | 1.09 | other | -0.896 | 0.000523 |
| FGF7 | 1.09 | growth factor | | 0.019 |
| RARRES2 | 1.09 | transmembrane rec | | 0.00958 |
| IL6 | 1.09 | cytokine | 0.159 | 0.000132 |
| FLT3 | 1.09 | kinase | | 0.0375 |
| ERCC3 | 1.09 | enzyme | | 0.0461 |
| NGF | 1.1 | growth factor | | 0.00896 |
| DMD | 1.1 | other | | 0.0279 |
| CFI | 1.1 | peptidase | | 0.00784 |
| GTF2H4 | 1.1 | transcription regula | | 0.0386 |
| LRP1B | 1.1 | transmembrane rec | | 0.0156 |
| DNM1 | 1.1 | enzyme | | 0.0386 |
| CDK2AP1 | 1.1 | other | | 0.00465 |
| ACAT1 | 1.11 | enzyme | | 0.0233 |
| ACOX1 | 1.11 | enzyme | 1 | 0.0269 |
| PHLPP1 | 1.11 | enzyme | | 0.0077 |
| SNTB1 | 1.11 | other | | 0.00784 |

| | | | | | |
|---|---|---|---|---|---|
| KRAS | 1.11 | enzyme | | 1.105 | 0.000000015 |
| ANLN | 1.11 | other | | | 0.00749 |
| ELANE | 1.11 | peptidase | | | 0.0307 |
| FOXL1 | 1.11 | transcription regula | | | 0.0156 |
| SPRY4 | 1.11 | other | | | 0.0386 |
| UTP3 | 1.12 | other | | | 0.031 |
| ZMYND10 | 1.12 | other | | | 0.0461 |
| RBM14 | 1.12 | transcription regula | | | 0.0386 |
| Raet1d/Rae | 1.12 | other | | | 0.00784 |
| CYP4A11 | 1.12 | enzyme | | | 0.000898 |
| PAX5 | 1.12 | transcription regula | | | 0.0467 |
| CARD14 | 1.12 | other | | | 0.0461 |
| SMAD3 | 1.12 | transcription regula | | 1.406 | 0.0381 |
| HOXD3 | 1.12 | transcription regula | | | 0.0106 |
| SHOX2 | 1.12 | transcription regula | | | 0.0461 |
| CYTH3 | 1.12 | other | | | 0.0156 |
| TFAP2C | 1.12 | transcription regula | | 1.919 | 0.00288 |
| TLR3 | 1.12 | transmembrane rec | | 1.948 | 0.0537 |
| CDK19 | 1.13 | kinase | | | 0.031 |
| TENM1 | 1.13 | transmembrane rec | | | 0.00212 |
| CREB1 | 1.13 | transcription regula | | -0.231 | 0.000552 |
| DNM1L | 1.13 | enzyme | | | 0.0461 |
| EFNA3 | 1.13 | kinase | | -1 | 0.000355 |
| ALB | 1.14 | transporter | | | 0.0467 |
| CELA1 | 1.14 | peptidase | | | 0.0156 |
| ADIPOQ | 1.14 | other | | 0 | 0.0443 |
| TIMP4 | 1.14 | other | | | 0.0386 |
| CSF2 | 1.14 | cytokine | | -0.483 | 0.00000356 |
| MITF | 1.14 | transcription regula | | -0.557 | 0.00272 |
| IL1A | 1.14 | cytokine | | -0.275 | 0.00914 |
| IKBKG | 1.14 | kinase | | 0.747 | 0.000101 |
| PBX3 | 1.14 | transcription regula | | | 0.0127 |
| OLR1 | 1.14 | transmembrane rec | | | 0.00597 |
| EIF4G2 | 1.15 | translation regulatc | | | 0.0106 |
| MAPT | 1.15 | other | | | 0.0193 |
| PHF19 | 1.15 | other | | | 0.031 |
| AGTR1 | 1.15 | G-protein coupled r | | | 0.0393 |
| RUVBL1 | 1.15 | transcription regula | | | 0.0393 |
| DDX1 | 1.15 | enzyme | | | 0.00784 |
| FOS | 1.15 | transcription regula | | -0.896 | 0.0289 |
| TSPYL5 | 1.16 | other | | | 0.0139 |
| TRPM7 | 1.16 | kinase | | | 0.0461 |
| DNAJC3 | 1.16 | other | | | 0.00523 |
| TANK | 1.16 | other | | | 0.034 |
| LRP8 | 1.16 | transmembrane rec | | | 0.0386 |
| PRDX4 | 1.16 | enzyme | | | 0.0233 |
| CGA | 1.16 | other | | | 0.00181 |
| ZEB1 | 1.17 | transcription regula | | | 0.00749 |
| PRL | 1.18 | cytokine | Inhibited | -2.622 | 0.00647 |
| PPARG | 1.18 | ligand-dependent n | | -0.088 | 0.000121 |
| CEBPB | 1.18 | transcription regula | | -0.309 | 0.00176 |
| FOXO4 | 1.18 | transcription regula | | | 0.00644 |
| ECSCR | 1.18 | other | | | 0.00784 |
| PREB | 1.18 | transcription regula | | | 0.0156 |
| CR1L | 1.19 | transmembrane rec | | | 0.026 |

| | | | | | |
|---|---|---|---|---|---|
| LMO1 | 1.19 | transcription regula | | | 0.00125 |
| ID2 | 1.19 | transcription regula | | | 0.0024 |
| EFNA5 | 1.2 | kinase | | -1 | 0.000423 |
| F2RL1 | 1.2 | G-protein coupled r | | | 0.0139 |
| HNF4A | 1.2 | transcription regula | | 1.342 | 0.000454 |
| FOLR1 | 1.2 | transporter | | | 0.0378 |
| CCL5 | 1.2 | cytokine | | -0.094 | 0.00035 |
| MMP10 | 1.22 | peptidase | | | 0.0156 |
| GRIN2B | 1.22 | ion channel | | | 0.0386 |
| ERAP1 | 1.23 | peptidase | | | 0.0461 |
| TLR4 | 1.23 | transmembrane rec | | 1.462 | 0.0106 |
| CCN2 | 1.24 | growth factor | Inhibited | -2.236 | 0.00144 |
| NFKBIA | 1.24 | transcription regula | | -0.295 | 0.0000106 |
| CCN1 | 1.24 | other | | | 0.0302 |
| NET1 | 1.25 | other | | | 0.0461 |
| ARHGDIG | 1.26 | other | | | 0.000355 |
| MAP3K8 | 1.26 | kinase | | | 0.0319 |
| KCNN4 | 1.27 | ion channel | | | 0.00264 |
| NCOA1 | 1.27 | transcription regula | | | 0.00104 |
| IL10RA | 1.28 | transmembrane rec | | -0.788 | 0.0033 |
| JMY | 1.28 | transcription regula | | | 0.0233 |
| CDKN2A | 1.3 | transcription regula | | 0.447 | 0.0296 |
| FBXO32 | 1.3 | enzyme | | 0.728 | 0.0000522 |
| KLF2 | 1.31 | transcription regula | | | 0.00147 |
| EGR1 | 1.31 | transcription regula | | 1.342 | 0.000118 |
| NR3C1 | 1.32 | ligand-dependent n | Inhibited | -2.163 | 0.00694 |
| RGS1 | 1.32 | enzyme | | | 0.006 |
| RET | 1.33 | kinase | | | 0.00206 |
| S100A9 | 1.35 | other | | -0.277 | 0.000383 |
| IL1B | 1.35 | cytokine | | -0.483 | 0.00026 |
| TNFRSF1B | 1.36 | transmembrane rec | | | 0.00827 |
| SPRY1 | 1.36 | other | | | 0.0176 |
| NFE2L2 | 1.37 | transcription regula | | -1.407 | 0.0178 |
| SATB1 | 1.42 | transcription regula | | 0.895 | 0.0000557 |
| ADM | 1.42 | other | | | 0.00749 |
| VLDLR | 1.45 | transporter | | | 0.0386 |
| NCF4 | 1.49 | enzyme | | | 0.00264 |
| MYC | 1.5 | transcription regula | | -1.597 | 0.000057 |
| RASSF6 | 1.53 | other | | | 0.0386 |
| SAMSN1 | 1.6 | other | | 1 | 0.00575 |
| ESR1 | 1.6 | ligand-dependent n | | 0.261 | 0.000041 |
| EFNA1 | 1.62 | other | | | 0.0145 |
| MDK | 1.63 | growth factor | | | 0.00683 |
| DPP4 | 1.65 | peptidase | | | 0.0189 |
| CFH | 1.7 | other | | | 0.031 |
| EGF | 1.72 | growth factor | | -0.071 | 0.0204 |
| NKX3-1 | 1.78 | transcription regula | | | 0.0000592 |
| ETV5 | 1.83 | transcription regula | | 0.447 | 4.26E-09 |
| TP63 | 1.98 | transcription regula | | 0.02 | 0.00351 |
| LPAR3 | 1.98 | G-protein coupled r | | | 0.031 |
| NECTIN3 | 2.06 | other | | | 0.00784 |
| SAA1 | 4.74 | transporter | | | 0.0163 |
| Ap2 | | group | | | 0.0324 |
| Laminin (fa | | group | | | 0.0233 |
| lysophosph | | chemical - other | | 0.86 | 0.00668 |

| | | | |
|---|---|---|---|
| 15-deoxy-d | chemical - endogen | -1.076 | 0.0148 |
| 3-deazanep | chemical drug | | 0.0128 |
| inosine | chemical - endogen | | 0.00749 |
| zidovudine | chemical drug | | 0.0139 |
| 8-hydroxyg | chemical - endogen | | 0.0461 |
| 6-dimethyl | chemical - endogen | | 0.031 |
| allopurinol | chemical drug | | 0.0284 |
| PD 180970 | chemical drug | | 0.031 |
| propylthiou | chemical drug | | 0.0104 |
| PD173074 | chemical reagent | -1.406 | 0.00035 |
| PP1 | chemical drug | | 0.0411 |
| darusentar | chemical drug | | 0.0386 |
| L-leucine | chemical - endogen | | 0.0156 |
| E. coli B5 li | chemical - endogen | 0.765 | 0.0169 |
| lipid A | chemical toxicant | | 0.0486 |
| E. coli B4 li | chemical toxicant | 0.943 | 0.00724 |
| pentosan p | chemical drug | | 0.0233 |
| beta-glucar | chemical drug | | 0.0386 |
| dextran sul | chemical drug | | 0.00366 |
| stigmaster | chemical - endogen | | 0.0233 |
| 17-alpha-e | chemical drug | -1 | 0.0339 |
| beta-estrac | chemical - endogen | -1.363 | 0.00000692 |
| chenodeox | chemical - endogen | | 0.0284 |
| hyodeoxycl | chemical - endogen | | 0.0233 |
| corticoster | chemical drug | | 0.0411 |
| hydrogen p | chemical - endogen | -0.355 | 0.0426 |
| GPIIB-IIIA | complex | | 0.0461 |
| bucladesin | chemical toxicant | -0.099 | 0.000469 |
| glucocortic | chemical drug | -1.807 | 0.00914 |
| mineralocc | chemical drug | | 0.0233 |
| corticoster | chemical - endogen | -1.076 | 0.00267 |
| Collagen ty | complex | | 0.0467 |
| 1-O-hexade | chemical - endogen | | 0.031 |
| dexametha | chemical drug | -0.65 | 0.000000425 |
| fluticasone | chemical drug | 0.254 | 0.000441 |
| deoxycortic | chemical drug | | 0.0151 |
| CARD9 | other | | 0.023 |
| triamcinolc | chemical drug | -0.647 | 0.000269 |
| cardiotoxin | chemical - other | | 0.0313 |
| trichloroetl | chemical toxicant | | 0.0107 |
| Hdac | group | -1.406 | 0.0103 |
| Creb | group | | 0.0382 |
| Rsk | group | | 0.031 |
| PI3K (comp | complex | -0.686 | 0.00867 |
| Tgf beta | group | 0.849 | 0.0137 |
| ERK1/2 | group | -0.527 | 0.00299 |
| DUB | group | | 0.0233 |
| P38 MAPK | group | -0.039 | 0.000174 |
| ATPase | group | | 0.0386 |
| RNA polym | complex | | 0.00599 |
| Camk | complex | | 0.0461 |
| MAP2K1/2 | group | 0.991 | 0.00363 |
| Calmodulir | group | -0.152 | 0.00119 |
| SERCA | group | | 0.0461 |
| Mek | group | 1.15 | 0.00933 |

| | | | |
|---|---|---|---|
| Fgf | group | | 0.0275 |
| Vegf | group | 0.97 | 0.047 |
| IL-1R | group | | 0.0176 |
| N-cor | group | | 0.0452 |
| Camkk | group | | 0.0156 |
| ZFAS1 | other | | 0.0233 |
| PDGF BB | complex | -0.324 | 0.00446 |
| Nr1h | group | 1.161 | 0.0326 |
| AMPK | complex | 0.603 | 0.000406 |
| CYP4A | group | | 0.0156 |
| NFkB (com | complex | 0.612 | 0.00449 |
| LDL | complex | 0.689 | 0.00382 |
| rutaecarpir | chemical - endogen | | 0.0156 |
| CRNDE | other | | 0.0324 |
| GC-GCR dir | complex | | 0.000246 |
| Dexametha | complex | | 0.0233 |
| Interferon a | group | 0.666 | 0.0082 |
| ERK | group | -0.651 | 0.00306 |
| Raf | group | | 0.0164 |
| olmesartar | chemical drug | | 0.0386 |
| FSH | complex | -0.41 | 0.00137 |
| RNU7-1 | other | | 0.0386 |
| E. coli sero | chemical - endogen | 1.387 | 0.0158 |
| colestimide | chemical drug | | 0.00784 |
| SNORD21 | other | | 0.00784 |
| cis-urocani | chemical drug | | 0.0127 |
| GPR119 | G-protein coupled r | | 0.0461 |
| gentamicin | chemical drug | -0.816 | 0.013 |
| Integrin alp | complex | | 0.0233 |
| ZNF366 | transcription regula | | 0.0156 |
| Tcp4 | other | | 0.00784 |
| Raet1a | other | | 0.0156 |
| Alpha cater | group | -0.739 | 0.00258 |
| Aldosteron | complex | | 0.0233 |
| Trk Recepto | group | | 0.0233 |
| IL-2R | complex | | 0.00166 |
| IgD | complex | | 0.0386 |
| Growth hor | group | 0.106 | 0.00972 |
| Ap1 gamma | group | | 0.0233 |
| black raspb | chemical drug | | 0.0121 |
| estrogen re | group | -0.35 | 0.000117 |
| AURK | group | | 0.00749 |
| CG | complex | -1.125 | 0.00014 |
| PTPase | group | | 0.0156 |
| beta-ionon | chemical - endogen | | 0.0156 |
| Cathepsin | group | | 0.0386 |
| Nc2 | complex | | 0.031 |
| CSF | group | | 0.0386 |
| Raet1b | other | | 0.0233 |
| CGB3 (inclu | other | | 0.00683 |
| entolimod | biologic drug | | 0.0461 |
| CR1 | transmembrane rec | | 0.0461 |
| IL3 | cytokine | | 0.0183 |
| TICAM2 | other | | 0.0189 |
| miR-291a-3 | mature microRNA | -0.017 | 0.00511 |

| | | | | |
|---|---|---|---|---|
| mir-223 | microRNA | | 0.152 | 0.0208 |
| mir-148 | microRNA | | | 0.000419 |
| miR-124-3p | mature microRNA | Activated | 2.236 | 0.0347 |
| miR-483-5p | mature microRNA | | | 0.0156 |
| miR-133a-3 | mature microRNA | | | 0.00749 |
| mir-133 | microRNA | | 1 | 0.000355 |
| miR-125b-5 | mature microRNA | | 1.067 | 0.00644 |
| miR-182-5p | mature microRNA | | | 0.0324 |
| miR-33-5p | mature microRNA | | | 0.00784 |
| mir-32 | microRNA | | | 0.031 |
| miR-709 (a | mature microRNA | | | 0.00784 |
| lynestrenol | chemical drug | | | 0.0156 |
| mir-150 | microRNA | | | 0.0275 |
| pituitary ac | biologic drug | | | 0.000242 |
| NRG1 | growth factor | | -0.025 | 0.00914 |
| ORM1 | other | | | 0.031 |
| FALEC | other | | | 0.00125 |
| GDF7 | growth factor | | | 0.0386 |
| 2-[[4-[(e)-s | chemical reagent | | | 0.031 |
| IFNA2 | cytokine | | -0.747 | 0.0381 |
| ITGA11 | other | | | 0.0461 |
| EPO | cytokine | | | 0.0271 |
| EPZ004777 | chemical reagent | | | 0.0233 |
| MEIS1 | transcription regula | | | 0.0291 |
| Hmgb2 (inc | transcription regula | | | 0.0461 |
| IL37 | cytokine | | | 0.0116 |
| ursodeoxyc | chemical reagent | | | 0.00784 |
| E-c-HDMAP | chemical reagent | | | 0.0156 |
| P2pal-18S | chemical reagent | | | 0.0233 |
| COG112 pe | chemical reagent | | | 0.0461 |
| C20-D3-vita | chemical drug | | | 0.0233 |
| J11-Cl | chemical reagent | | | 0.00321 |
| 101.10 pep | chemical reagent | | | 0.00683 |
| KRT7-AS | other | | | 0.00784 |
| cigarette sr | chemical toxicant | | 1.104 | 0.0468 |
| bleomycin | biologic drug | | -0.277 | 0.00318 |
| gentamicin | chemical drug | | | 0.0133 |
| PD 169316 | chemical drug | | | 0.0116 |
| halofuginor | chemical drug | | | 0.00075 |
| temocapril | chemical reagent | | | 0.0233 |
| phenylacet | chemical - endogen | | | 0.0233 |
| neopterin | chemical - endogen | | | 0.031 |
| rimonaban | chemical drug | | | 0.0176 |
| palytoxin | chemical toxicant | | | 0.0156 |
| LY294002 | chemical drug | | -0.042 | 0.00801 |
| gefitinib | chemical drug | | | 0.000397 |
| PD98059 | chemical - kinase in | | -0.655 | 0.000000104 |
| olmesartar | chemical drug | | | 0.0461 |
| carbamylch | chemical drug | | | 0.0393 |
| methylmer | chemical toxicant | | | 0.0151 |
| marimastat | chemical drug | | | 0.031 |
| tacrolimus | chemical drug | | | 0.0461 |
| Congo Red | chemical toxicant | | | 0.0461 |
| 1-butanol | chemical - endogen | | | 0.031 |
| JAK inhibito | chemical drug | | -1 | 0.0000557 |

| | | | | |
|---|---|---|---|---|
| quiflapon | chemical reagent | | | 0.0156 |
| tempol | chemical drug | | | 0.034 |
| dalfamprid | chemical drug | | | 0.0411 |
| rosiglitazor | chemical drug | | -1.526 | 0.0069 |
| 8-methyl-p | chemical reagent | | | 0.0156 |
| 4-phenylbu | chemical - endogen | | | 0.0276 |
| bortezomib | chemical drug | | -0.577 | 0.0375 |
| BB-2116 | chemical - protease | | | 0.0233 |
| famotidine | chemical drug | | | 0.0386 |
| emricasan | chemical drug | | | 0.0461 |
| phenylbuta | chemical drug | | | 0.0117 |
| indometha | chemical drug | | -0.57 | 0.00306 |
| nafoxidine | chemical drug | | | 0.00784 |
| okadaic aci | chemical toxicant | | 0.64 | 0.0000769 |
| nitrofurant | chemical drug | | -1.633 | 0.00201 |
| 4-methylni | chemical toxicant | | | 0.034 |
| pioglitazon | chemical drug | | -0.728 | 0.0427 |
| everolimus | chemical drug | | | 0.00199 |
| n-nitrosom | chemical toxicant | | | 0.0229 |
| ranitidine | chemical drug | | | 0.0386 |
| ethanol | chemical - endogen | | -1.198 | 0.000917 |
| fenamic ac | chemical reagent | | | 0.017 |
| H89 | chemical drug | | -1.192 | 0.0252 |
| U0126 | chemical drug | Inhibited | -2.064 | 0.000757 |
| troglitazon | chemical drug | Inhibited | -2.581 | 0.0299 |
| captopril | chemical drug | | -1.205 | 0.00302 |
| thioacetam | chemical toxicant | | -1.951 | 0.0000265 |
| paricalcitol | chemical drug | | | 0.0375 |
| LFM-A13 | chemical drug | | | 0.0156 |
| sphingosin | chemical - endogen | | 0.471 | 0.00575 |
| simvastatir | chemical drug | | -1.478 | 0.000286 |
| mevastatin | chemical drug | | | 0.00683 |
| calcitriol | chemical drug | | -0.788 | 0.0355 |
| triamteren | chemical drug | | | 0.0151 |
| oenothein | chemical - endogen | | | 0.0233 |
| 3,4-dihydro | chemical - endogen | | | 0.0233 |
| bexarotene | chemical drug | | -0.943 | 0.0182 |
| RP 73401 | chemical toxicant | | | 0.00212 |
| roflumilast | chemical drug | | | 0.0386 |
| (S)-2-((6Z,1 | chemical reagent | | | 0.0156 |
| SU5402 | chemical drug | | | 0.00451 |
| paxilline | chemical - endogen | | | 0.0386 |
| quinapril | chemical drug | | | 0.0461 |
| Ki16425 | chemical reagent | | | 0.0386 |
| nifedipine | chemical drug | | | 0.0212 |
| butyric acid | chemical - endogen | | -0.275 | 0.00903 |
| taprostene | chemical reagent | | | 0.0233 |
| trans-hydro | chemical drug | | -1 | 0.0136 |
| silver nitrat | chemical drug | | | 0.0233 |
| sodium chl | chemical - endogen | | | 0.0151 |
| cisplatin | chemical drug | Inhibited | -2.741 | 0.00568 |
| zymosan A | chemical - endogen | | | 0.0106 |
| isotretinoir | biologic drug | | -1 | 0.00288 |
| retinol acet | chemical drug | | | 0.00264 |
| arotinoid a | chemical toxicant | | | 0.00108 |

| | | | | |
|---|---|---|---|---|
| tretinoin | chemical - endogen | | -0.27 | 0.000000506 |
| alitretinoin | chemical drug | | 0.463 | 0.0346 |
| benzene | chemical toxicant | | | 0.0107 |
| Pam3-Cys-S | chemical reagent | | 0.391 | 0.0175 |
| vancomycin | biologic drug | | -0.246 | 0.000615 |
| actinomyci | chemical - endogen | | | 0.0233 |
| phosphorai | chemical - endogen | | | 0.0233 |
| actinomyci | biologic drug | | -1.997 | 0.119 |
| desmopres | biologic drug | | | 0.0368 |
| Z-VEID-FMI | chemical reagent | | | 0.0461 |
| N-Ac-Leu-L | chemical - protease | | | 0.0251 |
| romidepsin | biologic drug | | | 0.0164 |
| androgen | chemical drug | | | 0.000058 |
| doxorubicin | chemical drug | Inhibited | -2.2 | 0.223 |
| forskolin | chemical toxicant | | -0.33 | 0.000491 |
| betulinic ac | chemical drug | | | 0.023 |
| daidzein | chemical drug | | | 0.0368 |
| paclitaxel | chemical drug | | | 0.0093 |
| camptothe | chemical drug | | 0.632 | 0.00318 |
| D-lysergic a | chemical drug | | | 0.0156 |
| ciprofibrate | chemical drug | | -0.651 | 0.00594 |
| clofibrate | chemical drug | | -0.744 | 0.0011 |
| succinylace | chemical - endogen | | | 0.031 |
| palmitoleic | chemical - endogen | | | 0.00451 |
| poly rI:rC-R | biologic drug | | 1.758 | 0.0762 |
| mizoribine | chemical drug | | | 0.0233 |
| decitabine | chemical drug | | -1.856 | 0.0549 |
| cytarabine | chemical drug | | | 0.0151 |
| N-nitro-L-a | chemical drug | | 1.117 | 0.00597 |
| N-acetyl-L- | chemical drug | | -0.447 | 0.0314 |
| fosinopril | chemical drug | | | 0.0233 |
| creatine | chemical - endogen | | | 0.00958 |
| lipopolysac | chemical drug | | 0.117 | 0.0000147 |
| D-glucose | chemical - endogen | Inhibited | -2.092 | 0.0128 |
| cytokine | group | | | 0.0326 |
| allosamidin | chemical reagent | | | 0.0386 |
| tetradecan | chemical drug | | -0.414 | 2.71E-08 |
| cyproteron | chemical drug | | | 0.00523 |
| medroxypr | chemical drug | Inhibited | -2.219 | 0.00341 |
| methylpred | chemical drug | | -0.137 | 0.0000229 |
| desogestre | chemical drug | | | 0.0233 |
| aldosterone | chemical - endogen | | -1.159 | 0.00469 |
| tibolone | chemical drug | | | 0.0156 |
| promegeste | chemical drug | | | 0.000418 |
| 27-hydroxy | chemical - endogen | | | 0.00451 |
| metribolon | chemical reagent | Inhibited | -2.236 | 0.0888 |
| U18666A | chemical reagent | | | 0.006 |
| dihydrotest | chemical - endogen | | -1.662 | 0.00000206 |
| spironolact | chemical drug | | | 0.0243 |
| cholesterol | chemical - endogen | | 0.65 | 0.00194 |
| mifepristor | chemical drug | | 0.563 | 0.00216 |
| U73122 | chemical reagent | | | 0.0448 |
| eplerenone | chemical drug | | | 0.023 |
| 24-hydroxy | chemical - endogen | | | 0.006 |
| mibolerone | chemical drug | | -1 | 0.0224 |

| | | | |
|---|---|---|---|
| testosteror | chemical - endogen | -0.251 | 0.00744 |
| progestero | chemical - endogen | -0.196 | 0.000376 |
| phosphate | chemical - endogen | 0 | 0.00575 |
| TCF | group | | 0.00575 |

### Supplementary Table 5. The transcription factor list enriched in pDKO tumor cells

| Term | Overlap | P-value | Old P-valu | Old Adjust | Odds Ratic | Combined | Genes |
|---|---|---|---|---|---|---|---|
| EGR1 CHEA | 10/315 | 0.001472 | 0 | 0 | 3.311306 | 21.59365 | PIAS4;SLC6A6;NT5E;CLDN3;RGS3;CALD1;AKT1;ADAR;PDHB;FKBP5 |
| TCF3 CHEA | 21/1006 | 0.001416 | 0 | 0 | 2.20367 | 14.4561 | TLE3;RARG;B4GALT1;ANP32A;CSNK1A1;TMOD3;NR2F2;SLC1A5;PDHB;LPP;FOXP1;NT5E;CLDN3;CSRP1;FUT9;ST5;KIT;SLCO2A1;TYRO3;RHOU;SGK1 |
| FOXA1 ENCODE | 6/205 | 0.018461 | 0 | 0 | 2.999464 | 11.97408 | CALD1;DAPK3;AKT1S1;MISP;WASF2;FOXP1 |
| ZKSCAN1 ENCODE | 4/120 | 0.034069 | 0 | 0 | 3.410328 | 11.52473 | GM2A;TYRO3;WASF2;DUSP7 |
| **TP53 CHEA** | 8/319 | 0.0165 | 0 | 0 | 2.5705 | 10.55033 | BTG2;CALD1;SRD5A1;ANKRD11;APBB2;SCAND1;MSI2;IER5 |
| E2F1 CHEA | 17/859 | 0.00675 | 0 | 0 | 2.057537 | 10.28409 | KDM6B;SDC4;ANKRD11;KEAP1;MSI2;SLC1A5;DIAPH1;KLF5;WNK1;ST5;KIT;CNOT3;AKT1;FARSA;RABAC1;SOX4;MPZL2 |
| ZC3H11A ENCODE | 4/129 | 0.042614 | 0 | 0 | 3.163337 | 9.982164 | PPP1R37;WNT7B;PLCD3;PTPRF |
| FOXP2 ENCODE | 5/178 | 0.035372 | 0 | 0 | 2.864483 | 9.572612 | CSNK1A1;AKT1S1;WASF2;FOXP1;CORO1C |
| PPARD CHEA | 7/285 | 0.02669 | 0 | 0 | 2.50758 | 9.08609 | PTDSS2;RNF10;ANXA1;RGS3;PMM2;KIT;PCOLCE |
| KLF4 CHEA | 18/987 | 0.011919 | 0 | 0 | 1.89052 | 8.374211 | EIF4A1;RARG;ANP32A;CSNK1A1;WNT7B;TMOD3;SLC1A5;CORO1C;TAPBP;PYGO2;TUBB2B;KLF5;CSRP1;KCNS3;KIF26B;AKT1S1;SGK1;PLEC |
| CHD1 ENCODE | 13/655 | 0.016576 | 0 | 0 | 2.041441 | 8.369533 | KDM6B;CSNK1A1;SIGMAR1;ATXN7L3B;DIAPH1;POR;AKT2;CNOT3;AKT1S1;SCAND1;IER5;WASF2;RBM5 |
| REST CHEA | 22/1280 | 0.010907 | 0 | 0 | 1.791223 | 8.093363 | KCNK5;TLE3;RAMP2;PDCD6IP;KLF13;WDR1;BIK;SRD5A1;WNT7B;GP2;KCNJ15;NR2F2;MSI2;BTBD9;KLF5;WDR92;BRE;WNK1;SGK1;RABAC1;MISP;SOX4 |
| FOSL2 ENCODE | 5/196 | 0.049907 | 0 | 0 | 2.592152 | 7.770207 | CTTN;STX3;PKP3;MISP;WASF2 |
| UBTF ENCODE | 26/1631 | 0.014466 | 0 | 0 | 1.664966 | 7.052775 | RNF10;B4GALT1;ANP32A;ENDOD1;KEAP1;ATXN7L3B;GNAI2;CORO1C;ABHD12;SGPL1;DSTYK;NFS1;TMED2;SCAND1;IER5;YKT6;TLE3;KLF13;TSC22D3;DUSP7;KLF5;ARL2BP;WNK1;TYRO3;TBPL1;FKBP5 |
| TCF3 ENCODE | 15/840 | 0.02462 | 0 | 0 | 1.834816 | 6.796555 | TLE3;B4GALT1;KLF13;CSNK1A1;ANKRD11;ASPSCR1;AK3;NBEAL2;SGPL1;PPP3R1;DSTYK;ALCAM;RHOU;APBB2;FKBP5 |
| NANOG CHEA | 11/595 | 0.040919 | 0 | 0 | 1.884837 | 6.024245 | TLE3;RARG;TUBB2B;B4GALT1;KLF5;ANP32A;FUT9;KIT;SGK1;LPP;FKBP5 |
| NFE2L2 CHEA | 17/1022 | 0.031493 | 0 | 0 | 1.709003 | 5.909738 | ANXA1;GCNT1;ATXN7L3B;NR2F2;LPP;FOXP1;ABHD12;MAPK8;MEIS1;CLDN3;KLF5;BRE;ST5;KIF26B;SLC25A30;RHOU;SGK1 |
| MYOD1 ENCODE | 4/166 | 0.089134 | 0 | 0 | 2.436255 | 5.889936 | PLA2G15;TUBB2B;SDC4;SCAMP2 |
| ZBTB7A ENCODE | 31/2184 | 0.034149 | 0 | 0 | 1.47702 | 4.987914 | ANP32A;ANKRD11;CXXC1;KEAP1;AK3;MSI2;PDHB;PTDSS2;PPP3R1;DRAM1;AKT2;SH3BP2;STX3;RABAC1;WASF2;PIAS4;TMEM120A;ASPSCR1;NBEAL2;EMP2;CYBA;SCAMP2;FOXP1;PLA2G15;PYGO2;TUBB2B;CTTN;AKT1S1;TBPL1;PLCD3;PLEC |
| SP1 ENCODE | 12/707 | 0.057349 | 0 | 0 | 1.7268 | 4.936216 | EIF4A1;POR;RNF114;ANP32A;ANKRD11;CNOT3;HJURP;SUCO;CHP1;BAZ1B;WASF2;DNM2 |
| TP63 CHEA | 21/1399 | 0.0466 | 0 | 0 | 1.542285 | 4.728872 | DDR1;TMEM45B;GSDMC;PFKFB4;ANXA1;WDR1;BIK;KCNJ15;EMP2;SLC1A5;CORO1C;PTDSS2;MMP14;MEIS1;ALCAM;ST5;KIF26B;PKP1;STOM;S100A14;SGK1 |
| FLI1 ENCODE | 11/650 | 0.068122 | 0 | 0 | 1.717674 | 4.614449 | NAA50;CAP1;MFSD8;SEC61A1;BRE;DAPK3;CWC22;HJURP;TBPL1;FARSA;EIF3C |
| SMAD4 CHEA | 10/584 | 0.07498 | 0 | 0 | 1.73521 | 4.495108 | TLE3;BTG2;MEIS1;ALCAM;ST5;KCNS3;TMOD3;SUCO;PKP1;XDH |
| RAD21 ENCODE | 19/1265 | 0.056523 | 0 | 0 | 1.537394 | 4.417089 | NAA50;TSC22D3;WNT7B;NR2F2;PTPRF;DNM2;PTDSS2;DIAPH1;SGPL1;PPP3R1;RGS3;WNK1;KIF26B;TYRO3;APBB2;STX3;RABAC1;PLCD3;PLEC |
| FOXA2 ENCODE | 6/316 | 0.103561 | 0 | 0 | 1.914557 | 4.341435 | B4GALT1;KLF5;DAPK3;NR2F2;MISP;FOXP1 |
| CBX3 ENCODE | Feb-81 | 0.198719 | 0 | 0 | 2.483532 | 4.013049 | DSTYK;CHP1 |
| ZMIZ1 ENCODE | 14/914 | 0.082687 | 0 | 0 | 1.555309 | 3.876907 | CAP1;EIF4A1;BTG2;KLF13;CSNK1A1;DAPK3;AK3;SLC1A5;GNAI2;PYGO2;NFS1;SCAND1;IER5;PLEC |
| GATA2 CHEA | 12/772 | 0.095122 | 0 | 0 | 1.573739 | 3.702364 | DDR1;TLE3;BTG2;CCDC91;RGS3;WDR1;GNA12;PKP1;ADAR;SLC1A5;IER5;PTPRF |
| SMC3 ENCODE | 17/1181 | 0.092899 | 0 | 0 | 1.463072 | 3.476608 | TSC22D3;WNT7B;CXXC1;AK3;PDHB;PTPRF;DIAPH1;SGPL1;RGS3;WNK1;KIF26B;TYRO3;S100A14;STX3;RABAC1;PLCD3;PLEC |
| FOS ENCODE | 10/637 | 0.11509 | 0 | 0 | 1.584154 | 3.425007 | EIF4A1;TLE3;KLF5;DAPK3;CNOT3;HJURP;SUCO;BAZ1B;WASF2;DNM2 |
| PBX3 ENCODE | 18/1269 | 0.094829 | 0 | 0 | 1.442427 | 3.397888 | EIF4A1;TLE3;PFKFB4;ANP32A;PDPK1;ANKRD11;DAPK3;BAZ1B;SCAMP2;DNM2;AKAP13;POR;DSTYK;FAM83E;KLF5;CNOT3;GNA12;IER5 |
| CTCF ENCODE | 24/1790 | 0.097178 | 0 | 0 | 1.368949 | 3.191311 | NAA50;TLE3;TSC22D3;WNT7B;CXXC1;PTPRF;CORO1C;MFSD8;SLC6A6;DIAPH1;SGPL1;PPP3R1;ALCAM;WNK1;KIF26B;TYRO3;CHP1;APBB2;SCAND1;STX3;RABAC1;PLCD3;YKT6;PLEC |
| TCF7L2 ENCODE | 9/583 | 0.140164 | 0 | 0 | 1.553639 | 3.052811 | NAA50;TLE3;ANP32A;CSNK1A1;AKT2;CXXC1;AK3;SAMHD1;WASF2 |
| ATF2 ENCODE | 36/2852 | 0.095618 | 0 | 0 | 1.29992 | 3.051425 | PPP1R37;EIF4A1;HBS1L;BTG2;SRCAP;WDR1;CXXC1;ADAR;SLC1A5;MFSD8;PTDSS2;PPP3R1;DSTYK;FAM83E;WDR92;TMED2;CHP1;SCAND1;IER5;SOX4;NAA50;CAP1;TLE3;PDCD6IP;DAPK3;SUCO;BAZ1B;DNM2;DIAPH1;POR;BRE;WNK1;FUBP1;AKT1S1;TBPL1;PLCD3 |
| ELF1 ENCODE | 31/2483 | 0.129743 | 0 | 0 | 1.274932 | 2.603666 | EIF4A1;RNF10;WDR1;ANKRD11;CXXC1;AK3;ATXN7L3B;SLC1A5;PPP3R1;WDR92;TMED2;PIP4K2C;SCAND1;YKT6;WASF2;CAP1;PDCD6IP;CSNK1A1;SRD5A1;PDPK1;DAPK3;PMM2;ASPSCR1;SCAMP2;DNM2;PLA2G15;DIAPH1;MEIS1;ARL2BP;CTTN;TBPL1 |
| PPARG CHEA | 8/535 | 0.178397 | 0 | 0 | 1.500122 | 2.585822 | POR;CCDC91;RNF114;TMEM120A;TBPL1;CHP1;SLC1A5;PTRF |
| SOX2 CHEA | 11/775 | 0.165918 | 0 | 0 | 1.427267 | 2.56375 | TLE3;TUBB2B;ANP32A;CSRP1;CALD1;TMOD3;KIT;RHOU;MSLN;LPP;FOXP1 |
| EGR1 ENCODE | 8/537 | 0.180877 | 0 | 0 | 1.494295 | 2.555152 | PLA2G15;DIAPH1;DSTYK;KLF13;BRE;TSC22D3;KEAP1;ADAR |
| NFYB ENCODE | 44/3715 | 0.146971 | 0 | 0 | 1.21562 | 2.330973 | TMEM45B;EIF4A1;ANP32A;WDR1;ANKRD11;HJURP;SLC1A5;GNAI2;CORO1C;PTDSS2;AKAP13;DRAM1;WDR92;GM2A;GNA12;TMED2;CHP1;STOM;IER5;PTRF;WASF2;SOX4;RBM5;PIAS4;TLE3;MBNL2;ANXA1;KLF13;TMEM120A;DAPK3;SIGMAR1;SUCO;BAZ1B;SCAMP2;DUSP7;DNM2;PLA2G15;MEIS1;KLF5;ARL2BP;CNOT3;AKT1S1;TBPL1;SERINC3 |
| USF2 ENCODE | 13/965 | 0.183342 | 0 | 0 | 1.354406 | 2.29762 | NAA50;PIAS4;EIF4A1;PDCD6IP;PMM2;ASPSCR1;ADAR;DUSP7;PPP3R1;AKT1S1;PIP4K2C;IER5;FKBP5 |
| STAT3 CHEA | 3/177 | 0.267916 | 0 | 0 | 1.691638 | 2.228028 | AKAP13;APBB2;LPP |
| STAT3 ENCODE | 10/725 | 0.203298 | 0 | 0 | 1.382804 | 2.202919 | NAA50;DIAPH1;BTG2;B4GALT1;KLF5;ARL2BP;FUBP1;DAPK3;TMED2;PDHB |
| CEBPD ENCODE | 10/734 | 0.213661 | 0 | 0 | 1.36497 | 2.106648 | EIF4A1;PPP3R1;BTG2;PFKFB4;B4GALT1;DRAM1;WDR1;RSRP1;SLC1A5;IER5 |
| YY1 CHEA | 5/332 | 0.24862 | 0 | 0 | 1.503568 | 2.092708 | PIAS4;DIAPH1;AKT1S1;CXXC1;FARSA |
| USF1 ENCODE | 18/1423 | 0.19735 | 0 | 0 | 1.27366 | 2.066862 | NAA50;PIAS4;EIF4A1;TLE3;PDCD6IP;SRD5A1;ASPSCR1;ADAR;DUSP7;DIAPH1;PPP3R1;CLCN4;CLIP4;AKT1S1;PIP4K2C;STX3;IER5;FKBP5 |
| MYC CHEA | 8/573 | 0.227944 | 0 | 0 | 1.396469 | 2.064895 | KCNK5;EIF4A1;PTDSS2;KLF5;ANP32A;FUT9;EHMT2;FUBP1 |
| POU5F1 CHEA | 4/261 | 0.273808 | 0 | 0 | 1.528264 | 1.979606 | DIAPH1;RARG;MEIS1;SGK1 |
| NFYA ENCODE | 27/2250 | 0.203989 | 0 | 0 | 1.212781 | 1.927945 | EIF4A1;ANP32A;ANKRD11;HJURP;SLC1A5;GNAI2;AKAP13;GM2A;GNA12;CHP1;IER5;WASF2;SOX4;TLE3;MBNL2;KLF13;TMEM120A;DAPK3;SUCO;BAZ1B;SCAMP2;DUSP7;DNM2;PLA2G15;KLF5;ARL2BP;CNOT3 |
| SPI1 ENCODE | 19/1540 | 0.219121 | 0 | 0 | 1.24076 | 1.883637 | PPP1R37;CAP1;EIF4A1;RNF10;ANP32A;CSNK1A1;SRD5A1;NR2F2;GNAI2;CORO1C;POR;PPP3R1;PDLIM2;ARL2BP;CTTN;GM2A;CWC22;IER5;PLEC |
| BHLHE40 ENCODE | 5/348 | 0.279384 | 0 | 0 | 1.432253 | 1.826361 | EIF4A1:PDCD6IP:TMEM120A:ASPSCR1:PDHB |
| NR2C2 ENCODE | 5/350 | 0.283285 | 0 | 0 | 1.423803 | 1.795844 | NAA50;PLA2G15;NFS1;IKZF2;BAZ1B |
| RUNX1 CHEA | 16/1294 | 0.241724 | 0 | 0 | 1.239838 | 1.76052 | PIAS4;BTG2;KLF13;WDR1;TMOD3;BTBD9;SAMHD1;PTPRF;GNAI2;CTTN;WNK1;AKT1S1;SH3BP2;IER5;SOX4;PLEC |
| BCL3 ENCODE | 4/273 | 0.300972 | 0 | 0 | 1.459192 | 1.752108 | PPP1R37;ALCAM;FAM83E;CYBA |
| IRF8 CHEA | 2/121 | 0.348177 | 0 | 0 | 1.645387 | 1.735955 | PDLIM2;TAPBP |
| RCOR1 ENCODE | 9/702 | 0.284653 | 0 | 0 | 1.278886 | 1.606901 | ANP32A;PDPK1;TMOD3;TMEM120A;CNOT3;CXXC1;MSI2;FARSA;DUSP7 |
| TAF1 ENCODE | 38/3346 | 0.248243 | 0 | 0 | 1.147965 | 1.599512 | EIF4A1;HBS1L;BTG2;SRCAP;WDR1;ANKRD11;ATXN7L3B;SLC1A5;GNAI2;CORO1C;ABHD12;MFSD8;SEC61A1;DSTYK;CWC22;TMED2;CHP1;SCAND1;IER5;YKT6;WASF2;RBM5;CAP1;PIAS4;TLE3;PDCD6IP;CSNK1A1;SRD5A1;ASPSCR1;BAZ1B;PYGO2;POR;WNK1;FUBP1;AKT1S1;RSRP1;TBPL1;FARSA |
| GATA1 CHEA | 10/807 | 0.304631 | 0 | 0 | 1.235202 | 1.468227 | CAP1;BTG2;MEIS1;PPP1R14C;WDR92;KIT;GNA12;ENDOD1;IER5;PTPRF |
| CREB1 CHEA | 17/1444 | 0.297788 | 0 | 0 | 1.176579 | 1.425278 | PIAS4;KDM6B;EIF4A1;HBS1L;TLE3;BTG2;ANKRD11;CXXC1;ADAR;BTBD9;DUSP7;DIAPH1;BRE;CNOT3;CHP1;RABAC1;PLEC |
| SP2 ENCODE | 12/994 | 0.310101 | 0 | 0 | 1.203762 | 1.409433 | EIF4A1;AKAP13;KLF5;ANP32A;ANKRD11;DAPK3;CNOT3;GNA12;TBPL1;BAZ1B;WASF2;DNM2 |
| ZNF384 ENCODE | 9/730 | 0.323669 | 0 | 0 | 1.227419 | 1.38457 | NAA50;TLE3;RNF10;GM2A;CSNK1A1;AKT2;PMM2;SUCO;AK3 |
| SRF ENCODE | 4/299 | 0.36065 | 0 | 0 | 1.328814 | 1.355186 | CAP1;B4GALT1;WDR1;DAPK3 |
| SIX5 ENCODE | 13/1094 | 0.319344 | 0 | 0 | 1.184615 | 1.352222 | SRCAP;TMOD3;PMM2;CXXC1;SUCO;DNM2;PTDSS2;DSTYK;WNK1;CWC22;AKT1S1;FARSA;RABAC1 |
| RFX5 ENCODE | 7/559 | 0.340422 | 0 | 0 | 1.245148 | 1.341733 | AKAP13;MBNL2;CNOT3;GNA12;CHP1;SOX4;GNAI2 |
| IRF3 ENCODE | 8/663 | 0.360112 | 0 | 0 | 1.198951 | 1.224536 | EIF4A1;KLF5;ANKRD11;CNOT3;GNA12;BAZ1B;WASF2;DNM2 |
| SUZ12 CHEA | 19/1684 | 0.349211 | 0 | 0 | 1.12452 | 1.183083 | RAMP2;PPP1R14C;WNT7B;NR2F2;FUT4;DUSP7;NT5E;MEIS1;ALCAM;PDLIM2;ADAMTS1;KCNS3;KIF26B;SLCO2A1;PKP1;MAL;IER5;PTRF;SOX4 |
| CEBPB ENCODE | 2/144 | 0.430594 | 0 | 0 | 1.377269 | 1.160474 | EIF4A1;DRAM1 |
| ERG CHEA | 2/144 | 0.430594 | 0 | 0 | 1.377269 | 1.160474 | AK3;SCAMP2 |
| MAX ENCODE | 23/2073 | 0.357905 | 0 | 0 | 1.106182 | 1.136588 | NAA50;PIAS4;EIF4A1;TLE3;PDCD6IP;SRD5A1;PMM2;SIGMAR1;ASPSCR1;ADAR;SLC1A5;PDHB;DUSP7;PTDSS2;SLC6A6;DIAPH1;SEC61A1;KLF5;WDR92;BRE;AKT1S1;CHP1;SCAND1 |
| NELFE ENCODE | 3/234 | 0.424838 | 0 | 0 | 1.270519 | 1.087625 | KCNK5;SLC1A5;IER5 |
| EZH2 CHEA | 3/237 | 0.432897 | 0 | 0 | 1.254038 | 1.049952 | WNT7B;KIF26B;PTRF |
| ESR1 CHEA | 2/154 | 0.464711 | 0 | 0 | 1.286004 | 0.985517 | SLCO2A1;XDH |
| YY1 ENCODE | 29/2753 | 0.445128 | 0 | 0 | 1.044604 | 0.845496 | EIF4A1;HBS1L;RNF10;SRCAP;ENDOD1;CXXC1;KEAP1;PDHB;ABHD12;MFSD8;DSTYK;RNF114;CWC22;SCAND1;RABAC1;WASF2;RBM5;NAA50;PIAS4;TLE3;CSNK1A1;PDPK1;ASPSCR1;BAZ1B;WNK1;FUBP1;CNOT3;AKT1S1;FARSA |
| PML ENCODE | 17/1596 | 0.454656 | 0 | 0 | 1.054519 | 0.831186 | CAP1;PIAS4;EIF4A1;SRCAP;WDR1;KEAP1;CYBA;SLC1A5;PYGO2;DIAPH1;POR;DSTYK;RSRP1;TMED2;SCAND1;IER5;YKT6 |
| HNF4A ENCODE | 13/1276 | 0.535533 | 0 | 0 | 1.004051 | 0.627023 | B4GALT1;WDR1;CSNK1A1;TMEM120A;AK3;LTBP3;BAZ1B;SAMHD1;PDLIM2;BRE;AKT2;SH3BP2;SCAND1 |
| NRF1 ENCODE | 19/1882 | 0.545279 | 0 | 0 | 0.994031 | 0.602838 | NAA50;PIAS4;HBS1L;PFKFB4;PDCD6IP;SRCAP;CSNK1A1;SRD5A1;SUCO;ATXN7L3B;PDHB;SAMHD1;MFSD8;ARL2BP;ST5;AKT2;AKT1S1;RSRP1;SCAND1 |
| MYC ENCODE | 15/1515 | 0.578577 | 0 | 0 | 0.973245 | 0.532544 | EIF4A1;TLE3;SRD5A1;PMM2;ASPSCR1;SLC1A5;BAZ1B;SLC6A6;DIAPH1;WDR92;FUBP1;AKT1S1;SCAND1;FARSA;FKBP5 |
| FOXM1 ENCODE | Jan-95 | 0.621474 | 0 | 0 | 1.037655 | 0.493572 | HJURP |
| E2F6 ENCODE | 32/3245 | 0.600675 | 0 | 0 | 0.965901 | 0.49232 | KCNK5;EIF4A1;B4GALT1;KEAP1;SLC1A5;PDHB;FUT4;SEC61A1;DRAM1;CSRP1;SH3BP2;CHP1;STOM;SCAND1;IER5;NAA50;TLE3;PMM2;SIGMAR1;ASPSCR1;EMP2;BAZ1B;DUSP7;FOXP1;SLC6A6;DIAPH1;KLF5;CTTN;WNK1;KIF26B;SLC25A30;PLEC |
| GABPA ENCODE | 20/2082 | 0.637155 | 0 | 0 | 0.939986 | 0.423691 | NAA50;CAP1;EIF4A1;PDCD6IP;CSNK1A1;PDPK1;DAPK3;PMM2;PYGO2;DIAPH1;ERLEC1;DSTYK;NFS1;BRE;CWC22;AKT1S1;TBPL1;SCAND1;FARSA;YKT6 |
| ZEB1 ENCODE | 1/106 | 0.661847 | 0 | 0 | 0.92843 | 0.383182 | STX3 |
| AR CHEA | 10/1095 | 0.679247 | 0 | 0 | 0.893579 | 0.345611 | ANP32A;BRE;FUT9;ST5;RHOU;NR2F2;IKZF2;LPP;PTPRF;FKBP5 |
| ATF3 ENCODE | 1/114 | 0.68849 | 0 | 0 | 0.86235 | 0.321876 | PDCD6IP |
| SALL4 CHEA | 3/355 | 0.701803 | 0 | 0 | 0.828622 | 0.293417 | MEIS1;KLF5;SDC4 |
| SIN3A ENCODE | 10/1131 | 0.717581 | 0 | 0 | 0.863219 | 0.286476 | EIF4A1;DIAPH1;POR;WDR92;BRE;FUBP1;PMM2;HJURP;TMED2;SLC1A5 |
| SPI1 CHEA | 9/1056 | 0.75042 | 0 | 0 | 0.830798 | 0.23854 | KDM6B;EIF4A1;SLC6A6;POR;PPP3R1;RNF114;CSNK1A1;CWC22;STOM |
| IRF1 ENCODE | 2/263 | 0.749349 | 0 | 0 | 0.744782 | 0.214907 | FOXP1;DNM2 |
| BCLAF1 ENCODE | 7/851 | 0.765421 | 0 | 0 | 0.802006 | 0.2144 | CAP1;DSTYK;FAM83E;PDCD6IP;TBPL1;CHP1;SCAND1 |
| BRCA1 ENCODE | 29/3218 | 0.78568 | 0 | 0 | 0.867984 | 0.209363 | PFKFB4;SRCAP;ANKRD11;CORO1C;MFSD8;SEC61A1;PPP3R1;DSTYK;ALCAM;TMED2;APBB2;SCAND1;NAA50;TLE3;CSNK1A1;DAPK3;SUCO;BAZ1B;BTBD9;PYGO2;DIAPH1;POR;ARL2BP;BRE;WNK1;FUBP1;AKT1S1;PLCD3;FKBP5 |
| TAF7 ENCODE | 5/640 | 0.781831 | 0 | 0 | 0.76203 | 0.187548 | WDR1;KEAP1;SLC1A5;IER5;CORO1C |
| CREB1 ENCODE | 19/2238 | 0.826665 | 0 | 0 | 0.81799 | 0.155709 | EIF4A1;TLE3;WDR1;ANKRD11;DAPK3;CXXC1;SUCO;ADAR;MSI2;BTBD9;DNM2;WDR92;BRE;CWC22;AKT1S1;TMED2;CHP1;RABAC1;SERINC3 |
| ETS1 ENCODE | 1/156 | 0.797642 | 0 | 0 | 0.62734 | 0.141839 | CXXC1 |
| ZBTB33 ENCODE | 5/702 | 0.84429 | 0 | 0 | 0.691999 | 0.117127 | MFSD8;SEC61A1;SRCAP;FUBP1;TMED2 |
| TRIM28 CHEA | 1/210 | 0.883947 | 0 | 0 | 0.463973 | 0.057235 | KLF5 |
| REST ENCODE | 2/383 | 0.903183 | 0 | 0 | 0.507071 | 0.051635 | DAPK3;TMED2 |
| STAT5A ENCODE | 1/233 | 0.90846 | 0 | 0 | 0.417485 | 0.04008 | WDR92 |
| RELA ENCODE | 2/484 | 0.958927 | 0 | 0 | 0.398732 | 0.016723 | DRAM1;IER5 |
| E2F4 ENCODE | 3/710 | 0.976898 | 0 | 0 | 0.405021 | 0.009467 | WNK1;HJURP;BAZ1B |
| EZH2 ENCODE | 1/338 | 0.969122 | 0 | 0 | 0.285865 | 0.008966 | BIK |

### Supplementary Table 6. The transcription factor list enriched in pTKO tumor cells

| Term | Overlap | P-value | Old P-value | Old Adjuste | Odds Ratio | Combined | Genes |
|---|---|---|---|---|---|---|---|
| POU5F1 CHEA | 5/261 | 0.018727 | 0 | 0 | 3.422677 | 13.61474 | SFRP1;SPRED1;IL6ST;DUSP6;ETV5 |
| NANOG CHEA | 9/595 | 0.008267 | 0 | 0 | 2.74417 | 13.15963 | SFRP1;SPRED1;KRT1;SFTPD;SLC2A3;IL6ST;PVRL3;DUSP6;ETV5 |
| SOX2 CHEA | 10/775 | 0.015544 | 0 | 0 | 2.335593 | 9.72553 | SFRP1;SPRED1;APOC1;RNASE4;FBLN1;IL6ST;PVRL3;FBP1;DUSP6;ETV5 |
| FOXA2 ENCODE | 5/316 | 0.038239 | 0 | 0 | 2.809486 | 9.169913 | RASSF10;VNN3;CXCL2;DUSP6;ETV5 |
| SUZ12 CHEA | 17/1684 | 0.018591 | 0 | 0 | 1.85766 | 7.402876 | TBX1;ACE;BCL11B;LPAR3;ARHGAP6;SLC16A14;CH25H;THBD;SLC5A8;SFRP1;CCND2;MFSD4;RASSF10;CHST15;PROK2;PDZRN3;FBP1 |
| AR CHEA | 10/1095 | 0.107967 | 0 | 0 | 1.619191 | 3.604201 | THBD;SLC5A8;SFRP1;SPRED1;BCL11B;LPAR3;PDZRN3;FBP1;FAM149A;SLC16A14 |
| MYOD1 ENCODE | 2/166 | 0.253522 | 0 | 0 | 2.091092 | 2.869614 | TMEM119;PBXIP1 |
| PPARD CHEA | 3/285 | 0.233253 | 0 | 0 | 1.829134 | 2.662546 | LANCL1;THEM5;GRHL3 |
| STAT3 CHEA | 2/177 | 0.277363 | 0 | 0 | 1.958559 | 2.511714 | IL6ST;ETV5 |
| VDR CHEA | Jan-76 | 0.360293 | 0 | 0 | 2.276782 | 2.324223 | SP110 |
| FOXA1 ENCODE | 2/205 | 0.337671 | 0 | 0 | 1.686014 | 1.830477 | DUSP6;ETV5 |
| SALL4 CHEA | 3/355 | 0.344313 | 0 | 0 | 1.460153 | 1.556821 | SPRED1;SFTPD;IL6ST |
| TP63 CHEA | 10/1399 | 0.302257 | 0 | 0 | 1.244357 | 1.488846 | PFKFB3;MFSD4;RASSF10;KRT1;CHST15;LPAR3;GRHL3;KRT5;PDZRN3;DSC2 |
| KLF4 CHEA | 7/987 | 0.356345 | 0 | 0 | 1.227468 | 1.266571 | SFRP1;GABBR1;ATP10D;APOC1;SLC2A3;S1PR3;IL6ST |
| NFE2L2 CHEA | 7/1022 | 0.390111 | 0 | 0 | 1.182947 | 1.113535 | PHF3;SPRED1;RASSF6;CHST15;S1PR3;PDZRN3;ASGR1 |
| ZNF384 ENCODE | 5/730 | 0.42495 | 0 | 0 | 1.17968 | 1.00955 | SPRED1;CCND2;APOC1;SLC2A3;ETV5 |
| IRF8 CHEA | 1/121 | 0.509373 | 0 | 0 | 1.419756 | 0.95773 | RNASE4 |
| SMAD4 CHEA | 4/584 | 0.446817 | 0 | 0 | 1.17809 | 0.949078 | RASSF10;KRT13;PIP5K1B;DUSP6 |
| NFIC ENCODE | 2/279 | 0.487391 | 0 | 0 | 1.230953 | 0.884672 | SPRED1;WDFY1 |
| ZC3H11A ENCODE | 1/129 | 0.532008 | 0 | 0 | 1.330482 | 0.839663 | ASGR1 |
| EGR1 CHEA | 2/315 | 0.552427 | 0 | 0 | 1.087373 | 0.645283 | LPAR3;ETV5 |
| TCF3 CHEA | 6/1006 | 0.539847 | 0 | 0 | 1.020703 | 0.629231 | SFRP1;SPRED1;CHST15;IL6ST;DUSP6;ETV5 |
| TP53 CHEA | 2/319 | 0.559291 | 0 | 0 | 1.073433 | 0.623757 | CCND2;BCL11B |
| PPARG CHEA | 3/535 | 0.609226 | 0 | 0 | 0.957212 | 0.474362 | RNASE4;PDZRN3;DUSP6 |
| STAT3 ENCODE | 4/725 | 0.617045 | 0 | 0 | 0.940778 | 0.45422 | SP110;VNN3;RNASE4;IL6ST |
| RUNX1 CHEA | 7/1294 | 0.639055 | 0 | 0 | 0.919489 | 0.411715 | CD72;RNASE4;GRHL3;IL6ST;ARHGAP6;DUSP6;B4GALNT4 |
| RFX5 ENCODE | 3/559 | 0.638952 | 0 | 0 | 0.914758 | 0.409744 | CCND2;CD72;APOC1 |
| FOSL2 ENCODE | 1/196 | 0.685139 | 0 | 0 | 0.87038 | 0.32912 | RNASE4 |
| GATA1 CHEA | 4/807 | 0.699917 | 0 | 0 | 0.841094 | 0.300097 | SLC14A1;RASSF10;PIP5K1B;SLC2A3 |
| TRIM28 CHEA | 1/210 | 0.710212 | 0 | 0 | 0.8115 | 0.277688 | SFRP1 |
| SMC3 ENCODE | 5/1181 | 0.827653 | 0 | 0 | 0.710148 | 0.134332 | THBD;SELL;SLC16A14;ASGR1;CTHRC1 |
| UBTF ENCODE | 7/1631 | 0.850145 | 0 | 0 | 0.715478 | 0.116157 | SPRED1;CCND2;PFKFB3;ATP10D;PIP5K1B;IL6ST;PFN2 |
| SRF ENCODE | 1/299 | 0.829244 | 0 | 0 | 0.566564 | 0.106084 | PFKFB3 |
| HNF4A ENCODE | 5/1276 | 0.87441 | 0 | 0 | 0.653732 | 0.087734 | FBLN1;DSC2;ASGR1;DUSP6;ETV5 |
| TCF7L2 ENCODE | 2/583 | 0.85912 | 0 | 0 | 0.577774 | 0.087733 | CHST15;DUSP6 |
| SPI1 CHEA | 4/1056 | 0.871637 | 0 | 0 | 0.633635 | 0.087051 | CD72;SP110;ARHGAP6;DUSP6 |
| TCF3 ENCODE | 3/840 | 0.874138 | 0 | 0 | 0.598818 | 0.080551 | PBXIP1;IL6ST;ETV5 |
| EZH2 ENCODE | 1/338 | 0.864671 | 0 | 0 | 0.5 | 0.072703 | B4GALNT4 |
| BHLHE40 ENCODE | 1/348 | 0.872513 | 0 | 0 | 0.485342 | 0.06619 | SPRED1 |
| FOS ENCODE | 2/637 | 0.890816 | 0 | 0 | 0.527162 | 0.060949 | IL6ST;ETV5 |
| RCOR1 ENCODE | 2/702 | 0.920229 | 0 | 0 | 0.476596 | 0.039621 | SPRED1;IL6ST |
| CEBPD ENCODE | 2/734 | 0.931827 | 0 | 0 | 0.455001 | 0.032127 | SPRED1;CXCL2 |
| PBX3 ENCODE | 4/1269 | 0.944034 | 0 | 0 | 0.520984 | 0.030005 | SPRED1;WDFY1;IL6ST;ETV5 |
| SPI1 ENCODE | 5/1540 | 0.95226 | 0 | 0 | 0.53362 | 0.026103 | SP110;KRT13;EPHX3;PIP5K1B;DSC2 |
| GATA2 CHEA | 2/772 | 0.943544 | 0 | 0 | 0.431688 | 0.025087 | PIP5K1B;CXCL2 |
| CTCF ENCODE | 6/1790 | 0.956467 | 0 | 0 | 0.548388 | 0.024408 | THBD;SPRED1;SELL;SLC16A14;ASGR1;CTHRC1 |
| RELA ENCODE | 1/484 | 0.943564 | 0 | 0 | 0.346255 | 0.020115 | IL6ST |
| E2F1 CHEA | 2/859 | 0.963618 | 0 | 0 | 0.386099 | 0.014309 | SPRED1;RNASE4 |
| EGR1 ENCODE | 1/537 | 0.958982 | 0 | 0 | 0.311165 | 0.013032 | AP1S3 |
| ZBTB7A ENCODE | Jul-84 | 0.976961 | 0 | 0 | 0.517568 | 0.012064 | SPRED1;APOC1;WDFY1;ASGR1;ETV5;B4GALNT4;CTHRC1 |
| RAD21 ENCODE | 3/1265 | 0.981173 | 0 | 0 | 0.388293 | 0.00738 | SLC14A1;SELL;CTHRC1 |
| USF2 ENCODE | 2/965 | 0.978971 | 0 | 0 | 0.341686 | 0.007262 | SPRED1;PBXIP1 |
| SP2 ENCODE | 2/994 | 0.981939 | 0 | 0 | 0.331189 | 0.006036 | IL6ST;ETV5 |
| FLI1 ENCODE | 1/650 | 0.979287 | 0 | 0 | 0.255486 | 0.005347 | DUSP6 |
| CHD1 ENCODE | 1/655 | 0.979906 | 0 | 0 | 0.253467 | 0.005145 | CCND2 |
| IRF3 ENCODE | 1/663 | 0.980858 | 0 | 0 | 0.2503 | 0.004838 | ETV5 |
| ZBTB33 ENCODE | 1/702 | 0.984895 | 0 | 0 | 0.235895 | 0.00359 | ETV5 |
| SP1 ENCODE | 1/707 | 0.985348 | 0 | 0 | 0.234163 | 0.003456 | ETV5 |
| USF1 ENCODE | 3/1423 | 0.991289 | 0 | 0 | 0.342161 | 0.002994 | SPRED1;WDFY1;PBXIP1 |
| REST CHEA | 2/1280 | 0.996144 | 0 | 0 | 0.253181 | 9.78E-04 | SP110;LPAR3 |
| NFYA ENCODE | May-50 | 0.997828 | 0 | 0 | 0.35074 | 7.62E-04 | LANCL1;GSTT1;PBXIP1;IL6ST;ETV5 |
| ZMIZ1 ENCODE | 1/914 | 0.995867 | 0 | 0 | 0.179118 | 7.42E-04 | CCND2 |
| CREB1 CHEA | 2/1444 | 0.998457 | 0 | 0 | 0.222408 | 3.44E-04 | LANCL1;B4GALNT4 |
| SIX5 ENCODE | 1/1094 | 0.998639 | 0 | 0 | 0.1482 | 2.02E-04 | ARHGAP44 |
| NFYB ENCODE | 15-Aug | 0.99992 | 0 | 0 | 0.320267 | 2.55E-05 | LANCL1;ATP10D;APOC1;GSTT1;PHGDH;PBXIP1;IL6ST;ETV5 |
| E2F6 ENCODE | Jun-45 | 0.999939 | 0 | 0 | 0.277763 | 1.69E-05 | CCND2;PFKFB3;ACE;RNASE4;PFN2;CTHRC1 |
| BRCA1 ENCODE | 18-Mar | 0.999996 | 0 | 0 | 0.136433 | 5.83E-07 | SPRED1;RNASE4;ETV5 |
| MAX ENCODE | Jan-73 | 0.999994 | 0 | 0 | 0.074104 | 4.53E-07 | SPRED1 |
| TAF1 ENCODE | Feb-46 | 0.999996 | 0 | 0 | 0.086015 | 3.45E-07 | IL6ST;ETV5 |
| CREB1 ENCODE | Jan-38 | 0.999996 | 0 | 0 | 0.068002 | 3.02E-07 | APOC1 |
| ELF1 ENCODE | Jan-83 | 0.999996 | 0 | 0 | 0.060439 | 2.32E-07 | SP110 |
| YY1 ENCODE | Jan-53 | 0.999996 | 0 | 0 | 0.053663 | 2.05E-07 | WDFY1 |
| ATF2 ENCODE | Jan-52 | 0.999996 | 0 | 0 | 0.0515 | 1.98E-07 | CCND2 |

#### Supplementary Table 7. The CTL gene signiture used in GSEA in Figure 7

CD3
CD8
PRF1
GNLY
GZMB
GZMK
NKG7
IL2
CXCR6
ITGA1
ANXA2
IFNG
MYO1F
KLRD1
IFIT3
CCL4
XCL1
NKG7

#### Supplementary Table 8. Primers used in qRT-PCR

| Gene name | Forward | Reverse | |
|---|---|---|---|
| Mapk8 | GTGGAATCAAGCACCTTCACT | TCCTCGCCAGTCCAAAATCAA | |
| Kit | GGCCTCACGAGTTCTATTTACG | GGGGAGAGATTTCCCATCACAC | |
| Sox4 | GACAGCGACAAGATTCCGTTC | GTTGCCCGACTTCACCTTC | |
| Ccnd2 | GAGTGGGAACTGGTAGTGTTG | CGCACAGAGCGATGAAGGT | |
| Msln | CTTGGGTGGATACCACGTCTG | GTCCCCGGAGTGTAATGTTCT | |
| Ido1 | GCCTCCTATTCTGTCTTATGCAG | ATACAGTGGGGATTGCTTTGATT | |
| Myc | GAGCTCCTCGAGCTGTTTGA | GCTGTACGGAGTCGTAGTCG | |
| Sp1 | GCCGCCTTTTCTCAGACTC | TTGGGTGACTCAATTCTGCTG | |
| Srd5A1 | CTTGAGCCAGTTTGCGGTGTA | GCCTCCCCTGGGTATTTTGTATC | |
| Cd68 | TGTCTGATCTTGCTAGGACCG | GAGAGTAACGGCCTTTTTGTGA | |
| Cd40 | TGTCATCTGTGAAAAGGTGGTC | ACTGGAGCAGCGGTGTTATG | |
| Anxa1 | ATGTATCCTCGGATGTTGCTGC | TGAGCATTGGTCCTCTTGGTA | |
| Gilz | CATGGAGGTGGCGGTCTATC | CACCTCCTCTCTCACAGCGT | |
| L-Gilz | ACCGCAACATAGACCAGACC | CACAGCGTACATCAGGTGGT | |
| Saa3 | TGCCATCATTCTTTGCATCTTGA | CCGTGAACTTCTGAACAGCCT | |
| Msi2 | CAGAGGCTTCGGTTTCGTCA | TTGGGTCAATCGTCTTGGAATC | |
| Trp53 | GTCACAGCACATGACGGAGG | TCTTCCAGATGCTCGGGATAC | |
| Ptges | GGATGCGCTGAAACGTGGA | CAGGAATGAGTACACGAAGCC | |
| S100A8 | AAATCACCATGCCCTCTACAAG | CCCACTTTTATCACCATCGCAA | |
| Csf1 | GTGTCAGAACACTGTAGCCAC | TCAAAGGCAATCTGGCATGAAG | |
| Tnf | CCTGTAGCCCACGTCGTAG | GGGAGTAGACAAGGTACAACCC | |
| Mmp9 | GCAGAGGCATACTTGTACCG | TGATGTTATGATGGTCCCACTTG | |
| C3 | CCAGCTCCCCATTAGCTCTG | GCACTTGCCTCTTTAGGAAGTC | |
| Mst1 | CTCACCACTGAATGACTTCCAG | AAGGCCCGACAGTCCAGAA | |
| Pik3Ip1 | CCCAGAGACCACTTCCCAAG | TGGTAGGGCGTTAGCAGGA | |
| Nfkbie | GAGGCGCTCACATACATCTCT | GCTGTCTGGTAAAGGTTGTTCTG | |
| Ikbke | ACCACTAACTACCTGTGGCAT | ACTGCGAATAGCTTCACGATG | |
| Axin2 | TGACTCTCCTTCCAGATCCCA | TGCCCACACTAGGCTGACA | |
| Notch3 | TGCCAGAGTTCAGTGGTGG | CACAGGCAAATCGGCCATC | |
| Cyclin D1 | GCGTACCCTGACACCAATCTC | CTCCTCTTCGCACTTCTGCTC | |
| Lef1 | AACGAGTCCGAAATCATCCCA | GCCAGAGTAACTGGAGTAGGA | |
| Pygo2 | AAGGCCGGTCTGCAAATGAA | GTTAGAAGCGACCAGATGATCC | ORF-Expressed region |
| Pygo2 | AGGCAGCCCCTCTACTTAGC | CCAACCCACTTTCTTCCAAA | Endogenous region |

#### Supplementary Table 9. Clinical characteristics for the primary tumor cases

| | |
|---|---|
| **Number of Slides** | 36 |
| **Age** | |
| Medium | 66 |
| Range | |
| ≤50 | 3 |
| 51-60 | 7 |
| 61-70 | 11 |
| >70 | 15 |
| **Gleason sore** | |
| 7 | 12 |
| 9 | 6 |
| 10 | 18 |
| **pT-status** | |
| T1b | 18 |
| T2c | 3 |
| T3a | 4 |
| T3b | 8 |
| T4 | 2 |
| n.d | 1 |
| **pN-status** | |
| N1 | 17 |
| n.d | 19 |
| **Angiolymphatic invasion** | |
| Yes | 9 |
| No | 8 |
| n.d | 19 |
| **Perineural invasion** | |
| Yes | 17 |
| No | 0 |
| n.d | 19 |

### Supplemental Figure exemplifying the gating strategies for flow cytometry

CD4 and CD8 T lymphocytes gating strategy

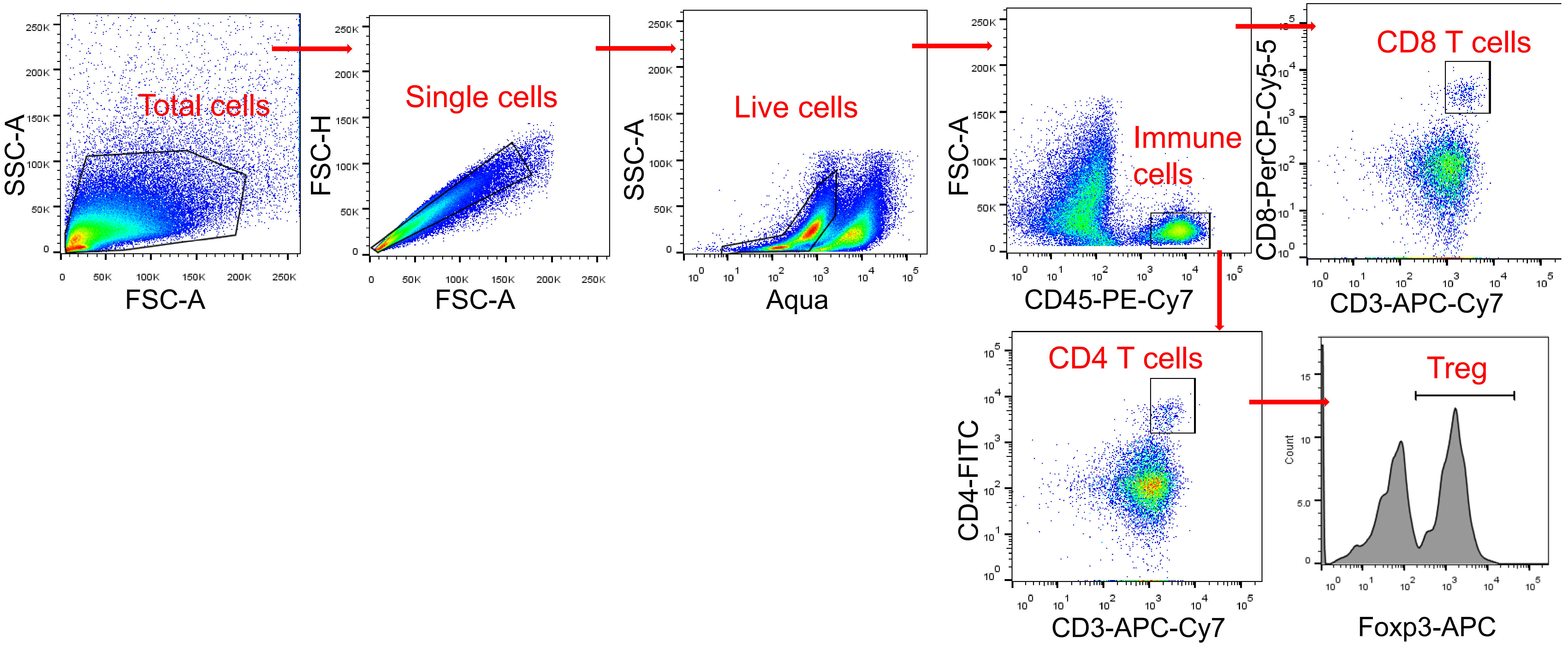


MDSCs and macrophages gating strategy

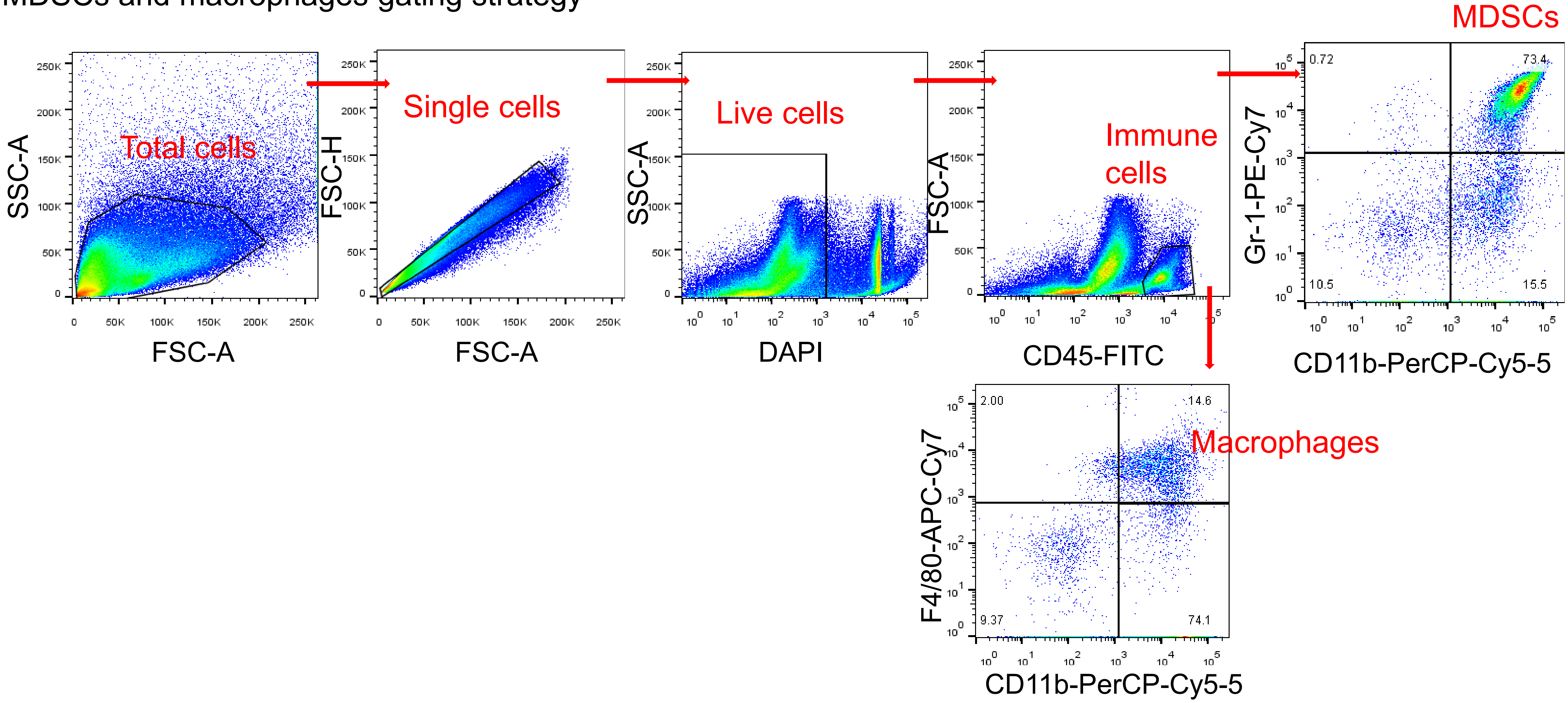
